# Effective mean-field variables explain how biophysical parameters control firing rate and regularity in a spiking network

**DOI:** 10.64898/2026.10.06.757141

**Authors:** Luciano Gonçalves Fernandes

**Affiliations:** Integrative Physiology and Neuroscience Department, College of Veterinary Medicine, Washington State University, Pullman, 99163, WA, USA

**Keywords:** Spiking neural networks, Mean-field theory, Linear stability, Parameter degeneracy, Global sensitivity analysis, Synaptic delay

## Abstract

Parameter sweeps of recurrent spiking networks are often interpreted as evidence that each nominal parameter controls activity through a distinct mechanism. We tested this interpretation in a 10,000 neuron Brunel type network using eight one factor sweeps, local sensitivity analysis, and a global Sobol design with 51,200 simulations. Under the standard calibration, seven of eight nominal parameters influenced the stationary state through effective quantities describing drive and recurrent coupling relative to threshold, input noise, and temporal scale ratios. Without fitting to simulation output, stationary Siegert theory predicted firing rates across the sweeps with a median error of 1%, while a renewal based approximation reproduced spike train regularity with ***R***^***2***^ **= 0.99** for the log coefficient of variation of interspike intervals. Applied without refitting to 10,240 parameter combinations, the theory explained 95% of firing rate variance and 90% of log regularity variance and reproduced the global sensitivity structure. Most apparent effects on rate and regularity therefore reflected the calibration. Synaptic delay contributed little to firing rate variance but was the leading total order factor for synchrony and bursting. Linear stability analysis predicted synchrony onset at 6.0 ms, within the observed transition from 5 to 9 ms, and reproduced oscillation frequencies. In a pre specified generality test, the theory also predicted firing rates in a fluctuation driven regime and reversals of threshold and synaptic time constant effects under alternative calibrations. These results support interpreting parameter dependence through effective drive, coupling, noise, and collective timing rather than nominal parameters alone.

## 1 Introduction

Cortical neurons fire irregularly in vivo, and their excitatory and inhibitory inputs are tightly coupled (Haider et al., 2006; Isaacson & Scanziani, 2011; Renart et al., 2010; Shadlen & Newsome, 1998). Recurrent networks of excitatory and inhibitory spiking neurons are the standard models of this regime (Ahmadian & Miller, 2021; van Vreeswijk & Sompolinsky, 1996, 1998). Depending on their parameters, these networks settle into qualitatively different collective states, which include regular or irregular asynchronous firing, fast population oscillations, bursting, and silence (Brunel, 2000; Brunel & Hakim, 1999; Brunel & Wang, 2003; Ostojic, 2014).

Because these networks can be simulated efficiently with standard tools (Brette et al., 2007; Gewaltig & Diesmann, 2007), they are often studied by sweeping one or two parameters at a time (Brunel, 2000; Kumar, Schrader, Aertsen, & Rotter, 2008; Ostojic, 2014) or by varying many parameters jointly in a global sensitivity design (Saltelli et al., 2008; Tennøe, Halnes, & Einevoll, 2018). These studies report how the firing rate, the regularity of firing, and the synchrony of the population change with each parameter (Brunel, 2000; Kumar et al., 2008; Tennøe et al., 2018). Read one parameter at a time, the sweeps analyzed here would suggest that denser connectivity suppresses activity, that stronger inhibition makes firing irregular (Brunel, 2000; Kumar et al., 2008), that a larger membrane capacitance silences the network, and that a higher threshold changes excitability (Section 3.2). Interpretations of this kind support the view that particular biophysical properties (Brunel & Wang, 2003; Isaacson & Scanziani, 2011), or particular features of graph structure (Haeusler, Schuch, & Maass, 2009; Sporns & Kötter, 2004), control cortical dynamics. Graph descriptors in particular are widely used to relate structure to activity (Bassett & Sporns, 2017; Bullmore & Sporns, 2009; Rubinov & Sporns, 2010). Related computational studies have classified network regimes from the topology of spike trains (Bardin et al., 2019), predicted the rates of single neurons from graph centrality (Fletcher & Wennekers, 2018), and related inhibition and dynamical regime to the computational properties of Brunel networks (Rosati et al., 2026). This reading is incomplete because, in the diffusion approximation, the stationary state of a population of leaky integrate and fire (LIF) neurons depends on the mean and variance of its input and on a few ratios of time constants, not on the nominal parameters themselves (Amit & Brunel, 1997; Brunel, 2000; Fourcaud & Brunel, 2002; Siegert, 1951). Different nominal parameters may therefore act through the same effective quantities, and in that case they are aliases of one another rather than independent controls (Section 2.2). Whether this happens depends on how the model is calibrated (Sections 2.1 and 3.9).

In the widely used NEST-Simulator implementation of the Brunel network (Brunel, 2000; Gewaltig & Diesmann, 2007), synaptic weights are normalized to a target postsynaptic potential, and the external Poisson rate is computed from the threshold. These conventions fix the external drive relative to threshold and tie the membrane and synaptic time constants to the effective synaptic strength (Section 2.2). In the same way, increasing connectivity in a random network with fixed indegree and fixed weights changes the realized graph and the total recurrent input together, so graph descriptors that covary with mean degree need not carry information beyond the recurrent gain (Hu et al., 2013; Kriener et al., 2008; Pernice et al., 2011; Roxin, 2011; Trousdale et al., 2012; Zhao et al., 2011). Structured connectivity can still change network dynamics when it is manipulated independently of coupling strength (Litwin-Kumar & Doiron, 2012).

We therefore evaluate parameter studies of this model family against two predictions that require no fitting (Sections 2.2 and 2.3). For each apparent effect of a parameter, we first ask whether stationary mean field theory predicts it from the effective variables that the calibration fixes (Amit & Brunel, 1997; Brunel, 2000; Siegert, 1951). If it does not, we ask whether the departure is predicted by the linear stability of the stationary state (Brunel, 2000; Brunel & Hakim, 1999). Effects that pass the first test are consequences of the calibration, however mechanistic they look on a parameter axis (Section 4.2). Effects that fail the first test but pass the second are the classical collective instabilities of the model (Brunel, 2000; Brunel & Hakim, 1999), and only effects that fail both would call for a new mechanism. The evaluation also gives a concrete form to parameter degeneracy, the observation that similar activity arises from very different cellular and synaptic parameters (Marder & Goaillard, 2006; Marder & Taylor, 2011; Prinz et al., 2004). Whenever several nominal parameters reduce to one effective variable, degeneracy follows directly from the input statistics seen by single neurons (Section 4.2).

We applied this scrutiny to a Brunel network of 10,000 neurons simulated in NEST-Simulator (Brunel, 2000; Gewaltig & Diesmann, 2007), which is described in Section 2.1. Eight sweeps varied one parameter each, namely the connectivity scaling *ϵ*, the relative inhibitory strength *γ*, the relative external drive *η*, the synaptic delay *δ*, the synaptic and membrane time constants *τ*_*s*_ and *τ*_*m*_, the membrane capacitance *C*_*m*_, and the threshold *θ* (Section 2.4). A local sensitivity design perturbed each parameter by *±*10% with matched random seeds (Section 2.7), and a global Sobol design (Saltelli et al., 2010; Sobol’, 1993) varied all eight parameters jointly in 51,200 simulations (Section 2.8). The spike trains of all simulations were classified into dynamical states (Section 2.5). For every condition, we derived the effective variables from the stored calibration (Section 2.2) and compared the simulation with the stationary Siegert theory (Amit & Brunel, 1997; Brunel, 2000; Siegert, 1951), using a simple renewal approximation of the recurrent input variance to predict regularity (Dummer et al., 2014). We then analyzed the linear stability of each stationary solution (Brunel, 2000; Brunel & Hakim, 1999; Richardson, 2007), including the synaptic filter and the delay of the implementation (Section 2.3). The graph structure of the connectivity sweep was analyzed as an inferential control (Section 2.6).

The effective variables predicted the rate and the regularity of firing for seven of the eight parameters along the sweeps (Section 3.2) and, out of sample, across the global design (Sections 3.4 and 3.7). The same theory reproduced the hierarchy of local sensitivities (Section 3.3) and the ranking of the global Sobol indices (Section 3.6). In a test whose predictions were recorded before the simulations were run, it also predicted the rates at an operating point driven by input fluctuations and under two alternative calibration conventions, including the reversal of the effects of threshold and synaptic time constant that these conventions produce (Section 3.9). The remaining departures were synchronous and bursting states (Section 3.4). The linear stability of the stationary state predicted their onset, their frequency, and their dependence on delay, within a bracket set by two approximations of the recurrent input noise (Section 3.8). Most of the apparent parameter dependence of this model is therefore fixed by its calibration, and most of what remains is the classical collective instability of the asynchronous state (Brunel, 2000; Brunel & Hakim, 1999).

## 2 Methods

### 2.1 Network model and calibration

We used the version of the sparse network of excitatory and inhibitory neurons of (Brunel, 2000) in which synaptic inputs are currents shaped as alpha functions, as implemented in NEST-Simulator 3.10.0 (Gewaltig & Diesmann, 2007) (Table 1; Fig. 1). The network contained *N*_*E*_ = 8000 excitatory and *N*_*I*_ = 2000 inhibitory leaky integrate and fire neurons, in which each synaptic event generates a current with the time course of an alpha function (Gewaltig & Diesmann, 2007; Rotter & Diesmann, 1999). The neurons had membrane time constant *τ*_*m*_, synaptic time constant *τ*_*s*_, capacitance *C*_*m*_, threshold *θ*, reset and leak potentials of 0 mV, and an absolute refractory period *t*_ref_ of 2 ms. The reference values were *ϵ* = 0.1, *γ* = 5, *η* = 2, *δ* = 1.5 ms, *τ*_*s*_ = 0.5 ms, *τ*_*m*_ = 20 ms, *C*_*m*_ = 250 pF, and *θ* = 20 mV. Each neuron received exactly *C*_*E*_ = int(*ϵN*_*E*_) excitatory and *C*_*I*_ = int(*ϵN*_*I*_) inhibitory inputs drawn with replacement (NEST-Simulator fixed indegree rule; autapses and multapses permitted), so *ϵ* is a connectivity scaling parameter rather than a pairwise connection probability. Excitatory and inhibitory neurons received identical indegrees, weights, and external drive.

**Table 1.** Summary of the network model, after Brunel (2000). Synaptic current weights and the external Poisson rate are calibrated as in Eqs. (1)–(3)

|  |  |
| --- | --- |
| **A. Model summary** |  |
| Population | 10,000 neurons: 8,000 excitatory and 2,000 inhibitory |
| Connections | Random fixed-indegree connectivity: $C_E = \text{int}(\epsilon N_E)$, $C_I = \text{int}(\epsilon N_I)$; autapses and multapses are permitted by the connection configuration |
| Neuron model | Leaky integrate and fire; threshold 20 mV, reset and leak potentials 0 mV, refractory period 2.0 ms |
| Synapse model | Postsynaptic current with the time course of an alpha function |
| **B. Connections** |  |
| EE | $E \rightarrow E$, random convergent, excitatory current weight $J_{\text{ex}}$ and delay $\delta$ |
| II | $I \rightarrow I$, random convergent, inhibitory current weight $-\gamma J_{\text{ex}}$ and delay $\delta$ |
| EI | $E \rightarrow I$, random convergent, excitatory current weight $J_{\text{ex}}$ and delay $\delta$ |
| IE | $I \rightarrow E$, random convergent, inhibitory current weight $-\gamma J_{\text{ex}}$ and delay $\delta$ |
| **C. Models** |  |
| Membrane potential ($V_m$) | $\tau_m dV_m/dt = -V_m + RI(t)$ outside the refractory period, where $I(t)$ is the sum of the synaptic currents |
| Action potential | If $V_m(t^-) < \theta$ and $V_m(t^+) > \theta$: (1) $t^* = t$; (2) an action potential is emitted at time $t^*$ |
| **D. Neuron parameters** |  |
| Membrane time constant ($\tau_m$) | 20.0 ms |
| Synaptic time constant ($\tau_s$) | 0.5 ms |
| Membrane capacitance ($C_m$) | 250.0 pF |
| Firing threshold ($\theta$) | 20.0 mV (the external rate is rescaled with $\theta$; Eq. 3) |
| Synaptic delay ($\delta$) | 1.5 ms |
| Relative inhibitory synaptic strength ($\gamma$) | 5.0 (dimensionless inhibitory/excitatory current weight ratio); target excitatory PSP amplitude $J = 0.1$ mV; $J_{\text{in}} = -\gamma J_{\text{ex}}$ |
| Relative external drive ($\eta$) | 2.0 (dimensionless; external rate/threshold rate) |
| Connectivity parameter ($\epsilon$) | 0.1 reference value; used to scale fixed indegrees |
| Number of recorded neurons ($N_{\text{reg}}$) | 500 per population (500 excitatory and 500 inhibitory) |
| Number of excitatory neurons ($N_E$) | 8,000 |
| Number of inhibitory neurons ($N_I$) | 2,000 |
| Excitatory indegree per target ($C_E$) | $\text{int}(\epsilon N_E)$ |
| Inhibitory indegree per target ($C_I$) | $\text{int}(\epsilon N_I)$ |
| Total nominal synaptic indegree | $C_E + C_I$ |

**Fig. 1.**
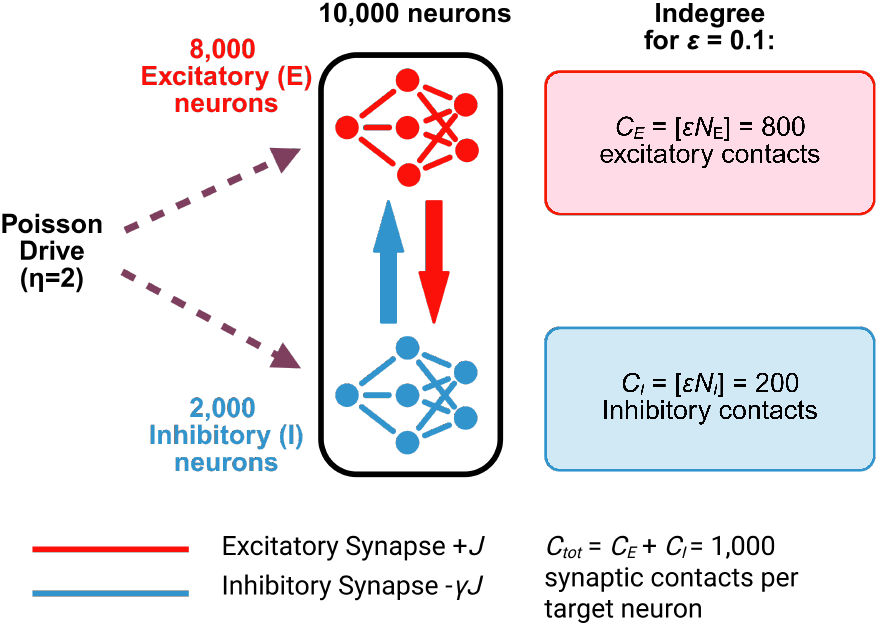
Network schematic. The reference network contains 8,000 excitatory and 2,000 inhibitory leaky integrate-and-fire neurons. Each neuron receives exactly *C*_*E*_ = int(*ϵN*_*E*_) excitatory and *C*_*I*_ = int(*ϵN*_*I*_) inhibitory recurrent inputs and independent Poisson external drive. Excitatory and inhibitory current weights are *J*_ex_ and −*γJ*_ex_, calibrated as described in Section 2.1. Reference parameters are listed in Table 1

The excitatory current weight was chosen so that a single excitatory event produced a peak postsynaptic potential (PSP) of *J* = 0.1 mV,

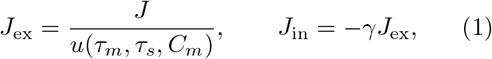

where *u* is the peak PSP produced by a unit-amplitude alpha current,

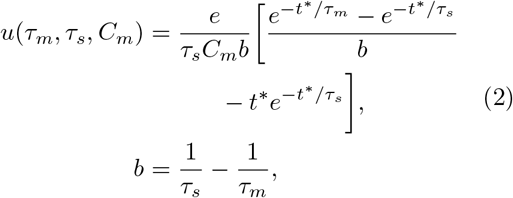

and *t*^∗^ is the time of the PSP peak, obtained from the Lambert-*W* expression. The external input was a Poisson process delivered through the excitatory synapse with rate

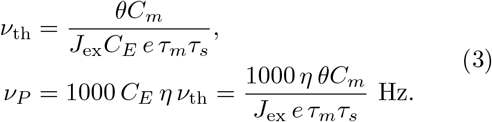

Equations (1)–(3) were re-evaluated for every parameter value in every sweep, with one exception. In the *C*_*m*_ sweeps (one-factor, local, and global), *J*_ex_ and *ν*_*P*_ were calibrated once at *C*_*m*,ref_ = 250 pF and held fixed while the neuronal capacitance was varied, so that a change in *C*_*m*_ was not cancelled by recalibration. All calibration quantities used by each simulation were stored in its metadata and were used directly in the mean-field calculations below.

### 2.2 Effective variables and stationary mean-field theory

#### Effective variables

In the diffusion approximation, an alpha current of weight *J*_*X*_ deposits the charge *J*_*X*_ *eτ*_*s*_ and is equivalent, for the slow membrane dynamics, to a voltage jump *w*_*X*_ = *J*_*X*_ *eτ*_*s*_*/C*_*m*_. Combining this with Eqs. (1)–(3) gives

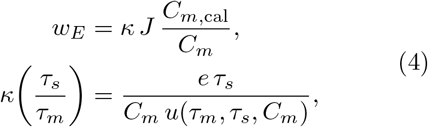

where *C*_*m*,cal_ is the capacitance used in the calibration (*C*_*m*,cal_ = *C*_*m*_ except in the *C*_*m*_ sweeps). The factor *κ*, the ratio between the delta-equivalent jump and the PSP peak, depends only on *τ*_*s*_*/τ*_*m*_; it equals 1.124 at the reference and ranged from 1.010 to 1.809 across the *τ*_*s*_ and *τ*_*m*_ sweeps. With the population rate *ν* expressed in spikes per ms, the mean and variance of the membrane input are

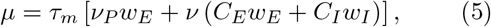

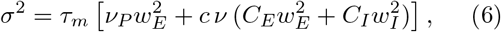

with *w*_*I*_ = −*γw*_*E*_ and *c* defined below. Substituting the calibration, the mean equation takes the dimensionless form

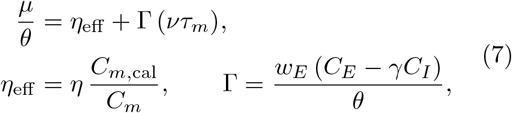

and the external noise satisfies 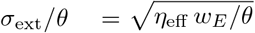. Thus, in this implementation the stationary input statistics depend on the eight nominal parameters only through *η*_eff_ (external drive relative to threshold), Γ (net recurrent coupling relative to threshold), the noise ratios *σ/θ*, and the time-scale ratios *t*_ref_ */τ*_*m*_ and *τ*_*s*_*/τ*_*m*_, with the rate naturally measured as *ντ*_*m*_. The calibration fixes *η*_eff_ = *η* in every sweep except *C*_*m*_. The connectivity scaling *ϵ* and inhibitory strength *γ* both act through *C*_*E*_ − *γC*_*I*_ in Γ; threshold acts through *w*_*E*_*/θ*; and *τ*_*s*_ and *τ*_*m*_ act through *κ*(*τ*_*s*_*/τ*_*m*_), with *τ*_*m*_ additionally setting the time unit. The delay *δ* does not enter the stationary equations at all. At the reference point, *η*_eff_ = 2, Γ = −1.124, and *σ*_ext_*/θ* = 0.106.

#### Stationary rate and regularity

The stationary rate was obtained from the self-consistency condition *ν* = Φ(*µ*(*ν*), *σ*(*ν*)) with the Siegert first-passage formula (Amit & Brunel, 1997; Brunel, 2000; Siegert, 1951)

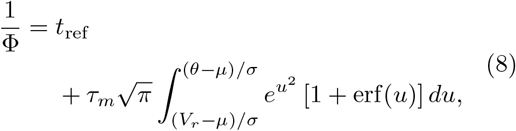

including the colored-noise correction of Fourcaud and Brunel (2002), which shifts threshold and reset by 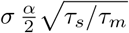 with 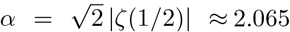. This correction was derived for exponentially filtered synaptic currents in the fast-synapse limit *τ*_*s*_ ≪ *τ*_*m*_; here it is applied to alpha-shaped currents, and 11% of the global design points have *τ*_*s*_*/τ*_*m*_ ≥ 0.2 (13 points exceed 1). Its use outside that asymptotic regime is therefore an empirical approximation, and we report its performance separately for *τ*_*s*_*/τ*_*m*_ *<* 0.2 and with the correction removed. Because excitatory and inhibitory neurons receive statistically identical input, *ν*_*E*_ = *ν*_*I*_ = *ν* holds exactly at this level of description and is not an additional assumption.

The coefficient of variation of the interspike interval (CV_ISI_) was computed from the corresponding first-passage moments (Brunel, 2000),

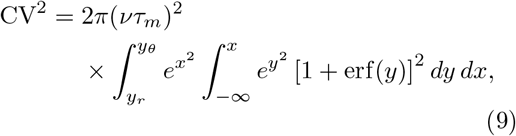

with the same integration limits as Eq. (8). Fixed points were located by scanning *ν* ∈ [0, 1*/t*_ref_] and refined by bisection; roots with a positive slope of the scalar residual, *∂*(Φ − *ν*)*/∂ν >* 0, were excluded as unstable branches of the one-dimensional stationary closure. This criterion concerns only the scalar self-consistency equation and is not a linear stability analysis of the recurrent network. When more than one such root existed (49 of 10,240 global designs, all excitation-dominated), the lowest one was used unless stated otherwise.

#### Renewal-based approximation to the recurrent variance

The classical diffusion approximation treats presynaptic spike trains as Poisson processes (*c* = 1 in Eq. 6). In a mean-driven network this overestimates the input variance seen by the membrane, because the power spectrum of a regular renewal spike train is suppressed below its firing rate and approaches *ν* CV^2^ at low frequency (Dummer et al., 2014). At the reference point the membrane corner frequency, 1*/*(2*πτ*_*m*_) ≈ 8 Hz, lies well below the firing rate of about 32 Hz, so the membrane integrates mainly the suppressed low-frequency part of the recurrent input. We therefore also evaluated a renewal-based low-frequency approximation to the recurrent input variance, in which *c* = CV^2^ was determined self-consistently together with *ν* by damped fixed-point iteration (external Poisson input was left unchanged). Because *ν* CV^2^ is only the zero-frequency limit of the presynaptic spectrum, this approximation is much simpler than the full self-consistent spectral treatment of Dummer et al. (2014) and is judged here by its empirical accuracy. Both versions are reported; the Poisson version is referred to as classical and the approximation as renewal-corrected.

#### Scope of the stationary theory

The stationary theory contains no transmission delay and no dynamics of the population rate. It predicts where the network sits if it is asynchronous, but not whether that asynchronous state is stable (Brunel, 2000; Brunel & Hakim, 1999; Roxin et al., 2005); synchrony and bursting are outside its scope. The stability of each stationary solution was therefore analyzed separately (Section 2.3). All calculations used only the calibration stored with each simulation; no parameter was fitted to simulation output.

### 2.3 Linear stability of the stationary state

To predict where the asynchronous state is lost, we linearized the population dynamics around each stationary solution, following Brunel and Hakim (1999) and Brunel (2000), but with the alpha-shaped synaptic currents and the delay of the implemented network. A small modulation of the population rate, *ν*_1_*e*^*λt*^, changes the mean and variance of the recurrent input after the delay *δ* and the synaptic filter, and the resulting modulation of the output rate closes the loop. Self-sustained modulations exist when

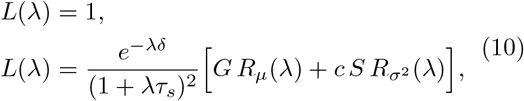

where *G* = *τ*_*m*_(*C*_*E*_*w*_*E*_ + *C*_*I*_ *w*_*I*_) and 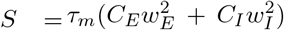 are the recurrent gains of the input mean and variance (Eqs. 5 and 6), (1 + *λτ*_*s*_)^−2^ is the Laplace transform of the unit-area alpha kernel, and *R*_*µ*_ and 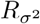 are the rate responses of a single LIF neuron to modulation of the mean and of the variance of its input at the stationary operating point. These responses were computed for complex *λ* by threshold integration of the linearized Fokker–Planck equation (Richardson, 2007), including the refractory period and the same boundary shift as the stationary theory. As *λ* → 0, *L* reduces to the slope of the stationary self-consistency equation, so a real root with *λ >* 0 is the rate instability of the one-dimensional closure; complex roots are oscillatory instabilities. The number of roots with Re *λ >* 0 was counted with the Nyquist criterion, and the fastest-growing root was located by Newton iteration to give the predicted oscillation frequency. The method reproduced the four example states of the original analysis (Brunel, 2000) (Section S3 of Online Resource 1).

Two approximations matter. First, the single-neuron response was computed for white input noise. With filtered synaptic noise the high-frequency response is larger (Brunel et al., 2001; Fourcaud & Brunel, 2002), which would favor fast oscillations. Second, the analysis needs the input variance at the operating point, and the two variance assumptions of the stationary theory differ in an informative way. The classical (Poisson) variance is the one used in the original analysis (Brunel, 2000). The renewal-based variance reproduces the observed regularity but keeps only the zero-frequency limit of the presynaptic spectrum, whereas a regular spike train carries close to Poisson-level power at and above its firing rate, where the oscillatory modes lie. We therefore evaluated stability at both operating points and report both; neither was adjusted to the simulations. Stability was evaluated at every sweep condition, along fine grids of each sweep parameter with the calibration recomputed as in the simulations, and at every global design point with its own delay.

### 2.4 Simulation protocol

Each simulation lasted 6 s at 0.1 ms resolution; the first second was discarded and all measures were computed over the remaining 5 s from 500 recorded excitatory and 500 recorded inhibitory neurons. Subthreshold dynamics were integrated exactly on the time grid (Morrison et al., 2007; Rotter & Diesmann, 1999) (Section S2 of Online Resource 1). Each condition of the one-factor sweeps was simulated with 10 independent seeds, and each realization was treated as one statistical unit. The sweep values are listed in Table 2; when one parameter was varied, all others were held at their reference values. Simulations were run in Python 3.14.4 (**?**) with - Simulator 3.10.0 on a Linux workstation, and the global design on cloud resources with the same code, resolution, duration, and seed conventions. Unless stated otherwise, values are reported as mean *±* standard error of the mean (SEM) across realizations, rounded to three significant figures.

**Table 2.** Variables and values used in the simulations.

| Variable | Description | Value |
| --- | --- | --- |
| $\epsilon$ | Connectivity scaling parameter used to set fixed excitatory and inhibitory indegrees (graph sweep) | 0.001; 0.002; 0.005; 0.01; 0.02; 0.05; 0.1; 0.2; 0.5; 1.0 |
| $N_E$ | Number of excitatory neurons | 8,000 neurons |
| $N_I$ | Number of inhibitory neurons | 2,000 neurons |
| $\gamma$ | Relative inhibitory synaptic strength (inhibitory/excitatory magnitude ratio) | 0; 1; 2; 3; 4; 5; 6; 7; 8; 9; 10 (dimensionless) |
| $\delta$ | Synaptic delay | 0.1; 0.15; 0.25; 0.45; 1.5; 3; 5; 9; 15; 18; 21; 27; 30; 50; 100; 300; 500 ms |
| $\tau_s$ | Synaptic time constant | 0.05; 0.1; 0.25; 0.5; 1.0; 2.5; 5.0 ms |
| $\tau_m$ | Membrane time constant | 2; 4; 10; 20; 40; 100; 200; 400 ms |
| $C_m$ | Membrane capacitance | 10; 25; 50; 100; 125; 250; 500; 1000; 1250; 2500 pF |
| $\eta$ | External Poisson input rate / threshold rate (dimensionless) | 1; 2; 3; 4; 5; 6; 7; 8; 9; 10 (dimensionless) |
| $\theta$ | Firing threshold (external drive rescaled with $\theta$) | 2; 4; 10; 20; 40; 100; 200; 400 mV |

### 2.5 Dynamical measures and state classification

Single-neuron regularity was quantified as the median CV_ISI_ across neurons with at least 10 interspike intervals. Collective synchrony was quantified as the multiscale consensus *S*_cons_, the median of variance-ratio synchrony indices computed with Gaussian kernels of 1, 2, and 5 ms. Spectral concentration was one minus the normalized spectral entropy of the population spike-count spectrum. States were assigned by a prespecified hierarchical classifier: quiescence (Q; mean rate ≤ 0.1 Hz and ≥95% silent neurons) and bursting (B; two-state Gaussian-mixture description of the smoothed population rate with a difference in Bayesian information criterion ΔBIC ≥ 10, rate ratio ≥ 2, and spike concentration in bursts ≥ 0.05) were detected first. Active non-bursting realizations were then classified independently along a regularity axis (regular for median CV_ISI_ ≤ 0.5, irregular for ≥ 0.8, unresolved in between) and a collective axis (asynchronous for *S*_cons_ ≤ 0.05, synchronous for ≥ 0.10, with a 999-surrogate circular-shift test in the grey zone), giving asynchronous regular (AR), synchronous regular (SR), asynchronous irregular (AI), synchronous irregular (SI), or “Other active” when either axis was unresolved. Thresholds were fixed before analysis, and the classifier was validated against synthetic positive controls and real-data anchors. Full definitions are given in Section S1 of Online Resource 1.

### 2.6 Graph characterization of the connectivity sweep

For the *ϵ* series, the realized recurrent connectivity of every simulation was exported and analyzed as a simple directed binary graph (multapses collapsed, autapses removed). Using NetworkX (Hagberg et al., 2008) and custom exact routines, degree statistics, degree-distribution entropy, exact strongly connected components, exact directed eccentricity, diameter and radius, and sampled outward closeness, betweenness, and clustering were computed, and directed three-node motifs were compared with an analytical null reproducing the fixed-indegree generator. To test whether realization-level graph variation predicted dynamics beyond the controlled connectivity gradient, Spearman correlations between four primary graph measures and four primary dynamical measures were computed across all 100 realizations and again after mean-centering both within each *ϵ*, with stratified bootstrap intervals, within-*ϵ* permutation tests, and Benjamini–Hochberg correction. Details are given in Sections S4 and S5 of Online Resource 1.

### 2.7 Local sensitivity analysis

Each of the eight parameters was perturbed by −10% and +10% around the reference (Table 3) using the same five seeds for all 17 conditions (85 simulations). For each seed *r*, parameter *p*, and output *Y*, the normalized centered elasticity was

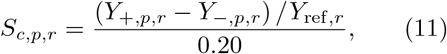

summarized as mean *±* SEM over seeds; one-sided coefficients and response asymmetry were also computed. Mean-field elasticities were obtained by applying the same formula to the mean-field predictions computed from the stored calibration of each of the 17 conditions.

**Table 3.** Parameter values used for the local *±*10% sensitivity analysis.

| Parameter | Reference | -10% | +10% |
| --- | --- | --- | --- |
| $\epsilon$ | 0.10 | 0.09 | 0.11 |
| $\gamma$ | 5.0 | 4.5 | 5.5 |
| $\eta$ | 2.0 | 1.8 | 2.2 |
| Delay (ms) | 1.50 | 1.35 | 1.65 |
| $\tau_s$ (ms) | 0.50 | 0.45 | 0.55 |
| $\tau_m$ (ms) | 20.0 | 18.0 | 22.0 |
| $C_m$ (pF) | 250 | 225 | 275 |
| $\theta$ (mV) | 20.0 | 18.0 | 22.0 |

### 2.8 Global sensitivity analysis

Eight parameters were sampled jointly with a scrambled Sobol sequence (SALib, design generation only; Herman & Usher, 2017): *ϵ* loguniform on [0.01, 0.5], *γ* uniform on [0, 10], *η* uniform on [1, 10], *δ* log-uniform on [0.1, 300] ms, *τ*_*s*_ log-uniform on [0.1, 5] ms, *τ*_*m*_ log-uniform on [4, 200] ms, *C*_*m*_ log-uniform on [50, 1000] pF, and *θ* log-uniform on [4, 200] mV. With base size *N* = 1024, the Saltelli design contained *N* (*D* + 2) = 10,240 points, each simulated with five seeds (51,200 simulations). The design was validated structurally (complete *A, B*, and *AB*_*i*_ blocks, exact parameter agreement) before analysis. First- and total-order indices were computed from design-level mean rates with the Saltelli and Jansen estimators (Saltelli et al., 2010; Sobol’, 1993),

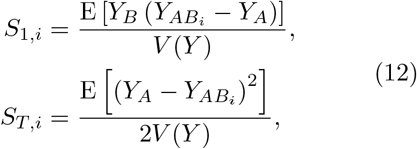

with 95% confidence intervals (CIs) from whole-block bootstrap resampling (10,000 resamples for the original raw-rate analysis, 2,000 for the reanalyses). Negative *S*_1_ estimates were retained.

Three additional analyses of the firing rate were performed. First, indices were recomputed for log(1 + *ν*), which prevents the near-refractory tail of excitation-dominated designs from dominating the variance. Second, the stationary mean-field rate was computed for every design point from its stored calibration and decomposed with the same estimators, which shows how much of the simulated sensitivity structure is predicted by the stationary theory. Third, because the Saltelli blocks cannot be subset without destroying the *A/AB*_*i*_*/B* structure, first-order indices for the inhibition-dominated region *γ* ≥ 4 (equivalently *C*_*E*_ − *γC*_*I*_ ≤ 0) were estimated from the independent *A* and *B* rows with a given-data correlation-ratio estimator, *S*_1_ ≈ Var(E[*Y* | *X*_*i*_ ∈ *b*_*k*_])*/*Var(*Y*) over equiprobable partitions *b*_*k*_ of each input (Saltelli et al., 2008). We used 16 bins, subtracted the finite-bin bias 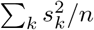 (where 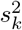 is the within-bin variance), and computed basic bootstrap intervals from 500 resamples. Robustness was checked with 8 and 32 bins, and the estimator was validated by comparing its full-domain values with the canonical Saltelli *S*_1_. Descriptive analyses of silent, high-rate, high-variability, and E/I-discrepant designs and of between-seed variability are reported in Section S11 of Online Resource 1.

#### Dynamical states of the global design

To decompose regularity and synchrony as well as rate, the stored spike trains of the first stochastic realization of every design point (10,240 simulations) were analyzed with the same classifier and settings used for the one-factor sweeps (Section S1 of Online Resource 1), yielding median CV_ISI_, the multiscale synchrony indices, spectral and burst measures, and a state label for each design. The replicate-1 rates agreed with the stored rates (*r* = 0.9997, median absolute difference 0.19 Hz). To measure seed dependence, replicates 2–5 of every design (40,960 further simulations) were classified in the same way; agreement across the five seeds was summarized by the fraction of unanimous designs and Fleiss’ *κ*, and the Sobol indices of the state indicators were recomputed from the five-seed probability of each state. The analyses below use replicate 1 unless stated otherwise. Because CV_ISI_ and synchrony are undefined in some states, three kinds of output were decomposed. State indicators, defined for every design, were analyzed with the full Saltelli estimators: synchronous (SR or SI), bursting (B), non-stationary (SR, SI, or B), and irregular (AI or SI). Two continuous outputs, log_10_CV_ISI_ and *S*_sync_ (the median of the 1, 2, and 5 ms synchrony indices), were defined for all non-quiescent designs. They were analyzed with the Saltelli estimators restricted to complete blocks (835 and 871 of 1,024 blocks, respectively) and with the given-data estimator on the *A* and *B* rows. The renewal-based mean-field CV_ISI_ was computed for every non-synchronous active design (AR, AI, or intermediate regularity; 4,469 designs).

### 2.9 Pre-specified generality test

All results above concern one operating point and one calibration convention. To test whether the reduction depends on either, we simulated 19 further conditions with three seeds each (57 simulations), with the same network, protocol, and classifier. Family A moved the network to a fluctuation-driven operating point (*J* = 0.3 mV, *γ* = 6, *η* = 1.5, other parameters at reference; predicted rate 8 Hz, CV_ISI_ 0.9, and mean input 0.39*θ*) and varied *ϵ* (0.05, 0.2), *γ* (5, 8), *θ* (10, 40 mV), *τ*_*m*_ (10, 40 ms), and *C*_*m*_ (125, 500 pF) one at a time under the standard calibration. Family B kept the reference parameters but changed the calibration convention. In B1, *θ* was varied (10, 15, 30, 40 mV) with the external Poisson rate held at its reference value instead of being rescaled with *θ*. In B2, *τ*_*s*_ was varied (0.25, 0.35, 1.0, 1.5 ms) with the synaptic current weight and the Poisson rate held at their reference values instead of being normalized to the PSP. For every condition the rate, CV_ISI_, linear stability, and state category were predicted from its calibration and recorded with a timestamp before the simulations were run; the analysis recomputed them from the calibration stored with each simulation and obtained identical values. The predicted state was quiescent if the predicted rate was below 0.1 Hz, non-stationary if the classical stationary state was linearly unstable, and otherwise AR, AI, or intermediate according to the renewal-based CV_ISI_ and the classifier thresholds.

## 3 Results

### 3.1 The reference state is mean driven and is predicted by stationary theory

Across 10 realizations, the reference network fired at 32.25*±*0.01 Hz (excitatory) and 32.28*±*0.01 Hz (inhibitory), with median CV_ISI_ = 0.164 *±* 0.0001, low spectral concentration (0.057 *±* 0.002) and low neuron-to-population coherence (0.057 *±* 0.001) (Fig. 2). All realizations were classified as asynchronous regular (AR). The dominant spectral peak was the least reproducible feature (eight realizations peaked at 31–33 Hz and two at 68 and 106 Hz), consistent with weak population rhythmicity.

**Fig. 2.**
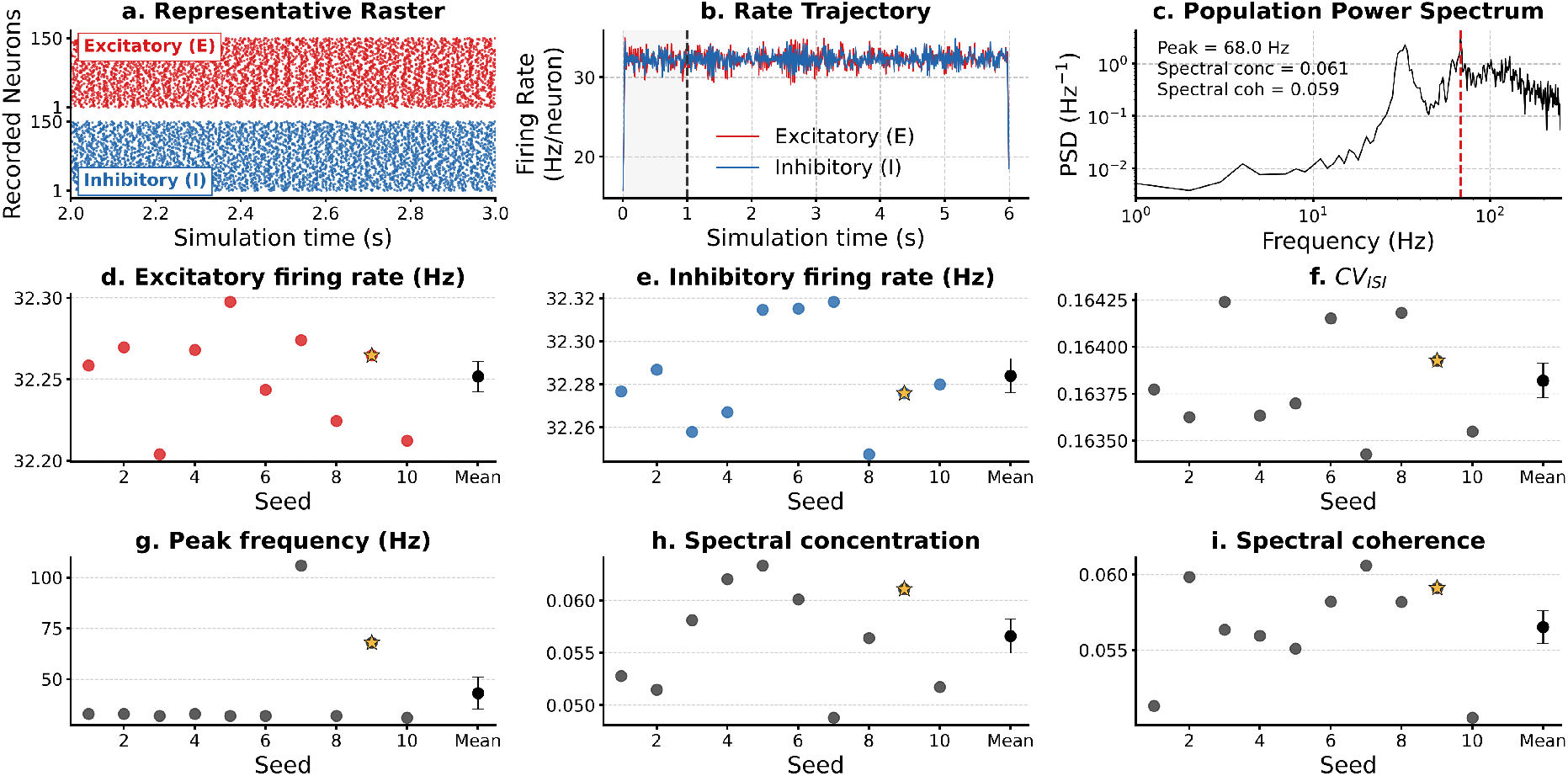
Reference-state dynamics across 10 independent realizations. Reference parameters: *ϵ* = 0.1, *γ* = 5, *η* = 2, *δ* = 1.5 ms, *τ*_*s*_ = 0.5 ms, *τ*_*m*_ = 20 ms, *C*_*m*_ = 250 pF, *θ* = 20 mV. The representative seed (gold star) was chosen objectively as the realization closest to the multivariate centroid of the z-scored reference metrics. **(a)** 1 s raster of 150 excitatory (red) and 150 inhibitory (blue) neurons. **(b)** Population rate trajectories; the first second is burn-in. **(c)** Welch power spectral density of the recorded population; dashed line, dominant frequency. **(d**,**e)** Excitatory and inhibitory rates for each seed. **(f–i)** CV_ISI_, dominant frequency, spectral concentration, and spectral coherence for each seed. In d–i, black symbols show the mean *±* SEM over seeds. Each seed is one statistical replicate

The effective variables of Section 2.2 place this state on the suprathreshold side of the Brunel phase diagram. The calibration delivers a mean external drive *µ*_ext_ = *ηθ* = 40 mV, and net recurrent feedback is inhibitory (Γ = −1.12). The classical Siegert theory predicted a self-consistent rate of 32.6 Hz with a mean input of *µ* = 25.4 mV, above the 20 mV threshold. It overestimated irregularity, however, predicting CV_ISI_ = 0.42, because it assigns the recurrent input a standard deviation of 7.2 mV appropriate for Poisson presynaptic trains. With the recurrent variance scaled by the self-consistent presynaptic CV^2^ (*c* = 0.030), the input standard deviation fell to 2.4 mV and the theory predicted 31.4 Hz and CV_ISI_ = 0.172, within 3% and 5% of the simulation. The reference network is therefore a mean-driven population of noisy oscillators whose irregularity is set mainly by the external Poisson drive, not the fluctuation-driven balanced regime often associated with Brunel networks (Ahmadian & Miller, 2021; Brunel, 2000; Renart et al., 2010; van Vreeswijk & Sompolinsky, 1998). Because the simulations did not measure the cancellation of excitatory and inhibitory currents, we do not describe this state as balanced.

### 3.2 Seven nominal parameters act through a few effective variable

Equation (7) predicts that, at fixed *η*_eff_, sweeps of *ϵ, γ, θ, τ*_*s*_, and *τ*_*m*_ should all fall on a single curve of rate in membrane time units, *ντ*_*m*_, as a function of Γ, up to differences in noise ratio and time-scale ratios. The simulations confirmed this directly, without reference to the mean-field solution (Fig. 3a). The five sweeps spanned Γ from −11.2 to +4.5 and *ν* from 1.8 to 390 Hz, yet traced one monotonic curve. Conditions from different sweeps that share the same (*η*_eff_, Γ) make the point most clearly (Table 4). Increasing connectivity to *ϵ* = 0.2 and increasing inhibition to *γ* = 6 produce the same Γ and gave *ντ*_*m*_ = 0.440 and 0.430. Raising the threshold to 40, 100, or 200 mV produced the same Γ as reducing connectivity to *ϵ* = 0.05, 0.02, or 0.01, and the corresponding rates agreed to within 1% (42.4 versus 42.5 Hz, 53.0 versus 52.6 Hz, and 57.7 versus 57.2 Hz). Similarly, *τ*_*s*_ = 1 ms and *τ*_*m*_ = 10 ms share *τ*_*s*_*/τ*_*m*_ and therefore *κ* and Γ; their rates differed twofold in hertz (30.6 versus 59.4 Hz) but agreed in membrane time units (0.611 versus 0.594).

**Fig. 3.**
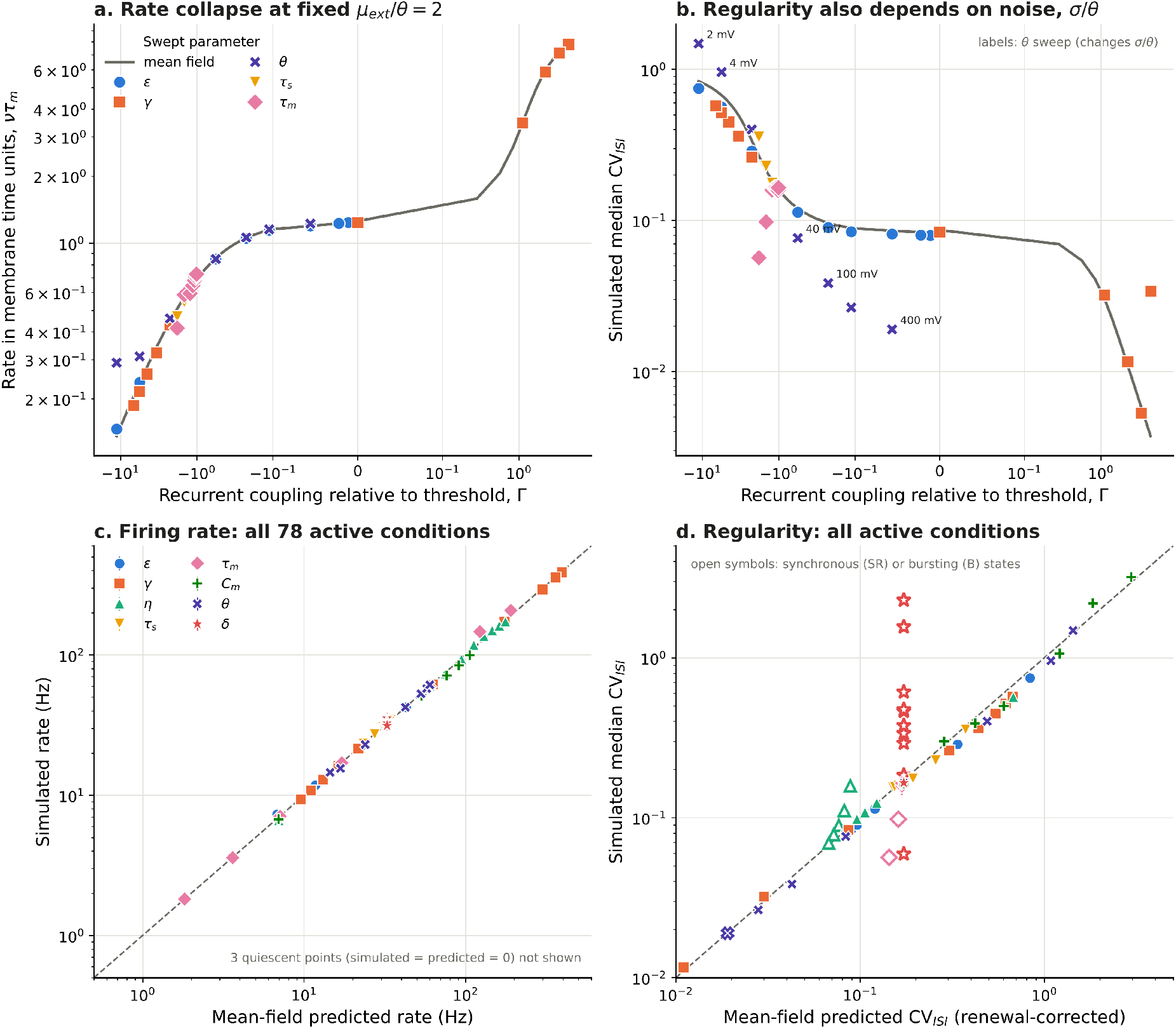
Collapse of the one-factor sweeps onto effective mean-field variables. **(a)** Simulated rate in membrane time units, *ντ*_*m*_, versus net recurrent coupling relative to threshold, Γ, for all sweeps with *η*_eff_ = 2 (*ϵ, γ, θ, τ*_*s*_, *τ*_*m*_). The grey line is the classical mean-field solution along the reference *ϵ* and *γ* family. Symmetric-log abscissa. **(b)** Simulated median CV_ISI_ for the same conditions; the grey line is the renewal-based approximation. Points from the *θ* sweep (labeled) depart from the curve because *θ* also changes the noise ratio *σ/θ*. **(c)** Simulated versus classical mean-field rate for all 78 active conditions of the eight sweeps (mean *±* SD over 10 realizations). **(d)** Simulated versus renewal-based CV_ISI_; open symbols mark conditions classified as synchronous regular or bursting, which account for all large departures. The out-of-sample test on the global design is shown in Fig. 7 and the delay departure in Fig. 6i

**Table 4.**
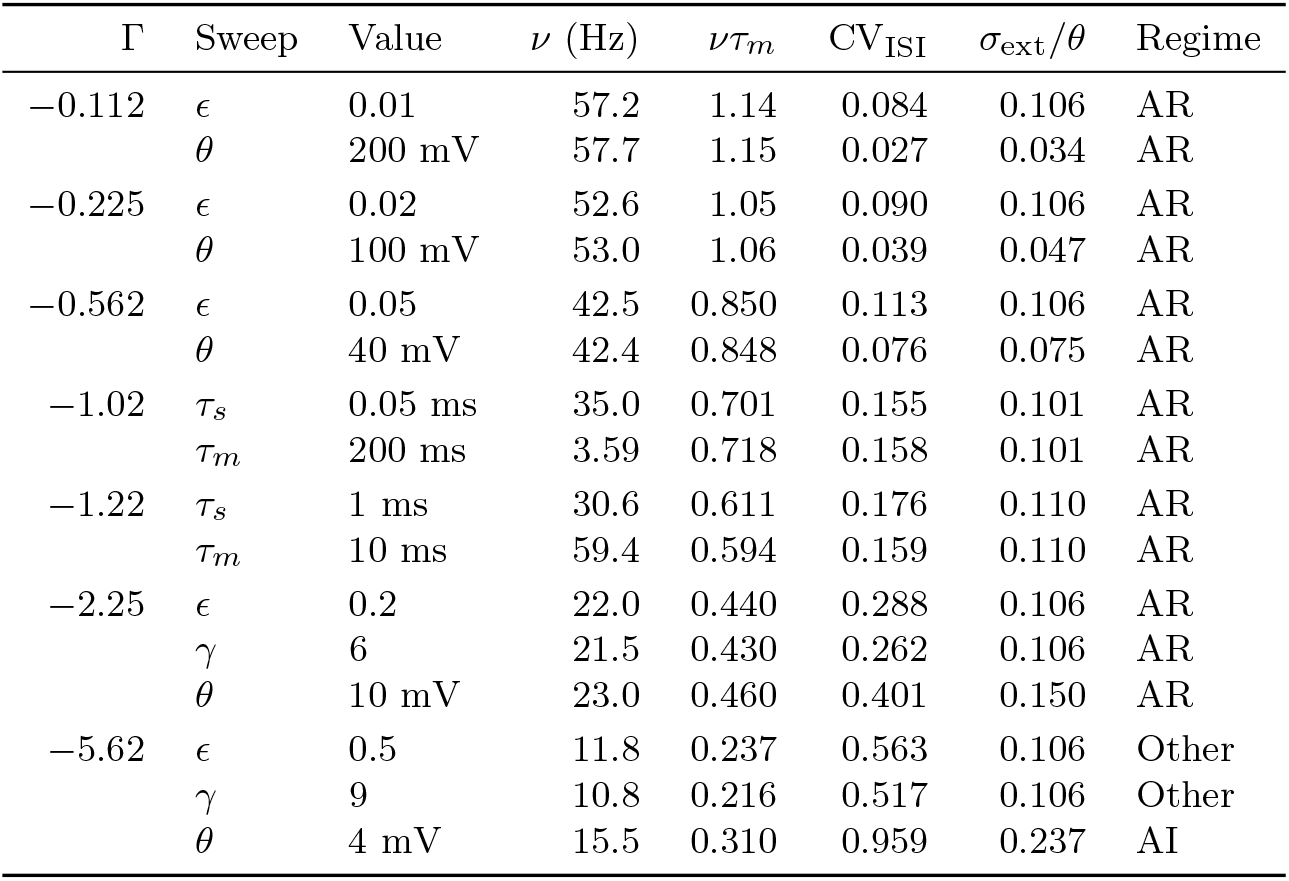
Conditions from different sweeps with the same effective drive and coupling. All conditions have *η*_eff_ = 2. Γ is the net recurrent coupling relative to threshold (Eq. 7); *σ*_ext_*/θ* is the external noise ratio; *ντ*_*m*_ is the simulated rate in membrane time units. Values are means over 10 realizations; regime is the modal classification

The counterintuitive features of several sweeps follow from the same variables. Firing rate increased with threshold because, with *µ*_ext_*/θ* fixed by calibration, raising *θ* only shrinks the fixed 0.1 mV PSP relative to threshold and drives Γ toward zero; the *θ* series is a coupling sweep, not an excitability sweep. Increasing *τ*_*m*_ from 10 to 400 ms reduced the rate from 59.4 to 1.8 Hz, but *ντ*_*m*_ changed only from 0.59 to 0.73 and CV_ISI_ remained between 0.158 and 0.165: above 10 ms, *τ*_*m*_ acted essentially as a change of time unit. In the *C*_*m*_ sweep, the fixed 250 pF calibration gives *η*_eff_ = 2 *×* 250*/C*_*m*_, which crosses 1 (external drive at threshold) at 500 pF and falls to 0.5 at 1000 pF, where the network was silent. At small *C*_*m*_, the same scaling raises the noise ratio (*σ*_ext_*/θ* = 2.65 at 10 pF), explaining why only the *C*_*m*_ and low-*θ* conditions reached the asynchronous irregular state.

Regularity separated sweeps that shared (*η*_eff_, Γ) only through the noise ratio (Fig. 3b; Table 4). For example, at Γ = −0.225, *ϵ* = 0.02 (*σ*_ext_*/θ* = 0.106) gave CV_ISI_ = 0.090, whereas *θ* = 100 mV (*σ*_ext_*/θ* = 0.047) gave 0.039 at the same rate. Conversely, at Γ = −11.2, *θ* = 2 mV (*σ*_ext_*/θ* = 0.335) was irregular (CV_ISI_ = 1.48) and fired at twice the rate of *ϵ* = 1 (CV_ISI_ = 0.75).

The stationary mean-field theory turned these qualitative relations into quantitative predictions for every condition (Fig. 3c,d). Across the 78 active conditions of all eight sweeps, the classical theory predicted the simulated rate with a median absolute error of 0.95%; 88% of conditions were within 5% and 99% within 10% (*R*^2^ = 0.999 on log rate). The three silent *C*_*m*_ conditions were predicted to be silent. The largest rate errors occurred in synchronous states (*τ*_*m*_ = 4 ms, 16%; *τ*_*m*_ = 2 ms, 8%; *δ* = 9 ms, 8%) and in the most strongly fluctuation-driven conditions (low *C*_*m*_ and *θ* = 4 mV, 7–8%). With the renewal-based approximation, the theory predicted CV_ISI_ across 59 non-synchronous active conditions spanning more than two orders of magnitude (0.005 to 3.2) with *R*^2^ = 0.992 on log CV and a median error of 5%. The classical theory could not do so, because it systematically overestimated irregularity in the mean-driven regime. In contrast, for the 19 synchronous or bursting conditions, *R*^2^ fell to 0.16 (Section 3.4).

### 3.3 Mean-field theory predicts the local sensitivity hierarchy

All five reference realizations and all 80 perturbed realizations of the *±*10% design were classified as AR, so the reference lies inside a locally robust region of state space. Quantitative sensitivity nevertheless differed strongly among parameters (Table 5; Fig. 4). The rate elasticities followed the hierarchy *γ* (−2.61) *> η* (1.39) *> C*_*m*_ (−0.943) ≈ *τ*_*m*_ (−0.912) *> ϵ* (−0.480) ≈ *θ* (0.452) ≫ *τ*_*s*_ (− 0.066) *> δ* (0.002), with excitatory and inhibitory elasticities differing by less than 0.005 for every parameter, as expected from the statistical identity of their inputs.

**Table 5.**
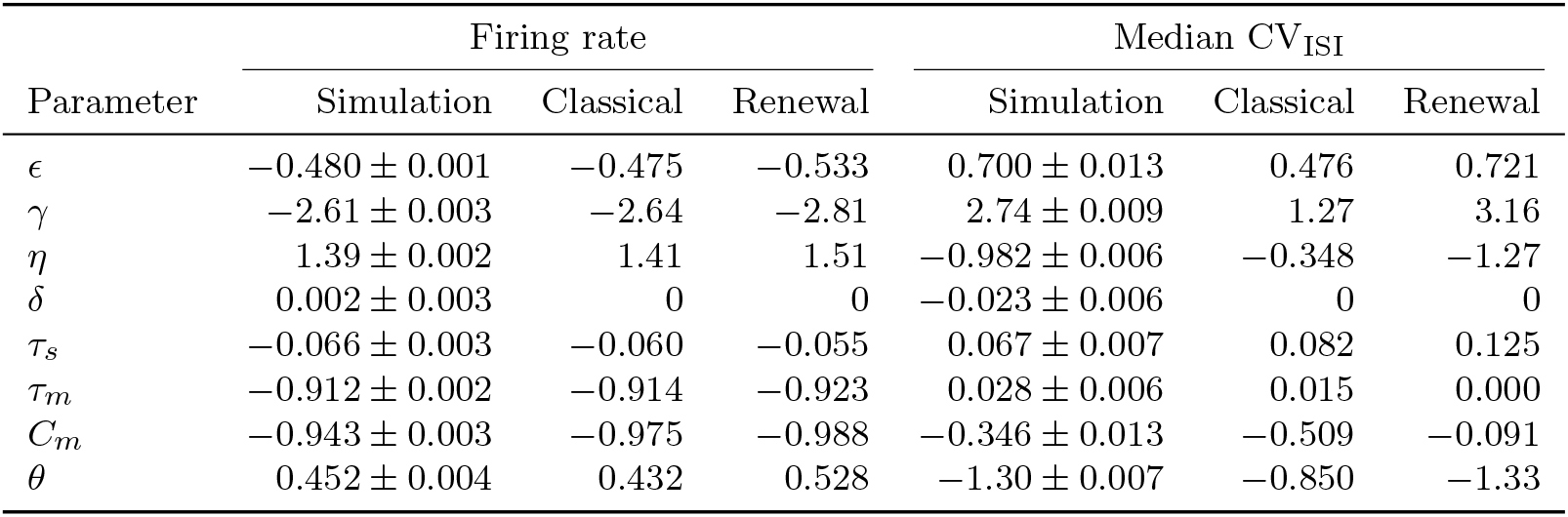
Local *±*10% elasticities: simulation versus stationary mean-field theory. Simulated values are mean *±* SEM over five matched seeds; the rate elasticity is the mean of the excitatory and inhibitory coefficients. Mean-field values were computed from the stored calibration of each condition with the colored-noise correction; “classical” assumes Poisson presynaptic trains and “renewal” scales the recurrent variance by the self-consistent CV^2^

**Fig. 4.**
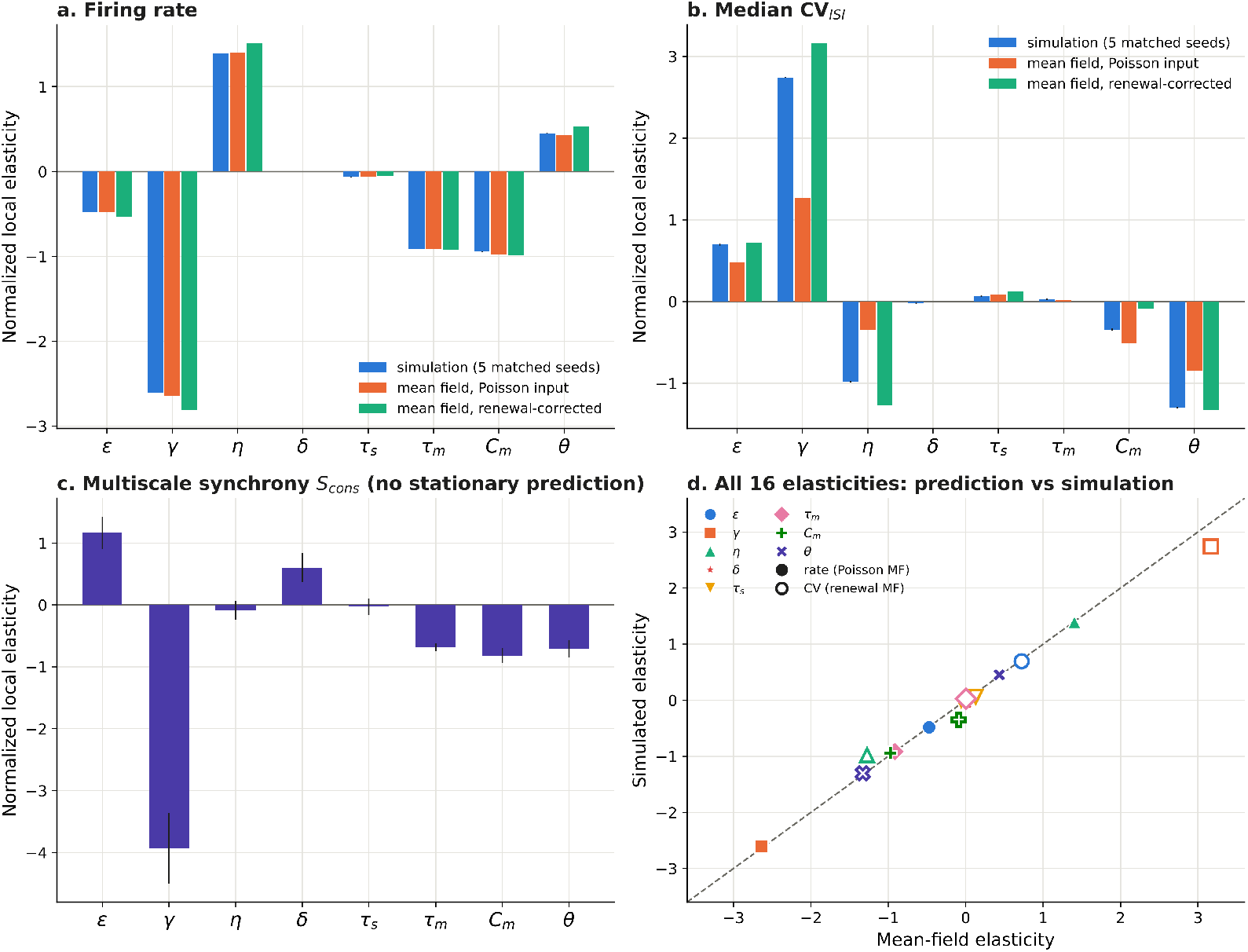
Local sensitivity around the reference state: simulation versus mean-field prediction. Each parameter was perturbed by *±*10% with five matched seeds; elasticities are normalized centered coefficients. **(a)** Firing-rate elasticities from simulation (mean *±* SEM) and from the classical and renewal-corrected mean-field theories. **(b)** Same for median CV_ISI_. **(c)** Elasticities of multiscale synchrony *S*_cons_, which have no stationary counterpart; note the nonzero delay coefficient. **(d)** All 16 rate (filled; classical theory) and CV (open; renewal-corrected theory) elasticities against their predictions; dashed line, identity. All 85 realizations were classified AR. Seed-level elasticities, excitatory/inhibitory comparisons, directional changes, and modified permutation entropy (mPE) elasticities are shown in Fig. S8 in Online Resource 1

The classical mean-field theory, evaluated at the stored calibration of each of the 17 conditions, reproduced this hierarchy and its magnitudes. The largest absolute error was 0.034 (for *γ*, −2.64 predicted versus −2.61 simulated), and every elasticity larger than 0.1 in magnitude was predicted to within 5% (Table 5; Fig. 4a,d). The theory also explains the ordering. *γ* and *ϵ* act through the same quantity Γ ∝ *C*_*E*_ − *γC*_*I*_, but the elasticity of Γ is *γC*_*I*_*/*(*γC*_*I*_ − *C*_*E*_) = 5 with respect to *γ* and 1 with respect to *ϵ*, which accounts for the roughly fivefold ratio of their rate elasticities. *η* is the only parameter that changes the external drive directly. *τ*_*m*_ acts mainly as a time unit, so its elasticity is close to −1. *τ*_*s*_ changes only *κ*(*τ*_*s*_*/τ*_*m*_), which is nearly flat at the reference. Delay has no stationary effect. The renewal-corrected theory predicted the regularity elasticities, including the strong dependence on *γ* (3.16 predicted versus 2.74 simulated) and *θ* (−1.33 versus −1.30), and the near-zero values for *τ*_*m*_ and *δ*. The classical theory underestimated most of these coefficients by a factor of two or more. The one regularity elasticity that neither version captured well was that of *C*_*m*_ (−0.35 simulated, −0.51 classical, −0.09 renewal-corrected), which changes the mean and the variance of the input simultaneously and therefore falls between the two variance assumptions.

Synchrony elasticities (*S*_cons_; Fig. 4c) have no stationary counterpart. They were largest for *γ* (−3.9 *±* 0.6), *ϵ* (1.2 *±* 0.3), and, notably, *δ* (0.60 *±* 0.24), the only parameter with a negligible rate elasticity but a clear synchrony elasticity. This is the first indication that delay acts on a separate dynamical axis.

### 3.4 Departures from stationary theory locate synchronous and delay-driven states

The conditions in which the stationary theory failed were not scattered at random. All 19 conditions with large regularity departures were classified as synchronous regular or bursting (Fig. 3d), and the regime map shows where they occur (Fig. 5). Several such transitions occurred while the rate stayed close to its stationary value: strong external drive (*η* ≥ 6; *S*_cons_ rose from 0.027 at *η* = 5 to 0.361 at *η* = 6 while the rate exceeded its prediction by only 4.5%), short membrane time constants (*τ*_*m*_ = 2 and 4 ms, where *t*_ref_ */τ*_*m*_ is large and the network locked into synchronous regular firing), weak recurrent inhibition (*γ* = 0, close to the refractory ceiling), and very high threshold (*θ* = 400 mV, *S*_cons_ = 0.92). These are the conditions under which the asynchronous stationary state of the Brunel model is expected to lose stability to population oscillations (Brunel, 2000; Brunel & Hakim, 1999), which we test directly in Section 3.8.

**Fig. 5.**
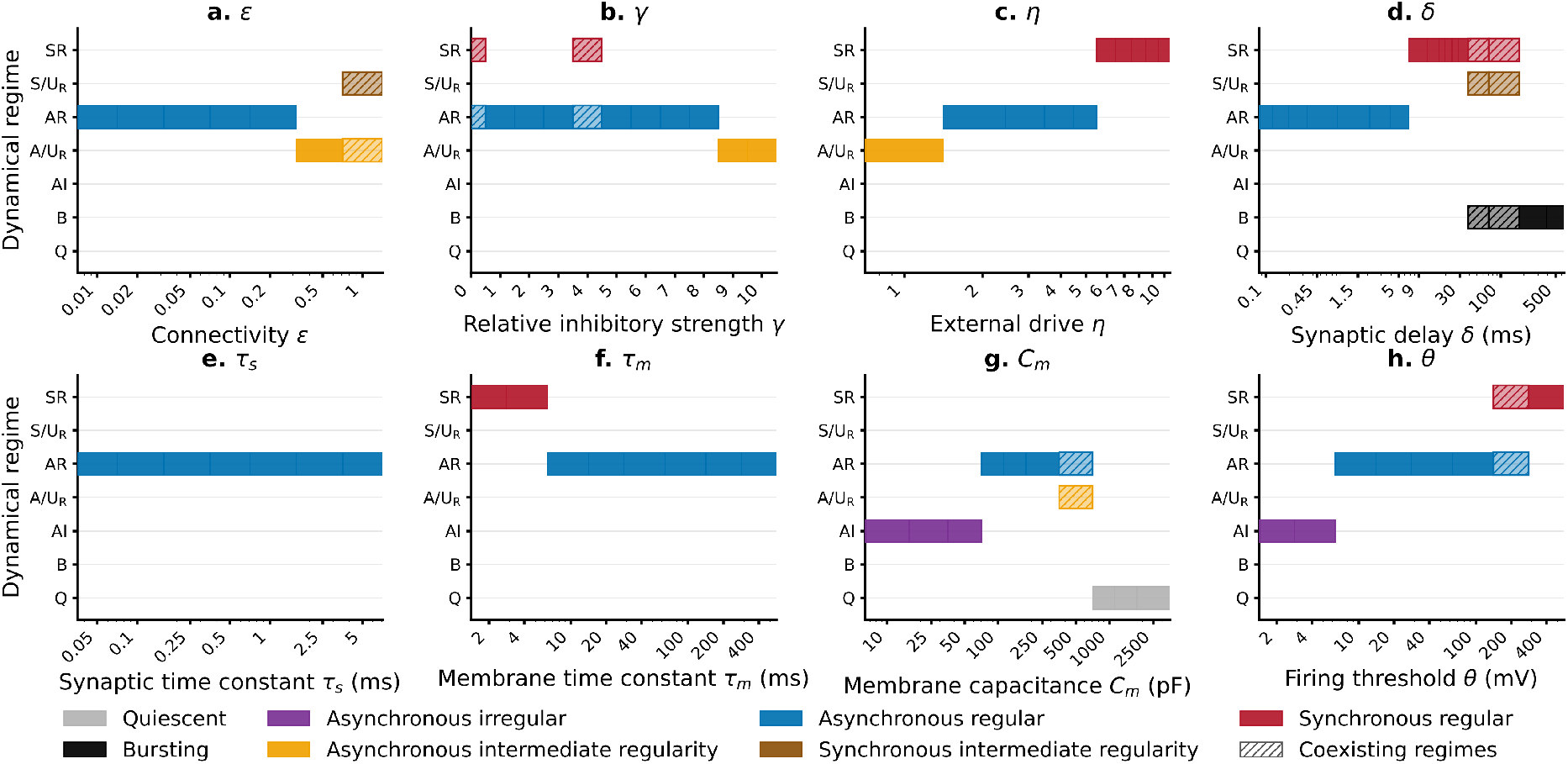
Dynamical regimes across the eight one-factor sweeps. **(a)** *ϵ*, **(b)** *γ*, **(c)** *η*, **(d)** *δ*, **(e)** *τ*_*s*_, **(f)** *τ*_*m*_, **(g)** *C*_*m*_, **(h)** *θ*; 10 realizations per value. States: quiescent (Q), bursting (B), asynchronous irregular (AI), asynchronous with intermediate regularity (*A/U*_R_), asynchronous regular (AR), synchronous with intermediate regularity (*S/U*_R_), and synchronous regular (SR). Hatching marks values at which realizations were split between states. Horizontal extents are bounded by midpoints between tested values (geometric midpoints on logarithmic axes) and are not estimates of transition boundaries. No realization in these sweeps was classified as synchronous irregular; this state does occur elsewhere in parameter space (2.7% of the global design; Section 3.7)

Synaptic delay isolated this second dynamical axis most cleanly (Fig. 6). Delay does not enter any stationary quantity, and the rate indeed remained within 4% of the stationary prediction at every delay up to 500 ms (except 9 ms, +8%). In contrast, *S*_cons_ increased about 100-fold, from 0.0064 *±* 0.0002 at 0.1 ms to 0.658 *±* 0.0004 at 9 ms, spectral coherence rose to 0.94, and all realizations from 9 to 30 ms were synchronous regular. At longer delays the network passed through mixed synchronous and bursting states (50–100 ms) into bursting at 300 and 500 ms (CV_ISI_ = 1.56 *±* 0.03 and 2.29 *±* 0.03, against a stationary prediction of 0.17), still at about 32 Hz. Delay-induced oscillations and bursting are classical properties of recurrent networks (Brunel & Hakim, 1999; Roxin et al., 2005); the point here is quantitative. Within one implementation, delay is the parameter whose effect is invisible to the stationary rate theory, to the local rate elasticities, and to the global rate Sobol indices (below), yet it controls the collective state over the widest range. The synaptic time constant, in contrast, changed rate and regularity exactly as the stationary theory predicted through *κ*, and all 70 *τ*_*s*_ realizations remained AR.

**Fig. 6.**
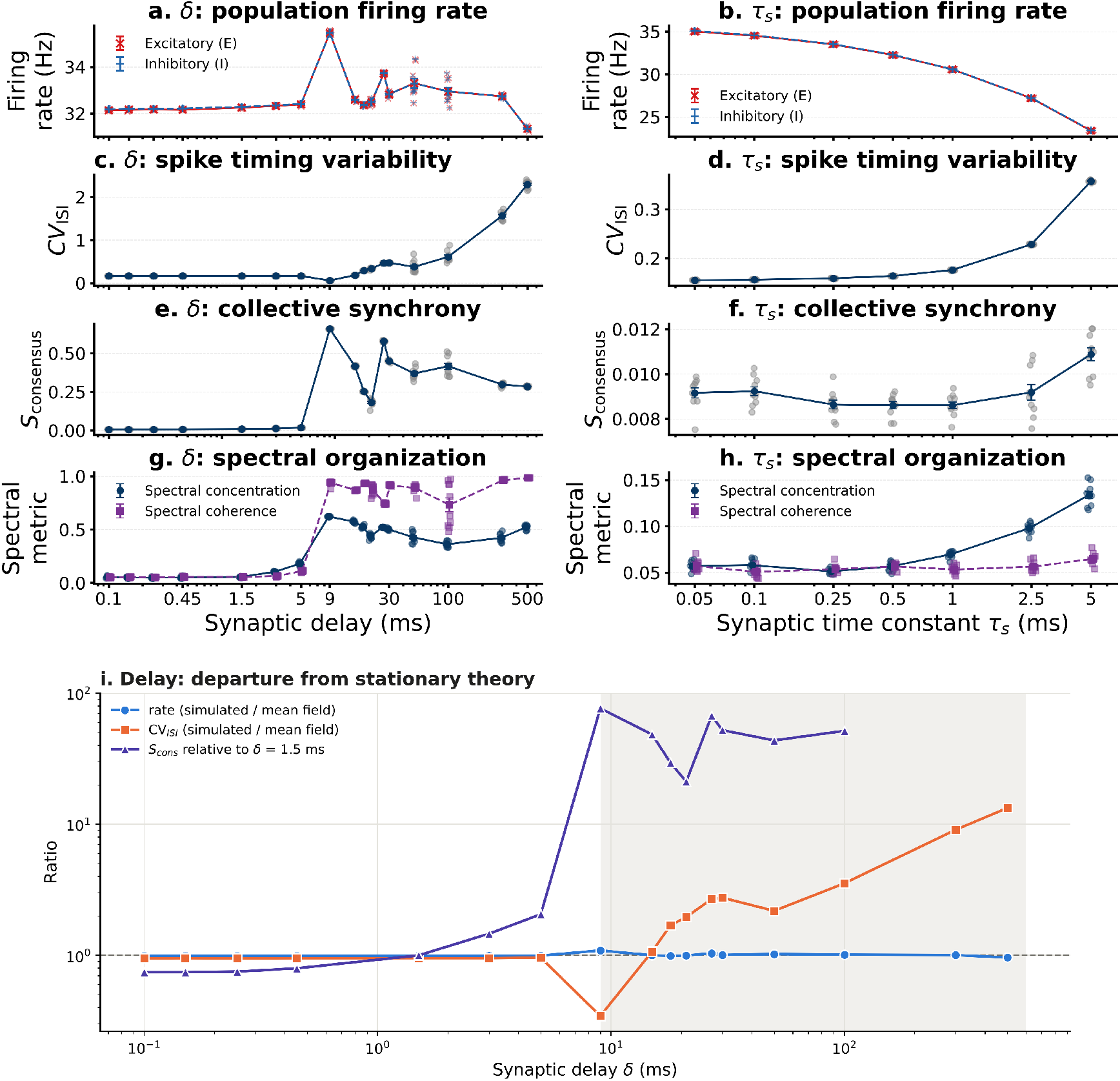
Synaptic delay versus synaptic time constant. Left column, delay *δ*; right column, *τ*_*s*_. **(a**,**b)** Excitatory (red) and inhibitory (blue) rates. **(c**,**d)** Median CV_ISI_. **(e**,**f)** *S*_cons_. **(g**,**h)** Spectral concentration and coherence. Points are realizations; connected symbols are mean *±* SEM; logarithmic parameter axes. Delay reorganized synchrony, regularity, and bursting while leaving the rate essentially unchanged; *τ*_*s*_ changed rate and regularity as predicted by *κ*(*τ*_*s*_*/τ*_*m*_) and never left the AR state. For bursting realizations, *S*_cons_ was reconstructed as the median of the stored 1, 2, and 5 ms synchrony indices. **(i)** Ratio of simulated to stationary mean-field rate and CV_ISI_ across delays, and *S*_cons_ relative to *δ* = 1.5 ms. The rate remains on the stationary prediction at every delay, whereas regularity and synchrony depart from it at *δ* ≥ 9 ms (shaded); *S*_cons_ is not defined for bursting realizations at 300 and 500 ms. Representative rasters are shown in Fig. S5 in Online Resource 1

The global design confirmed this partition out of sample (Fig. 7). Across the 10,240 random parameter combinations, none of which was used to build or tune the theory, the stationary prediction explained 94.6% of the simulated rate variance. Designs with *δ <* 10 ms were predicted within max(2 Hz, 10%) in 87% of cases, against 69% for *δ* ≥ 10 ms. Accuracy declined with delay more steeply in the inhibition-dominated region (from 91% for *δ* ≤ 1.5 ms to 60% for *δ >* 100 ms) than in the excitation-dominated region (from 86% to 66%), as expected if the departures reflect the delay-driven oscillatory instability of inhibition-dominated networks (Fig. 7b; Brunel, 2000; Brunel & Hakim, 1999). Overall, 69% of the 1,057 designs with large errors had *δ* ≥ 10 ms, compared with 42% of all designs. Large errors were also enriched for strong recurrent coupling (median recurrent gain −1.26 versus −0.18 mV/Hz in the full design) and low threshold (median 14.7 versus 28.3 mV), as expected for an oscillatory instability of the asynchronous state.

**Fig. 7.**
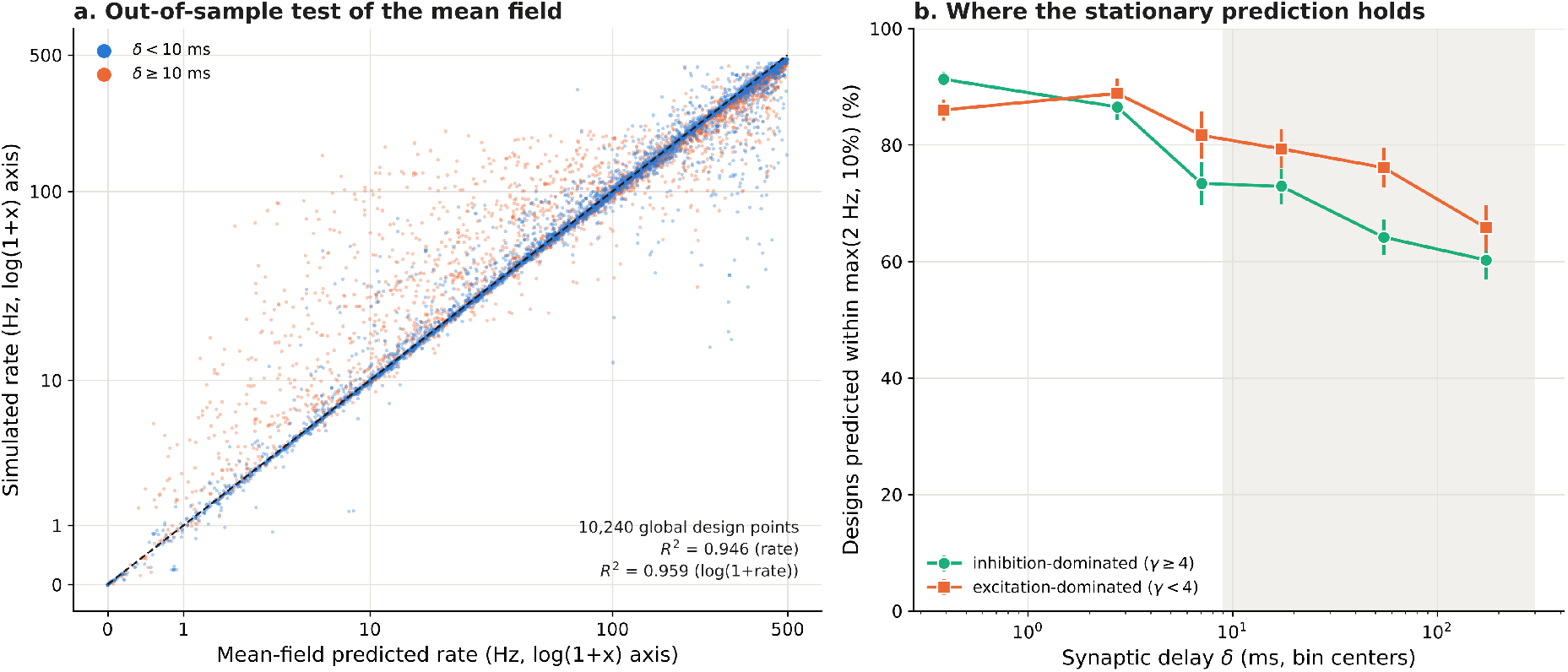
Out-of-sample test of the stationary mean-field theory on the global design. **(a)** Classical mean-field rate, computed from the stored calibration of each design with no fitted parameters, versus the simulated rate (mean of five seeds) for all 10,240 Saltelli design points, colored by synaptic delay; dashed line, identity; log(1 + *x*) axes. **(b)** Percentage of designs whose simulated rate was predicted within max(2 Hz, 10%), by delay bin, separately for inhibition-dominated (*γ* ≥ 4) and excitation-dominated (*γ <* 4) designs (error bars, 95% binomial intervals). Prediction accuracy falls with delay (shaded, *δ* ≥ 9 ms), most steeply when feedback is inhibitory

### 3.5 Connectivity: graph reorganization adds no detectable information beyond recurrent coupling

Because synaptic weights were fixed, increasing *ϵ* from 0.001 to 1 increased both the realized graph and Γ (from 0.011 to 11.2), and the activity changes along the *ϵ* series were reproduced by the stationary theory through |Γ| alone (Fig. 3). The rate fell from 62.0 to 7.3 Hz and CV_ISI_ rose from 0.080 to 0.75. The conditions *ϵ* = 0.2 and *γ* = 6 produced closely matched rates (22.0 and 21.5 Hz) and similar regularity (CV_ISI_ 0.288 and 0.262), although the second manipulation leaves the graph unchanged (Table 4). The binary graph changed as expected for a random fixed-indegree graph of increasing mean degree (Section S4 of Online Resource 1; Figs. S1 and S2 in Online Resource 1). Unique directed edges increased from 9.99 *×* 10^4^ to 6.32 *×* 10^7^; the exact directed diameter fell from 7 to 2; clustering in sampled subgraphs tracked density; and at *ϵ* = 1, 36.8% of synapses were redundant multapses collapsed in the binary graph (Table S2 in Online Resource 1). Directed three-node motif frequencies matched the analytical fixed-indegree null (Jensen–Shannon divergence ≤ 7 *×* 10^−4^), which validates the sampler but is not a biological finding (Fig. S3 in Online Resource 1).

Across the full series, all 16 primary graph– dynamics Spearman correlations were strong (|*ρ*| = 0.92–0.99) and survived false-discovery-rate correction. After centering within each *ϵ*, none survived (largest |*ρ*| = 0.13, minimum *q* = 0.89), so realization-level graph variation did not predict dynamics beyond the controlled gradient. In a fixed-indegree ensemble whose realizations are statistically interchangeable, this null result is expected. It serves as a control against pseudoreplication rather than as evidence that topology is dynamically irrelevant in other ensembles (Hu et al., 2013; Pernice et al., 2011; Roxin, 2011; Zhao et al., 2011).

### 3.6 Global sensitivity: the stationary theory reproduces the ranking, and log-scale analysis revises it

On raw firing rate, the total-order indices ranked *γ* first (*S*_*T*_ = 0.527, 95% CI 0.464–0.592), followed by *C*_*m*_ (0.273), *τ*_*m*_ (0.260), *θ* (0.139), *ϵ* (0.130), *η* (0.096), *τ*_*s*_ (−0.030), and *δ* (−0.020) (Fig. 8a); excitatory and inhibitory indices were indistinguishable. *ϵ* and *θ* had first-order indices consistent with zero (0.003 and 0.018) despite appreciable total-order indices.

**Fig. 8.**
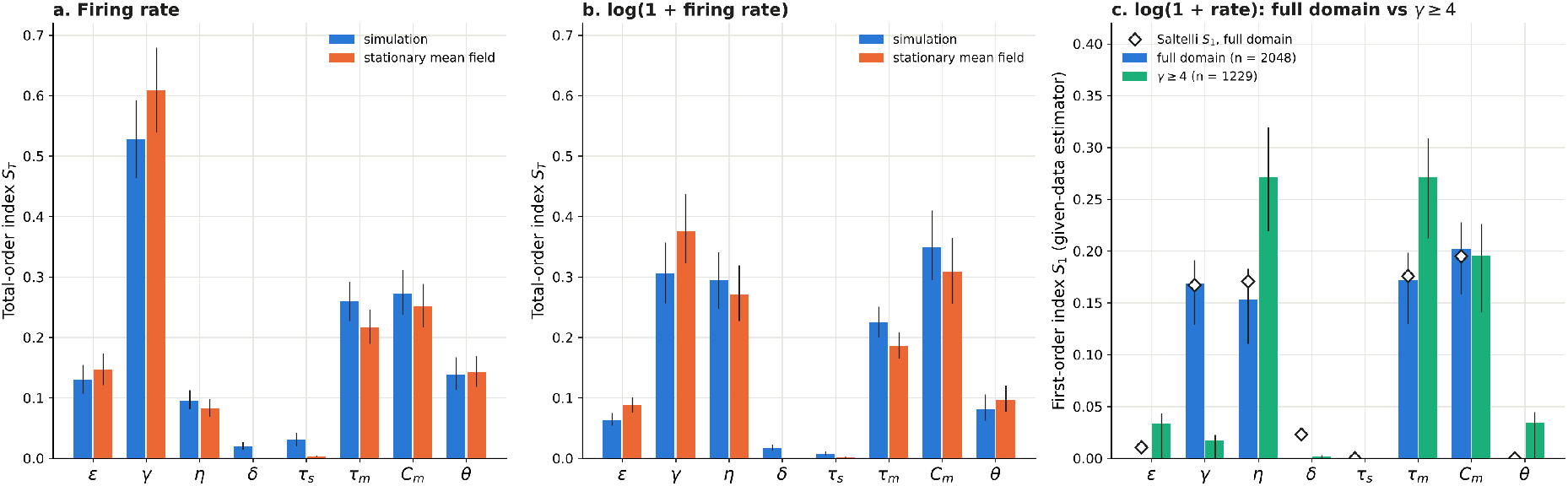
Global Sobol sensitivity: simulation, mean-field prediction, and log-scale reanalysis. 10,240 Saltelli design points, five seeds each (51,200 simulations). **(a)** Total-order indices *S*_*T*_ of the simulated population rate (mean of E and I) and of the stationary mean-field rate computed at the same design points from their stored calibration. **(b)** Same for log(1 + *ν*). Error bars are 95% whole-block bootstrap intervals (2,000 resamples). Delay does not enter the stationary theory, so its mean-field indices are zero by construction. **(c)** First-order indices of simulated log(1 + *ν*) from the given-data correlation-ratio estimator (16 equiprobable bins) applied to the independent *A* and *B* rows, for the full domain (*n* = 2048) and for the inhibition-dominated region *γ* ≥ 4 (*n* = 1229), with 95% basic bootstrap intervals; diamonds show the canonical Saltelli *S*_1_ on the full domain (negative estimates plotted at zero) as validation of the estimator. All indices are conditional on the sampled ranges and output definition. The original raw-rate excitatory and inhibitory indices with *S*_1_, and the response-region summaries, are shown in Fig. S9 in Online Resource 1

The stationary theory, evaluated without fitting at the same 10,240 design points, explained 94.6% of the variance of the simulated design-level rates (95.9% on log(1 + *ν*); Fig. 7a) and reproduced the simulated sensitivity structure (Fig. 8a,b). The agreement did not depend on the colored-noise correction being used outside its fast-synapse domain: restricted to *τ*_*s*_*/τ*_*m*_ *<* 0.2 (9,113 of 10,240 designs) the variance explained was 94.2% (96.1% on log rate), and with the correction removed it was 94.4% (95.6%). Nor did it depend on the branch choice: in all 49 designs with two candidate roots the simulation was silent and the lower root was also the one nearest to the observation, and selecting the upper root instead reduced the variance explained only to 93.1%. On raw rate, the mean-field indices were *γ* 0.608, *C*_*m*_ 0.251, *τ*_*m*_ 0.216, *ϵ* 0.146, *θ* 0.142, *η* 0.083, with near-zero first-order indices for *ϵ* and *θ*, the same signature of non-additivity seen in the simulations. This non-additivity is therefore a property of the effective variables: *ϵ* and *θ* act only through Γ, whose effect depends on the sign of *C*_*E*_ − *γC*_*I*_, which *γ* controls. The simulated indices that the theory could not reproduce were those of delay (*S*_*T*_ = 0.020 versus 0) and *τ*_*s*_ (0.030 versus 0.003), precisely the timing parameters that the theory omits or treats only through *κ*.

Two features of the raw-rate ranking reflect the output scale rather than the network. First, 40% of the sampled domain has *γ <* 4, where net recurrent feedback is excitatory and rates approach the 500 Hz refractory ceiling, so the raw-rate variance is dominated by whether a design lies above or below *γ* = 4. On log(1 + *ν*), the ranking changed to *C*_*m*_ (*S*_*T*_ = 0.349), *γ* (0.306), *η* (0.294), and *τ*_*m*_ (0.224), with *η* tripling in importance; the stationary theory reproduced this reordering (*C*_*m*_ 0.308, *γ* 0.376, *η* 0.272, *τ*_*m*_ 0.186; Fig. 8b). Second, within the inhibition-dominated region (*γ* ≥ 4), *γ* itself contributed almost no first-order variance of log(1 + *ν*) (*S*_1_ = 0.017, 95% CI 0–0.023), whereas *η* (0.271) and *τ*_*m*_ (0.271) dominated, followed by *C*_*m*_ (0.196) (Fig. 8c). This result was insensitive to the partition: with 8, 16, and 32 bins the *γ* index was 0.022, 0.017, and 0.012, and those of *η, τ*_*m*_, and *C*_*m*_ changed by at most 0.011. Applied to the full domain, the same estimator agreed with the canonical Saltelli *S*_1_ to within 0.024 for every parameter (correlation 0.988 on log rate and 0.997 on raw rate; diamonds in Fig. 8c). Within this region, rate is set mainly by the external drive (*η*_eff_ = *η C*_*m*,cal_*/C*_*m*_) and by the time unit *τ*_*m*_, which spans a 50-fold range in the design. Inhibitory strength dominates the global raw-rate ranking because the design straddles the boundary between excitation- and inhibition-dominated feedback, not because it is the main control of activity in the inhibition-dominated regime.

Descriptive analyses of silent (435 designs, 4.25%), high-rate, high-variability, and E/I-discrepant designs, of between-seed variability, and of the near-silent boundary are reported in Section S11 of Online Resource 1.

### 3.7 Dynamical states of the global design: delay controls the collective axis

Classifying the spike trains of every design point turned the global analysis from a rate-only decomposition into one of the dynamical state itself (Fig. 9). Across the 10,240 designs, 33% were asynchronous regular, 5.5% asynchronous irregular, 39% synchronous regular, 2.7% synchronous irregular, 10% bursting, 4.6% quiescent, and 4.8% of intermediate regularity. Synchronous irregular states, absent from the one-factor sweeps, therefore do occur in this network, but only in regions of parameter space that the sweeps did not visit. These labels were reproducible across stochastic realizations. All five seeds of a design received the same label in 95.5% of designs (Fleiss’ *κ* = 0.97) and agreed on whether the design was non-stationary in 96.2% (*κ* = 0.96); replicate 1 matched the modal label of its design in 98.8% of cases, and the state fractions over all 51,200 simulations differed from those of replicate 1 by less than 0.1 percentage point.

**Fig. 9.**
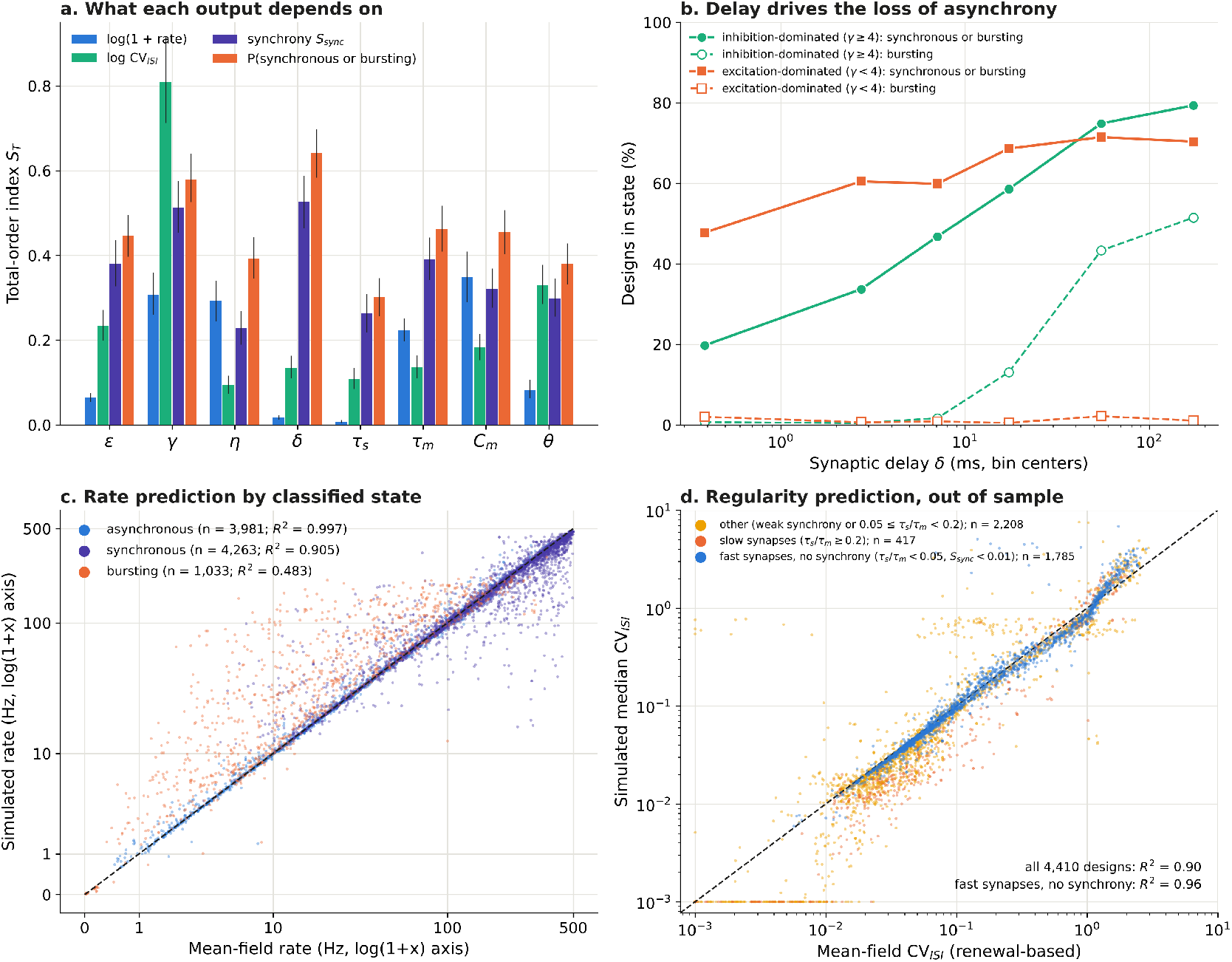
Dynamical states across the global design. The publication classifier was applied to the first realization of each of the 10,240 Saltelli design points. **(a)** Total-order Sobol indices for log rate, log CV_ISI_, multiscale synchrony *S*_sync_, and the probability of a synchronous or bursting state (95% whole-block bootstrap intervals). CV and synchrony use complete blocks only (835 and 871 of 1,024). Delay is negligible for rate but leading for synchrony and state. **(b)** Percentage of designs in synchronous or bursting states (solid) and in bursting states alone (dashed) by delay bin, for inhibition- and excitation-dominated feedback. **(c)** Classical mean-field rate versus simulated rate, by classified state. **(d)** Renewal-based mean-field CV_ISI_ versus simulated median CV_ISI_ for all non-synchronous active designs; dashed lines, identity

The mean-field partition reported above held exactly along these independently assigned labels. For the 3,981 asynchronous designs (AR or AI), the stationary theory predicted the simulated rate with *R*^2^ = 0.997 and a median error of 1%, and 96% of designs fell within max(2 Hz, 10%). For synchronous designs (SR or SI; *n* = 4,263) the corresponding values were *R*^2^ = 0.905 and 75%, and for bursting designs (*n* = 1,033) *R*^2^ = 0.48 and 34% (Fig. 9c). Regularity was the more demanding out-of-sample test (Fig. 9d). Across the 4,410 non-synchronous active designs with a defined CV_ISI_, the renewal-based theory explained 90% of the variance of log CV (classical theory, 61%). It was, however, less accurate than in the sweeps: the median error was 15%, and 58% of designs fell within 20%. (Values below 10^−3^, under the resolution set by the 0.1 ms time grid, were set to 10^−3^; this affected 248 simulated and 87 predicted values.) The errors were systematic, not random.

Where the diffusion approximation is expected to hold, that is with fast synapses (*τ*_*s*_*/τ*_*m*_ *<* 0.05) and no detectable synchrony (*S*_sync_ *<* 0.01; 1,785 designs), the theory explained 96% of the variance, with a median error of 9%, and 77% of designs fell within 20%. Accuracy fell outside that domain:

- with slow synaptic filtering (*τ*_*s*_*/τ*_*m*_ ≥ 0.2; median error 69%), where the simulated neurons fired more regularly than the white-noise first-passage formula predicts;
- with weak synchrony below the classifier’s threshold (*S*_sync_ ≥ 0.01; 30%);
- near refractory saturation (rates above 100 Hz; 38%).

In every subset the renewal-based approximation was much more accurate than the classical theory, whose median errors ranged from 93% to 194%.

The Sobol decomposition separated the two axes sharply (Fig. 9a). Log rate was largely additive (*S*_1_ = 0.72) and controlled by the parameters that set *η*_eff_, Γ, and the time unit, with delay contributing almost nothing (∑*S*_*T*_ = 0.017). Log CV_ISI_ was also largely additive (∑*S*_1_ = 0.67) and dominated by *γ* (*S*_1_ = 0.47, *S*_*T*_ = 0.81), which sets the sign of the recurrent feedback. The collective outputs behaved in the opposite way. Synchrony and the state indicators were strongly non-additive (∑ *S*_1_ = 0.11–0.33), and delay was the leading total-order factor for synchrony (*S*_*T*_ = 0.53, 95% CI 0.46–0.59), for bursting (0.69, 0.59–0.80), and for leaving the asynchronous state (0.64, 0.58–0.70), with *γ* second in each case. Recomputed from the five-seed probability of each state, every total-order index changed by at most 0.03 (delay: 0.63 for leaving the asynchronous state and 0.69 for bursting; Section S10 of Online Resource 1). In the inhibition-dominated region, the fraction of synchronous or bursting designs rose from 20% for *δ* ≤ 1.5 ms to 79% for *δ >* 100 ms, and the fraction of bursting designs from 0.7% to 52% (Fig. 9b). In excitation-dominated designs, synchronous or bursting states were already common at short delays (48%, almost all synchronous regular), consistent with the locking of near-refractory, highly regular neurons. The parameter that is invisible to the rate decomposition is therefore the dominant control of the collective state, and its effect depends strongly on the other parameters, as expected for an instability boundary in a multidimensional space.

### 3.8 Stability of the stationary state predicts where synchrony begins

The departures from the stationary theory are where the asynchronous state is expected to lose stability. We tested this with the linear stability analysis of Section 2.3, evaluated without fitting at every sweep condition, along fine grids of every sweep parameter, and at every global design point. At the reference operating point, the classical loop *L*(*iω*) passed to the left of the critical point for *δ* = 1.5 ms and encircled it for *δ* = 9 ms (Fig. 10a). Along the delay axis, the growth rate of the fastest mode crossed zero at *δ* = 6.0 ms, between the last asynchronous (5 ms) and the first synchronous (9 ms) simulated delays (Fig. 10b). The predicted frequency at onset was 44 Hz and fell with delay, and it matched the population frequency of the synchronous states at 9, 15, 18, and 21 ms (predicted 36, 26, 23, and 20 Hz; observed 35, 30, 25, and 21 Hz). At long delays several oscillatory modes were unstable at once (three at 100 ms and nine at 300 ms), where the simulations turned from synchronous oscillation to bursting, a transition that a linear analysis can signal but not describe.

**Fig. 10.**
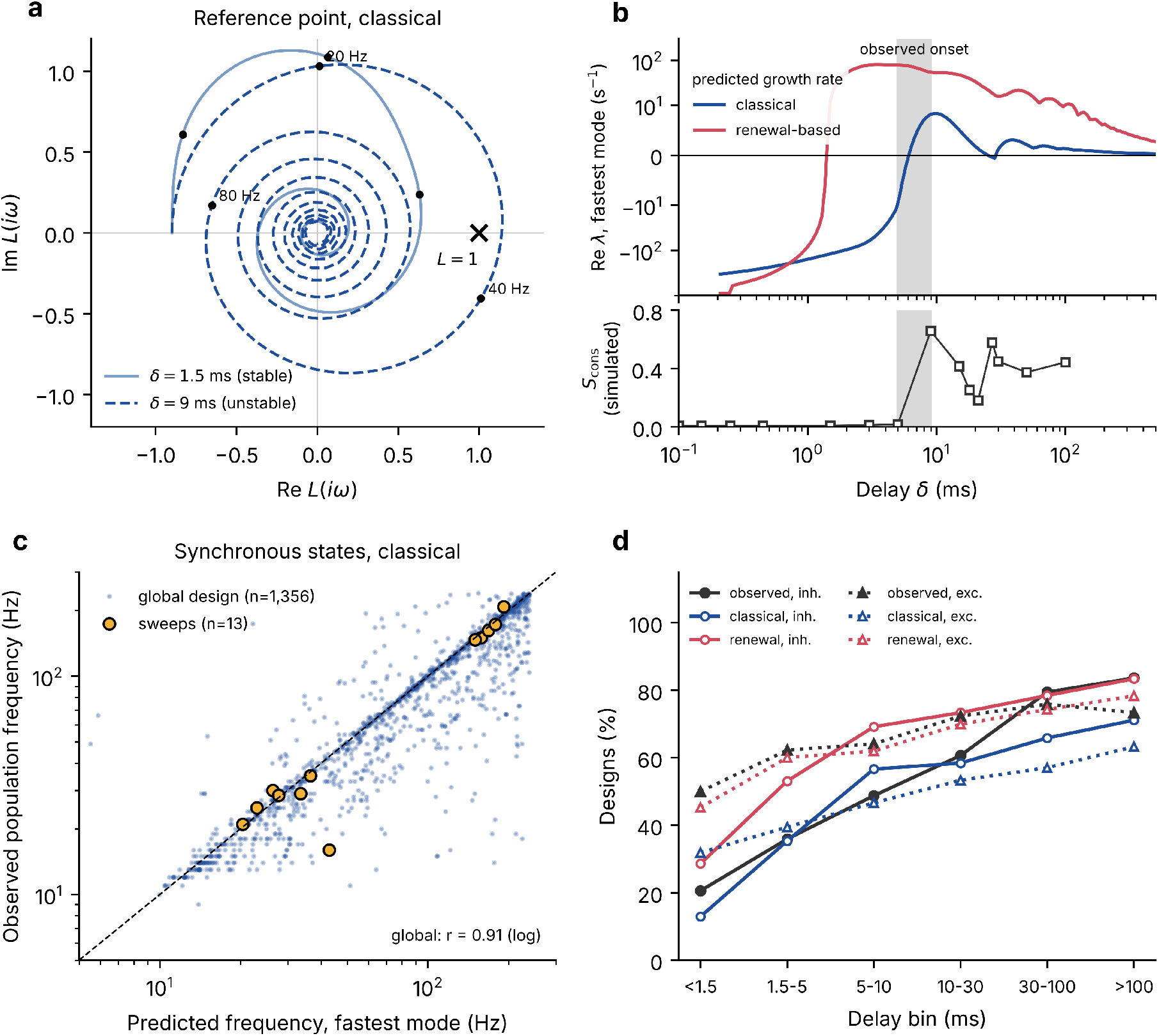
Linear stability of the stationary state predicts the collective axis. **(a)** Nyquist diagram of the loop *L*(*iω*) (Eq. 10) at the reference operating point (classical variance) for *δ* = 1.5 ms, which leaves the critical point *L* = 1 (cross) outside, and for *δ* = 9 ms, which encircles it; dots mark 20, 40, and 80 Hz. **(b)** Top, growth rate of the fastest-growing mode versus delay at the classical (blue) and renewal-based (red) operating points (symmetric-log axis); bottom, simulated *S*_cons_ of the delay sweep (undefined for the bursting states at 300 and 500 ms); grey band, observed onset between 5 and 9 ms. **(c)** Predicted frequency of the fastest-growing mode (classical) against the observed dominant population frequency for synchronous states of the sweeps (orange) and of the global design (blue dots); dashed line, identity. **(d)** Global design: observed percentage of non-stationary (SR, SI, or B) designs and predicted percentage of linearly unstable designs, by delay bin, for inhibition-dominated (*γ* ≥ 4, solid) and excitation-dominated (*γ <* 4, dotted) feedback; quiescent designs excluded

Table 6 compares the predicted and observed boundaries along all sweeps. The classical analysis predicted no instability along the *ϵ, τ*_*s*_, and *C*_*m*_ sweeps, in which every realization was asynchronous or quiescent, and it placed the *δ, τ*_*m*_, and *γ* boundaries inside the observed transition intervals. It placed the *η* boundary too late (7.25 against an observed 5–6) and predicted no instability up to *θ* = 400 mV, where the network was synchronous. The renewal-based operating point erred in the opposite direction: it predicted every observed transition but placed each boundary on the asynchronous side of the observed interval. In every sweep with a transition, the classical boundary therefore fell inside the observed transition interval or on its synchronous side, and the renewal-based boundary on its asynchronous side. Condition by condition, the classical analysis identified 15 of the 19 synchronous or bursting conditions and predicted no instability in any of the 62 asynchronous or quiescent ones; the renewal-based analysis identified all 19 but also predicted instability in 28 asynchronous conditions (Table S3 in Online Resource 1). These false alarms were weak (median loop gain at the phase crossover 1.12, against 3.86 in the synchronous conditions), and in many of them the simulated spectrum had a weak peak close to the predicted frequency (for example 125–136 Hz for *γ* = 6– 10, predicted 120–157 Hz), as expected for weakly damped modes near the boundary. In the synchronous regular states produced by strong drive or short membrane time constants, the predicted frequency tracked the firing rate, as the simulations did (*η* = 8–10: predicted 158–178 Hz, observed 150–173 Hz; *τ*_*m*_ = 4 ms: 150 against 147 Hz).

**Table 6.** Instability boundaries along the one-factor sweeps: simulation versus linear stability of the stationary state. Observed: last asynchronous and first synchronous tested values. Predicted: parameter ranges in which the stationary state is linearly unstable, from fine grids with the calibration recomputed as in the simulations, at the classical and renewal-based operating points. All other parameters at reference

| Sweep | Observed transition | Unstable, classical | Unstable, renewal-based |
| --- | --- | --- | --- |
| $\delta$ (ms) | AR at 5 → SR at 9 | 6.0–25, $> 28.7$ | $> 1.4$ |
| $\eta$ | AR at 5 → SR at 6 | $> 7.25$ | 1.85–2.75, 2.95–3.85, $> 3.95$ |
| $\tau_m$ (ms) | AR at 10 → SR at 4 | $< 5.5$ | $< 22.7$ |
| $\gamma$ | AR at 1 → SR at 0 | $< 0.15$ | $< 1.35$, 2.45–2.85, $> 4.95$ |
| $\theta$ (mV) | AR at 200 → SR at 400 | none ($\leq 400$) | 14.6–22.7, $> 189$ |
| $\epsilon$ | none | none | $> 0.094$ |
| $\tau_s$ (ms) | none | none | $< 0.59$ |
| $C_m$ (pF) | none | none | 79–262 |

The global design tested the same analysis out of sample, with each design’s own delay, synaptic time constant, and calibration (Fig. 10c,d). Among the 9,765 active designs, the classical analysis predicted instability in 69% of non-stationary (SR, SI, or B) designs and stability in 86% of the others; the renewal-based analysis reached 85% and 73%. As in the sweeps, the two operating points bracketed the observed behavior: of the designs predicted stable at both operating points, 19% were non-stationary; of those predicted unstable only with the renewal-based variance, 59%; and of those unstable at both, 86%. Because the state labels were reproducible across seeds, these disagreements reflect the theory rather than label noise. Designs whose seeds disagreed on whether the state was non-stationary were rare in every category but were most common where the two predictions differ (7% of the designs unstable only with the renewal-based variance, against 4–5% of those stable at both operating points and 2% of those unstable at both).

Bursting and synchronous irregular designs were predicted unstable in 91% and 92% of cases by the classical analysis. The analysis also reproduced the dependence on delay that dominated the Sobol decomposition of the dynamical state: in inhibition-dominated designs the observed fraction of non-stationary states rose from 21% at *δ* ≤ 1.5 ms to 84% above 100 ms, against predicted fractions of unstable designs rising from 13% to 71% (classical) and from 29% to 83% (renewal-based) (Fig. 10d). A Sobol decomposition of the predicted instability had the same two leading total-order factors as that of the observed state (largest *S*_*T*_ : classical, *γ* 0.56, *δ* 0.48; renewal-based, *γ* 0.55, *δ* 0.55; observed, *δ* 0.64, *γ* 0.58), and, like the observed state, it was non-additive (∑ *S*_1_ = 0.47 and 0.27, against 0.11 observed). For synchronous designs predicted unstable, the predicted frequency of the fastest mode matched the observed population frequency (log correlation 0.91, median relative difference 6%, *n* = 1, 356 with both frequencies below 240 Hz; Fig. 10c). The main systematic miss was in excitation-dominated designs: 473 non-stationary designs there were predicted stable at both operating points, almost all synchronous regular (95%) and firing close to the refractory ceiling (median 385 Hz), where regular neurons lock to one another in a way the diffusion-based analysis does not capture.

### 3.9 A pre-specified test: a fluctuation-driven regime and other calibration conventions

The theory passed the pre-specified generality test (Fig. 11; Table S4 in Online Resource 1). Across the 18 active conditions, the classical theory predicted the simulated rate with a median error of 4.2%, every condition within 10% (*R*^2^ = 0.996 on log rate), and the predicted silence at *C*_*m*_ = 500 pF was observed (0.002 Hz predicted, 0.003 Hz simulated). In the fluctuation-driven family, where the mean input ranged from −0.43*θ* to 0.75*θ* and CV_ISI_ from 0.53 to 1.15, the rate was overestimated by 3–10% (median 6%), the same direction and size as in the most fluctuation-driven sweep conditions (6.5–8.4% at *C*_*m*_ ≤ 50 pF and *θ* = 4 mV).

**Fig. 11.**
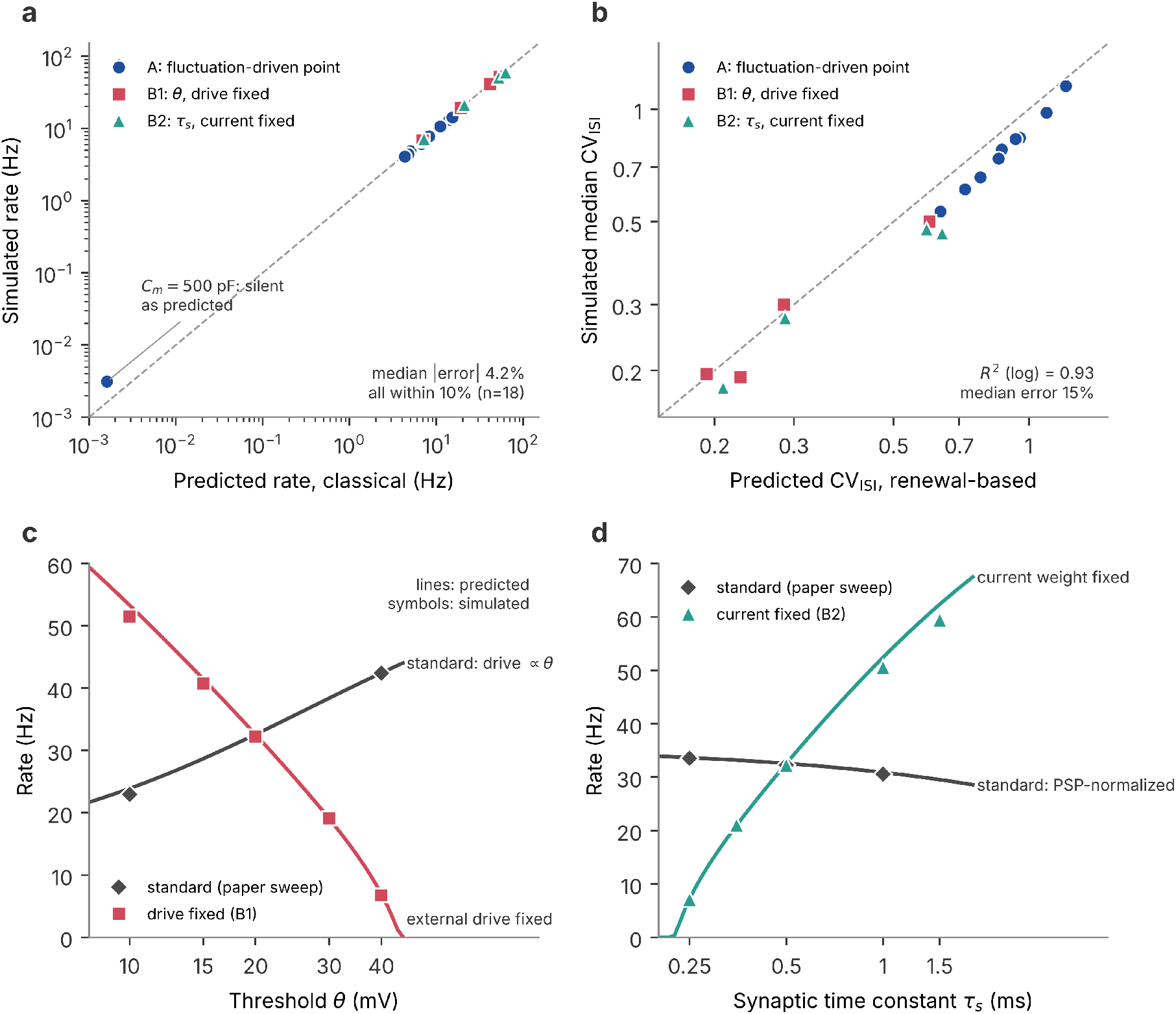
Pre-specified generality test. Nineteen conditions, three seeds each, with all predictions recorded before the simulations. Family A: fluctuation-driven operating point (*J* = 0.3 mV, *γ* = 6, *η* = 1.5) with one-factor variations under the standard calibration. Family B1: *θ* varied with the external drive held fixed. Family B2: *τ*_*s*_ varied with the synaptic current weight held fixed. Families are distinguished by color and symbol. **(a)** Simulated rate (mean *±* SD of three seeds) against the classical prediction; the silent *C*_*m*_ = 500 pF condition is plotted at its rate. **(b)** Simulated median CV_ISI_ against the renewal-based prediction. **(c)** Rate against threshold under the standard calibration (diamonds, simulated sweep of the main study) and with the external drive held fixed (squares); lines are the predictions. **(d)** Rate against synaptic time constant under the standard PSP-normalized calibration (diamonds) and with the current weight held fixed (triangles). The two conventions coincide at the reference point (*θ* = 20 mV, *τ*_*s*_ = 0.5 ms), where the curves cross. Dashed lines in a and b, identity

The alternative conventions tested the central claim most directly, because they change what a nominal parameter means without changing the network. With the external drive held fixed (B1), raising the threshold from 10 to 40 mV lowered the rate from 51.5 to 6.8 Hz (predicted 53.3 to 7.0 Hz), whereas under the standard calibration the same change raised it from 23.0 to 42.4 Hz (Fig. 11c). With the current weight held fixed (B2), length-ening *τ*_*s*_ from 0.25 to 1.5 ms raised the rate from 7.1 to 59.4 Hz (predicted 7.2 to 62.4 Hz), whereas under PSP normalization *τ*_*s*_ had almost no effect (33.5 to 30.6 Hz between 0.25 and 1 ms; Fig. 11d). In both cases the sign of the parameter’s effect was set by the convention, and the effective variables predicted it before the simulations were run.

The renewal-based theory predicted CV_ISI_ with *R*^2^ = 0.93 on log CV and a median error of 15%, the same accuracy as in the global design, and again overestimated the irregularity of the fluctuation-driven conditions (by 5–19%). The classical stability analysis predicted no instability in any condition, and all 57 simulations were asynchronous or quiescent (*S*_cons_ ≤ 0.025). As in the sweeps, the renewal-based analysis predicted weak instabilities in three mean-driven conditions (B1 at *θ* = 10 and 15 mV, and B2 at *τ*_*s*_ = 1 ms) that were asynchronous regular in the simulations. The predicted state category matched the observed one in 14 of 19 conditions; the five mismatches lay within 0.15 of a CV_ISI_ threshold of the classifier, across which the overestimate of the CV had moved the prediction.

## 4 Discussion

We examined the apparent dependence of network activity on nominal parameters by comparing the simulations with two theoretical descriptions obtained without fitting to the simulated outputs. The first describes the stationary state implied by the model calibration, while the second describes the linear stability of that state. This strategy builds directly on the classical mean field theory of sparse excitatory and inhibitory networks, in which firing activity is determined by effective input statistics rather than by nominal model parameters considered individually (Amit & Brunel, 1997; Brunel, 2000; Fourcaud & Brunel, 2002). Our purpose was not to introduce a new stationary theory, but to ask whether this established theoretical structure can explain the apparent effects that emerge when the parameters of a commonly used spiking network implementation are varied one at a time or jointly over a broad domain.

The results show that much of this apparent parameter dependence can indeed be reduced to a small set of effective variables. Seven of the eight nominal parameters influence the stationary state through the external drive relative to threshold, the net recurrent coupling relative to threshold, the input noise ratio, and ratios between the relevant time scales. When evaluated using the calibration associated with each simulated condition, stationary Siegert theory predicted firing rates across 78 sweep conditions with a median error of approximately 1% and reproduced the hierarchy of local rate sensitivities. After the recurrent variance was modified to account for the low frequency suppression associated with regular presynaptic firing, the same framework also reproduced spike train regularity over more than two orders of magnitude.

The reduction remained predictive when the parameters were varied jointly rather than one at a time. Without refitting, the stationary theory explained approximately 95% of the firing rate variance and 90% of the variance of log regularity across 10,240 independently sampled parameter combinations, while reproducing the main features of the global Sobol sensitivity structure. A pre specified generality test provided a stronger test because it moved the network to a different operating regime and changed two calibration conventions. The theory remained accurate under those conditions and predicted two reversals in the direction of nominal parameter effects before the corresponding simulations were performed. These results extend the classical mean field description by showing that it can also serve as an inferential framework for understanding what parameter sweeps actually manipulate.

The departures from stationary theory were equally informative. They were concentrated in synchronous and bursting states rather than distributed randomly across parameter space. Such departures are expected from the classical theory of recurrent spiking networks because a stationary solution can exist while being dynamically unstable to collective fluctuations (Brunel, 2000; Brunel & Hakim, 1999). When the stationary solutions were subjected to linear stability analysis, the resulting instability structure predicted the onset of collective oscillations, their dependence on synaptic delay, and their dominant frequencies. The overall picture is therefore hierarchical. Stationary firing rate and regularity are largely organized by a small set of effective variables imposed by the calibration, while the principal departures from that reduction arise when the asynchronous stationary state loses stability and collective timing becomes dynamically relevant.

### 4.1 Why the collapse is not guaranteed by construction

It could be argued that the collapse onto (*η*_eff_, Γ, *σ/θ*) follows automatically from the way these quantities were obtained from the calibration in Eqs. 1 through 7. Such an interpretation would be incomplete because the algebra guarantees only that the first two moments of the *input* can be expressed through these effective quantities. It does not guarantee that the *output* of the recurrent network, including firing rate and spike train regularity, will depend only on those moments.

This distinction is central to mean field descriptions of recurrent networks. The reduction from microscopic interactions to population input statistics requires assumptions about weak correlations, sufficiently large and sparse connectivity, and an adequate diffusion description of synaptic fluctuations (Amit & Brunel, 1997; Brunel, 2000; Fourcaud & Brunel, 2002). Even when those conditions are approximately satisfied, the stationary solution need not remain dynamically stable. Previous work has shown that recurrent networks can leave an asynchronous state through collective instabilities without requiring a large change in the stationary mean input (Brunel, 2000; Brunel & Hakim, 1999; Roxin et al., 2005). The agreement between effective variables and simulated outputs is therefore an empirical result rather than an algebraic identity.

The simulations demonstrate both sides of this distinction. Across the asynchronous conditions, firing rate and regularity were predicted with high accuracy, indicating that the nominal parameters contributed little additional explanatory information once the effective input variables were known. In synchronous and bursting conditions, however, the regularity prediction deteriorated markedly and the collapse ceased to provide a complete description of the observed dynamics. Importantly, these failures occurred where the stability analysis independently predicted that collective modes should become weakly damped or unstable. The reduction is therefore informative not only where it succeeds but also where it fails. Successful collapse identifies regions in which nominal parameter dependence can be explained through stationary input statistics, whereas failure of the collapse identifies conditions in which population dynamics introduce information that stationary statistics cannot contain.

This separation is related to a broader distinction in recurrent network theory between the properties of a stationary solution and the dynamics that develop around it. Similar mean firing rates can coexist with substantially different temporal organizations of network activity, and asynchronous states themselves can differ qualitatively in their internal dynamics (Ahmadian & Miller, 2021; Ostojic, 2014). The present results provide a particularly direct example because some parameters strongly alter temporal organization while producing little or no change in the stationary firing rate. A successful stationary reduction should therefore be interpreted as a statement about the variables that determine the mean operating point, not as a complete characterization of the dynamical state of the network.

### 4.2 Nominal parameters as aliases

The most direct consequence of the effective variable reduction is that several effects that appear distinct when plotted against nominal parameters are different routes through the same lower dimensional control variables. Connectivity, inhibitory strength, and firing threshold all modify the recurrent coupling variable Γ. A network with *ϵ* = 0.2 closely matches the rate and regularity obtained with *γ* = 6, while *θ* = 100 mV produces nearly the same firing rate as *ϵ* = 0.02 as shown in Table 4. The nominal manipulations are physically different, but under the adopted calibration they place the network at similar locations in effective input space.

This result provides a specific mechanistic interpretation of parameter degeneracy. Studies of biological neurons and circuits have repeatedly shown that similar activity patterns can arise from widely different combinations of cellular and synaptic parameters (Marder & Goaillard, 2006; Marder & Taylor, 2011; Prinz et al., 2004). Those findings established that observed activity does not uniquely determine the underlying microscopic parameter set and motivated the broader concepts of degeneracy, compensation, and robustness. The present results identify a simpler source of degeneracy that can operate even in a homogeneous model without adaptive compensation. Different nominal parameter combinations can produce similar outputs because they map mathematically onto the same effective drive and recurrent coupling. In this model family, a many to one relation between nominal parameters and activity can therefore arise directly from a many to one relation between nominal parameters and input statistics.

This interpretation is particularly important for parameter sweeps because the label attached to a parameter axis can encourage an intuitive mechanistic explanation that is not actually isolated by the manipulation. The threshold sweep provides a clear example. Under the standard calibration, increasing *θ* increases firing rate even though a higher threshold would ordinarily be interpreted as reduced neuronal excitability. The apparent paradox disappears when the calibration is considered because external drive is rescaled with threshold while the postsynaptic potential remains fixed. Raising *θ* therefore weakens recurrent coupling relative to threshold and moves Γ toward zero. Under this convention, the threshold sweep behaves primarily as a recurrent coupling manipulation rather than as an isolated test of excitability.

The capacitance and membrane time constant sweeps illustrate the same principle in different ways. Increasing *C*_*m*_ reduces the effective drive because the calibration is held at its reference capacitance, eventually moving the network below the level required for sustained activity. Changes in *τ*_*m*_ above approximately 10 ms mainly rescale the temporal unit of the system, so large changes in firing rate measured in hertz correspond to much smaller changes when activity is expressed in membrane time units. These results do not imply that capacitance or membrane time constant lack physiological meaning. They show that the consequence of varying a nominal parameter depends on what the calibration allows to change with it.

The pre specified generality test provides the strongest evidence for this interpretation because it altered the calibration while leaving the underlying neuron and network model unchanged. When external drive was no longer rescaled with threshold, increasing *θ* reduced firing rate rather than increasing it. When synaptic current weight was no longer adjusted to preserve postsynaptic potential amplitude, increasing *τ*_*s*_ became a strong excitatory manipulation rather than having only a modest effect. The effective variables predicted both reversals before the simulations were performed. The direction of a parameter effect is therefore not necessarily an intrinsic property of that parameter but can be determined by the calibration convention through which the parameter enters the effective network equations.

This result suggests a practical change in the interpretation of computational parameter studies. A nominal parameter should not be credited with an independent mechanism until it has been established that the observed effect cannot already be explained by the effective drive, coupling, noise, or temporal scaling induced by the calibration. Reporting *η*_eff_, Γ, the relevant noise ratio, and the principal time scale ratios alongside the nominal parameter values would make such dependencies visible and would allow apparently different parameter manipulations to be compared on a common theoretical basis.

The effective reduction does not imply that all nominal parameter differences are dynamically equivalent. Conditions with similar firing rates can differ substantially in regularity when *σ/θ* differs, and parameters that contribute similarly to stationary input statistics can have different effects on stability or collective timing. The effective variables therefore identify both the dimensions along which nominal parameter differences become degenerate and the additional dimensions along which that degeneracy is broken.

### 4.3 Why the classical theory overestimates irregularity

The classical diffusion approximation predicted firing rate accurately but substantially overestimated spike train irregularity at the reference condition. This discrepancy follows from a specific assumption of the classical treatment. Presynaptic spike trains are approximated as Poisson processes, which assigns recurrent fluctuations a flat spectrum with variance proportional to the firing rate. That approximation is appropriate in strongly irregular regimes but becomes inaccurate when presynaptic neurons fire regularly.

Previous theoretical work has shown that the power spectrum of a regular renewal process is strongly suppressed at frequencies below the mean firing rate and approaches its Poisson value only at higher frequencies (Dummer et al., 2014). Dummer and colleagues developed a self consistent description of this spectral structure for sparsely connected spiking networks and demonstrated that recurrent spike statistics need not be represented accurately by a Poisson approximation. The present network provides a particularly favorable setting for observing this effect because its reference state is mean driven and highly regular. The membrane corner frequency is approximately 8 Hz, substantially below the firing rate of approximately 32 Hz, so membrane integration emphasizes precisely the low frequency range in which regular spike trains contain much less fluctuation power than a Poisson process.

The renewal based correction used here is deliberately simpler than the full spectral treatment developed by Dummer and colleagues (Dummer et al., 2014). It replaces the Poisson recurrent variance with a low frequency approximation determined by the self consistent CV^2^. Despite discarding the frequency dependence of the presynaptic spectrum, this correction reduced the large overestimate of irregularity at the reference state and reproduced CV across 59 nonsynchronous sweep conditions. It also remained substantially more accurate than the classical approximation across the global parameter design.

The success of this simple correction suggests that low frequency suppression of recurrent fluctuations captures the dominant departure from Poisson statistics for regularity in the mean driven regime studied here. At the same time, the systematic errors in the global design identify the limits of this approximation. Accuracy deteriorated when synaptic filtering became slow, when weak synchrony was present, and when firing approached the refractory ceiling. These are precisely the conditions in which the temporal spectrum of the input becomes more important and cannot be represented adequately by its zero frequency limit.

The result is therefore consistent with earlier spectral theory while providing a useful practical approximation for the present model family. The renewal correction should not be interpreted as an alternative to a self consistent spectral theory. Rather, it isolates the part of that theory that matters most for the stationary regularity of these mean driven states. This distinction becomes especially important in the stability calculation because oscillatory modes occur at frequencies where the presynaptic spectrum can differ substantially from its zero frequency value.

### 4.4 Stability of the stationary state predicts the collective axis

The principal departures from stationary theory are consistent with the classical theory of collective instability in recurrent spiking networks. Brunel and Hakim showed that recurrent interactions with finite transmission delays can destabilize asynchronous firing and generate coherent population oscillations (Brunel & Hakim, 1999). Brunel subsequently placed these oscillatory regimes within a broader phase description of sparsely connected excitatory and inhibitory networks (Brunel, 2000), while Roxin and colleagues showed that transmission delays can organize a wide range of spatiotemporal dynamics in large recurrent networks (Roxin et al., 2005). The present results do not therefore identify delay induced oscillations as a new phenomenon. Their contribution is to show how this established collective instability emerges within the same parameter sensitivity framework that otherwise appears to be dominated by stationary effective variables.

Linear stability analysis of the stationary solutions predicted the onset of delay driven synchrony at 6.0 ms, which lies between the last asynchronous condition at 5 ms and the first synchronous condition at 9 ms. The same analysis predicted the membrane time constant below which synchronous locking appeared and correctly indicated the absence of classical instability along the connectivity, synaptic time constant, and capacitance sweeps. It also predicted the frequencies of the synchronous population rhythms. In mean driven synchronous states the dominant population frequency remained close to the single neuron firing rate, whereas in the delay driven states the frequency decreased as transmission delay increased. These trends are consistent with the established distinction between oscillations generated by neuronal firing periodicity and oscillations whose period is strongly constrained by recurrent delay (Brunel, 2000; Brunel & Hakim, 1999; Brunel & Wang, 2003; Roxin et al., 2005).

The global parameter design provided a substantially broader test than the individual sweeps. Across thousands of parameter combinations, the stability calculation reproduced the increase in nonstationary states with delay and predicted the dominant frequency of synchronous states with high accuracy. This result is important because the theory was not fitted to the transition boundaries observed in the simulations. Instead, each condition was evaluated from its own stationary operating point, synaptic filtering, delay, and calibration. The collective state can therefore be predicted from the stability of the same stationary solution that describes firing rate in the asynchronous regime.

The classical and renewal based stability calculations did not yield an identical transition boundary because the damping of collective modes depends on the assumed recurrent fluctuation spectrum. The classical Poisson variance assigns more low frequency power to the recurrent input and tends to stabilize the stationary state, placing the predicted transition toward the synchronous side of the observed boundary. It therefore missed some synchronous conditions but produced no false instability predictions in the one factor sweeps. The renewal based variance reduces the recurrent variance more strongly and shifts the predicted transition toward the asynchronous side. It detected every observed transition but also classified some asynchronous conditions as weakly unstable.

This difference follows naturally from the spectral approximation discussed above. Regular spike trains have strongly suppressed power near zero frequency but recover substantial power near and above their firing rate (Dummer et al., 2014). The membrane integrates the low frequency component when determining stationary irregularity, while the oscillatory instability depends on the neuronal response at considerably higher frequencies. A variance correction based only on the zero frequency spectrum can therefore describe CV accurately while overestimating the reduction in fluctuation power relevant to fast oscillatory modes. A self consistent spectral treatment combined with the frequency dependent response of the LIF neuron to filtered noise (Brunel et al., 2001; Dummer et al., 2014; Fourcaud & Brunel, 2002) should reduce the separation between the classical and renewal based stability predictions.

The two predictions should consequently be interpreted as informative limits rather than competing descriptions of the same boundary. Their separation identifies a specific theoretical uncertainty associated with the treatment of recurrent noise. The fact that the observed transitions generally fall between them supports this interpretation and provides a quantitative target for a more complete spectral theory.

Some dynamical regimes remain beyond the scope of this linear treatment. Networks with excitation dominated feedback can approach the refractory ceiling and lock into highly regular synchronous states that are not captured by the diffusion based stability approximation. At very long delays, several oscillatory modes can also become unstable simultaneously and the network can progress from coherent oscillation to bursting. Linear analysis can identify the loss of stability of the stationary solution but cannot describe the nonlinear attractor that develops after that loss. These limitations are consistent with the general role of linear stability analysis, which determines when a stationary solution becomes unstable but does not specify the full nonlinear dynamics far beyond the bifurcation.

Synaptic delay provides the clearest separation between the stationary and collective dimensions of network behavior. Delay does not enter the stationary rate equations and was correspondingly almost invisible to the stationary firing rate prediction, to the local rate elasticity, and to the global Sobol decomposition of rate. Nevertheless, changing delay reorganized the network from asynchronous regular activity to coherent synchronous oscillations and ultimately to bursting while leaving mean firing rate nearly unchanged. This separation provides a concrete example of a broader point emphasized in studies of recurrent cortical dynamics, namely that similar population averaged firing levels can coexist with qualitatively different temporal organizations of network activity (Ahmadian & Miller, 2021; Ostojic, 2014).

The global sensitivity results show that this distinction has practical consequences. Delay accounted for only a small fraction of the variance of log firing rate but was the leading total order factor for bursting and for departure from the asynchronous state in Section 3.7. A sensitivity analysis that considers firing rate alone will therefore systematically underestimate parameters whose principal influence is exerted through temporal coordination. The distinction between rate and collective timing is consequently not merely a theoretical issue but affects how parameter importance is ranked and interpreted.

### 4.5 Graph structure

The connectivity sweep altered the realized network graph substantially, including large changes in edge count, density, path length, and related graph descriptors. Strong correlations between these structural quantities and network activity could therefore be observed when conditions were compared across values of *ϵ*. Taken alone, such correlations might suggest that the changes in firing rate or regularity were caused by graph topology. The effective variable analysis shows why that interpretation is not justified in this particular ensemble.

Changing *ϵ* simultaneously changes recurrent indegree and total recurrent coupling. The stationary theory accounted for the resulting firing rate trend and much of the regularity trend through the effective recurrent coupling, while graph variation among realizations at the same value of *ϵ* added no detectable predictive information. A complementary comparison produced the same conclusion because changing *γ* to 6 left the realized graph unchanged while generating firing rate and regularity close to those obtained with *ϵ* = 0.2. The activity associated with the connectivity sweep can therefore be reproduced without the corresponding graph reorganization when effective recurrent coupling is changed by another route.

This result does not imply that topology is dynamically irrelevant. A substantial literature demonstrates that network structure can influence spiking dynamics when structural properties are varied independently of average coupling. Degree distributions can alter network activity (Roxin, 2011), while reciprocity, second order connectivity statistics, and motif structure can shape correlations and synchronization (Hu et al., 2013; Pernice et al., 2011; Trousdale et al., 2012; Zhao et al., 2011). Structured connectivity can also generate slow dynamics and high variability that are absent from homogeneous random networks (Litwin-Kumar & Doiron, 2012). The present findings are compatible with those results because the current connectivity sweep does not independently manipulate the structural features examined in those studies.

The relevant inference is therefore narrower. In a random fixed indegree ensemble with fixed synaptic weights, changing connectivity alters graph statistics and recurrent gain at the same time. Graph descriptors that covary with *ϵ* cannot consequently be assigned a causal role unless coupling is controlled independently. Genuine tests of topological mechanisms would require designs in which indegree and effective recurrent gain remain fixed while degree heterogeneity, reciprocity, community structure, or motif statistics are systematically altered. The present analysis therefore serves as a control against attributing mechanistic meaning to graph activity correlations that can already be explained by a shared dependence on mean connectivity.

### 4.6 Interpreting global sensitivity

The global sensitivity analysis provides a complementary demonstration that parameter importance is not an intrinsic property of a nominal parameter. Variance based sensitivity indices quantify how much of the variability of a chosen output is associated with a parameter over a specified input distribution, so their numerical values necessarily depend on both the sampled domain and the definition of the output (Saltelli et al., 2010, 2008; Sobol’, 1993; Sobol, 2001). The present simulations illustrate this principle directly in a recurrent neural network.

When firing rate was analyzed on its raw scale, *γ* had the largest total order contribution. The reason was not simply that inhibitory strength is universally the dominant controller of network activity. The sampled domain crossed the boundary between inhibition dominated and excitation dominated recurrent feedback, and the excitation dominated designs frequently approached the refractory ceiling. As a result, a large fraction of the raw firing rate variance reflected whether a design lay on one side or the other of this feedback boundary.

Transforming the output to log(1 + *ν*) reduced the influence of the near refractory tail and produced a substantially different ranking, in which *C*_*m*_, *γ*, and *η* made comparable contributions. Restricting the analysis to the inhibition dominated region changed the interpretation further because the first order contribution of *γ* became very small while external drive and the membrane time scale became dominant. These changes are expected from global sensitivity theory because an index measures variance over the domain supplied to the analysis rather than a context independent strength of a mechanism.

The stationary mean field theory reproduced these changes in ranking. It also reproduced the near zero first order but appreciable total order contributions of *ϵ* and *θ*. This agreement provides a mechanistic interpretation of the apparent Sobol interactions. Both parameters influence activity through Γ, and the effect of changing Γ depends on the sign and magnitude of recurrent feedback, which is strongly controlled by *γ*. The nonadditive Sobol structure can therefore arise from the nonlinear mapping between nominal parameters and effective variables without requiring an independent biological interaction between every pair of nominal parameters.

This result has two implications for the use of global sensitivity analysis in neural network models. First, sensitivity rankings should always be interpreted together with the sampled parameter distributions and the chosen output transformation. Second, mechanistic interpretation should be informed by the equations that map nominal parameters onto effective dynamical quantities. A large total order index can reflect a parameter that determines which region of effective variable space is sampled rather than a unique microscopic mechanism.

The close agreement between the simulated and theoretical sensitivity structures also suggests a practical use of the stationary theory. Because it predicts design level firing rates with *R*^2^ ≈ 0.95, the theory can provide an inexpensive surrogate for exploring parameter domains before large simulation campaigns are performed. Such a surrogate could be used to define ranges that remain within a chosen feedback regime, to identify parameter regions in which output variance will be dominated by saturation or silence, and to concentrate computational resources near conditions where the stationary approximation predicts that additional collective dynamics may become important.

### 4.7 Limitations

Several limitations define the range over which these conclusions can be interpreted quantitatively. The first concerns the diffusion approximation and the treatment of synaptic filtering. The Fourcaud and Brunel correction used in the stationary calculation was derived for exponentially filtered currents in the fast synapse limit (Fourcaud & Brunel, 2002), whereas the simulations considered alpha shaped currents and included conditions in which *τ*_*s*_*/τ*_*m*_ was no longer small. Its use over the entire parameter design should therefore be regarded as an empirical approximation rather than an exact result of the original asymptotic theory. The robustness analyses nevertheless indicate that this limitation has little effect on the principal firing rate conclusions because restricting the analysis to *τ*_*s*_*/τ*_*m*_ *<* 0.2 or removing the correction produced explained variances of 94.2% and 94.4%, respectively, compared with 94.6% in the complete design.

The second limitation concerns the treatment of recurrent spike train statistics. The renewal based variance approximation retains only the zero frequency limit of the presynaptic spectrum, while the first passage calculation of CV assumes white input noise. Previous work has shown that a complete description of recurrent fluctuations requires the full self consistent spike train spectrum (Dummer et al., 2014). Consistent with this limitation, the regularity prediction was most accurate for rapidly filtered nonsynchronous states and deteriorated when synaptic filtering became slow, weak synchrony was present, or firing approached refractory saturation. The renewal correction should therefore be viewed as a useful low frequency approximation rather than a complete theory of recurrent fluctuation statistics.

The stability calculation introduces related approximations because the single neuron response was evaluated for white input noise and because the location of the operating point depends on the recurrent variance model. Earlier work has shown that synaptic filtering can modify the frequency response of integrate and fire neurons, particularly at high frequencies (Brunel et al., 2001; Fourcaud & Brunel, 2002). The separation between the classical and renewal based instability predictions is therefore not expected to disappear within the present approximation.

Reporting the two results as a bracket makes this theoretical uncertainty explicit, while the simulated transition boundaries provide a benchmark for future treatments that combine self consistent recurrent spectra with frequency dependent neuronal response.

A further limitation is the simplicity of the network itself. The model contains current based leaky integrate and fire neurons, two homogeneous populations, random fixed indegree connectivity, and no synaptic plasticity, conductance dynamics, spatial organization, or cellular heterogeneity. This simplicity is useful because it makes the calibration and effective variables analytically transparent, but it limits direct generalization to more biologically detailed circuits. In networks with heterogeneous neuronal properties, conductance based interactions, structured connectivity, or multiple cell classes, additional effective variables may be required and some nominal parameters may no longer collapse onto the combinations identified here.

Some parameter values were intentionally extended beyond ranges that would normally be interpreted as physiologically representative. In particular, the largest values of synaptic delay, membrane time constant, capacitance, and firing threshold were included to probe the structure and limits of the effective variable reduction rather than to model specific neuronal populations. The one factor sweeps reached delays of 300 to 500 ms, membrane time constants of 200 to 400 ms, capacitances of 1000 to 2500 pF, and thresholds of 200 to 400 mV, while the global design also sampled broad ranges that included values outside those commonly associated with cortical neurons. These extreme conditions therefore should not be interpreted as estimates of biological parameter ranges. Their purpose was to stress the theoretical reduction, expose changes in dynamical regime, and determine whether the predicted mapping from nominal parameters to effective variables remained valid far from the reference state. Consequently, the global sensitivity rankings reported here describe the specified computational domain and should not be interpreted as rankings of physiological importance in biological networks.

This limitation is particularly relevant when the results are compared with biological parameter degeneracy. Biological circuits can maintain similar outputs through distributed compensation among many conductances and synaptic properties (Marder & Goaillard, 2006; Marder & Taylor, 2011; Prinz et al., 2004). The degeneracy identified here is more restricted because it follows directly from the calibration and from the statistics of a homogeneous model. The present results therefore provide one explicit mechanism capable of generating many to one parameter mappings but should not be interpreted as a complete account of degeneracy or homeostatic compensation in biological networks.

The generality test also covered only one additional operating regime and two alternative calibration conventions. Its predictive success shows that the principal interpretation is not confined to the reference state, but it does not establish quantitative universality across the many spiking network formulations currently in use. Further tests should examine fluctuation dominated regimes over broader ranges, conductance based models, heterogeneous networks, and calibrations in which external drive and synaptic efficacy are defined independently of threshold or membrane parameters.

The categorical state classifier introduces another level of approximation because its thresholds convert continuous dynamical measures into discrete labels. The observed transition intervals therefore summarize the parameter values that were sampled rather than defining exact bifurcation points. The stability analysis partly resolves this limitation by providing continuous estimates of where the asynchronous stationary solution becomes unstable, but it cannot determine the nonlinear state that emerges far beyond the instability.

Finally, the recurrence quantification and permutation entropy analyses reported in Online Resource 1 should remain descriptive. These measures detect changes in temporal organization but were not evaluated against surrogate processes that preserve firing rate and spike train regularity. They therefore cannot establish deterministic chaos or distinguish nonlinear deterministic structure from changes produced by stochastic firing statistics. This conservative interpretation is particularly important given the long history of difficulty in inferring deterministic dynamics from finite and noisy neuronal time series.

## 5 Conclusions

The present study shows that much of the apparent dependence of spiking network activity on nominal model parameters can be understood through a smaller set of effective variables imposed by the calibration. Across changes in connectivity, inhibitory strength, external drive, synaptic and membrane time constants, capacitance, and firing threshold, stationary firing rate and spike train regularity were largely determined by effective drive, recurrent coupling, input noise, and temporal scaling. These relationships were predicted without fitting by stationary mean field theory and remained accurate across a broad global parameter design and under alternative calibration conventions.

This reduction also clarifies where stationary theory ceases to be sufficient. Synaptic delay had little influence on stationary firing rate but strongly reorganized collective dynamics, producing transitions from asynchronous activity to coherent oscillations and eventually to bursting. These departures were not unexplained failures of the stationary description because their onset and dominant frequencies were largely predicted by the linear stability of the same stationary solutions. The results therefore support a hierarchical interpretation in which effective variables determine the stationary operating point, while stability determines whether that operating point persists as a dynamical state.

A major implication is that the effect associated with a nominal parameter cannot always be interpreted as a mechanism intrinsic to that parameter. The direction and magnitude of an apparent parameter effect can depend strongly on the calibration convention, and different nominal manipulations can become dynamically equivalent when they map onto the same effective variables. Parameter studies in recurrent spiking networks should therefore report effective drive, recurrent coupling, noise, and relevant time scales alongside nominal parameter values and should compare observed effects with stationary theory and stability analysis before assigning them an independent mechanistic interpretation. This framework provides a principled way to distinguish effects that arise from calibration, effects that arise from collective instability, and effects that genuinely require additional mechanisms.

## Supplementary material

Online Resource 1 (SM.pdf) contains the definition and validation of the state classifier, the numerical integration scheme, the numerical method and validation of the linear stability analysis, the graph and motif analyses of the connectivity sweep, condition-level results of the one-factor sweeps and of the generality test, recurrence and permutation-entropy analyses, the original local sensitivity summary, the seed reproducibility of the global state labels, and descriptive analyses of response regions of the global design.

## Statements and Declarations

### Ethics approval and consent

Not applicable.

### Data availability

Condition-level and design-level summary data underlying all figures and tables, the per-run metadata with the stored calibrations, and the advance predictions of the generality test are available at https://doi.org/10.5281/zenodo.23198365. The raw spike trains of the global design (about 1.0 TB) are available from the author on reasonable request.

### Code availability

All simulation, classification, mean-field, stability, and analysis code (Section 2) is openly available under the GNU General Public License, version 2 or later, at https://doi.org/10.5281/zenodo.23198365.

### Author contributions

L.G.F. conceived and designed the study, performed the simulations and analyses, and wrote the manuscript.

