## Supplemental Material for "Effective mean-field variables explain how biophysical parameters control firing rate and regularity in a spiking network"

Luciano Gonçalves Fernandes

#### Contents

**S1 Automatic classification of dynamical regimes 1**

**S2 Numerical integration 2**

**S3 Linear stability analysis: numerical method and validation 3**

**S4 Graph analysis of the connectivity sweep 4**

**S5 Validation of directed three-node motif sampling 4**

**S6 One-factor sweeps: condition-level results 7**

**S7 Generality test: condition-level results 9**

**S8 Recurrence quantification and permutation entropy 9**

**S9 Original local sensitivity summary 12**

**S10 Seed reproducibility of the global state labels 12**

**S11 Global design: response regions and between-seed variability 14**

#### S1 Automatic classification of dynamical regimes

Each realization was one statistical unit. Spike trains of the 500 excitatory and 500 inhibitory recorded neurons were combined, and all measures were computed over the 5 s analysis window. Labels were assigned by the prespecified hierarchy of Table S1, not by clustering. Quiescence and bursting were tested first. Active non-bursting realizations were then classified independently on a regularity axis and a collective axis, following the distinction between single-neuron regularity and population synchrony of Brunel (2000).

**Synchrony index.** Each spike train was binned at 1 ms and convolved with Gaussian kernels of standard deviation $\sigma_k = 1$, 2, and 5 ms. For each scale,

$$S_{\sigma_k} = \frac{\mathrm{Var}_t[N^{-1} \sum_i x_{i,\sigma_k}(t)]}{N^{-1} \sum_i \mathrm{Var}_t[x_{i,\sigma_k}(t)]}, \tag{S1}$$

and the classifier used the consensus $S_{\text{cons}} = \text{median}(S_1, S_2, S_5)$, so that synchrony had to be supported across time scales.

**Grey-zone surrogate test.** Realizations with $0.05 < S_{\text{cons}} < 0.10$ were tested against 999 surrogates in which each neuron's binned spike train (2 ms bins) was circularly shifted by an independent random offset, which preserves each neuron's spike count and temporal structure and destroys relative timing. The test statistic was the spectral concentration $C_{\text{spec}} = 1+\sum_k q_k \ln q_k / \ln K$ of the population spectrum (Welch, 1000 ms windows, 50% overlap, DC excluded; $q_k$ is the normalized power in bin $k$). A realization was called synchronous if $p_{\text{sur}} = (1 + \#\{C^{\text{sur}} \geq C^{\text{obs}}\})/1000 \leq 0.01$ and unresolved otherwise. The surrogate random stream was seeded from a fingerprint of the spike content, so labels do not depend on processing order or on duplicate runs.

**Validation.** The classifier passed all 22 tests of its validation suite: synthetic positive controls of all six states, stress tests (regular neurons with random phases, high-rate asynchronous firing, a common irregular oscillatory drive, and an intermediate broadband collective state), prespecified real-data anchors, and checks that labels do not depend on processing order or duplicate runs. Pairwise correlations, spectral coherence, peak frequency, oscillation strength, and auxiliary burst diagnostics were stored for description only and did not enter the labels.

**Table S1 Classification rules.** Applied in order; thresholds were fixed before analysis.

| Step | Criterion |
|---|---|
| Quiescent (Q) | Mean rate $\leq$ 0.1 Hz and $\geq$ 95% of recorded neurons silent. |
| Bursting (B) | Population rate (5 ms bins, 20 ms Gaussian smoothing): two-component Gaussian mixture on $\log(1 + r)$ preferred by $\Delta$BIC $\geq$ 10; high-rate bins merged across gaps $\leq$ 10 ms; $\geq$ 2 episodes of $\geq$ 20 ms; $r_{\text{high}}/r_{\text{low}} \geq 2$; burst concentration $f_{\text{spikes,burst}} - f_{\text{time,burst}} \geq 0.05$. |
| Regularity axis | Median $\text{CV}_{\text{ISI}}$ over neurons with $\geq$ 10 ISIs: regular (R) $\leq$ 0.50; irregular (I) $\geq$ 0.80; unresolved ($U_R$) in between, or if $Q_{25} < 0.5$ and $Q_{75} > 0.8$. |
| Collective axis | Asynchronous (A) if $S_{\text{cons}} \leq 0.05$; synchronous (S) if $S_{\text{cons}} \geq 0.10$; grey zone resolved by the surrogate test ($p_{\text{sur}} \leq 0.01$), otherwise unresolved ($U_S$). |
| Final label | (R,A)$\rightarrow$AR; (R,S)$\rightarrow$SR; (I,A)$\rightarrow$AI; (I,S)$\rightarrow$SI; any unresolved axis $\rightarrow$ Other active. |

### S2 Numerical integration

The membrane potential of each neuron obeys $C_m\, dV/dt = -(C_m/\tau_m)(V - E_L) + I_{\text{syn}}(t)$, where each synaptic event contributes a current with the time course of the alpha function $(e/\tau_s)\, t\, e^{-t/\tau_s}$ Rotter and Diesmann (1999). Spikes are emitted and received on the time grid (Morrison et al., 2007); the off-grid precise-spiking variants were not used. After a spike the membrane potential was clamped at the reset value for the 2 ms refractory period.

### S3 Linear stability analysis: numerical method and validation

**Linear response by threshold integration.** For white-noise input of mean $\mu(t)$ and variance $\sigma^2(t)$, the membrane-potential density $P(V,t)$ and probability flux $J(V,t)$ satisfy

$$\frac{\partial P}{\partial t} = -\frac{\partial J}{\partial V} + \nu(t - t_{\mathrm{ref}})\,\delta(V - V_r), \qquad \tau_m J = (\mu - V)P - \frac{\sigma^2}{2}\frac{\partial P}{\partial V}, \tag{S2}$$

with $P(\theta) = 0$ and $J(\theta,t) = \nu(t)$. Writing $\mu = \mu_0 + \mu_1 e^{\lambda t}$, $\sigma^2 = \sigma_0^2 + \sigma_1^2 e^{\lambda t}$, $P = P_0 + P_1 e^{\lambda t}$, $J = J_0 + J_1 e^{\lambda t}$, and $\nu = \nu_0 + \nu_1 e^{\lambda t}$ gives, away from $V_r$,

$$\frac{\partial J_1}{\partial V} = -\lambda P_1, \qquad \frac{\partial P_1}{\partial V} = \frac{2}{\sigma_0^2}\left[(\mu_0 - V)P_1 + \mu_1 P_0 - \frac{\sigma_1^2}{2}\frac{\partial P_0}{\partial V} - \tau_m J_1\right], \tag{S3}$$

and a jump $J_1(V_r^+) - J_1(V_r^-) = \nu_1 e^{-\lambda t_{\mathrm{ref}}}$. Following Richardson (2007), three solutions were integrated downward from $P_1(\theta) = 0$: one with $J_1(\theta) = 1$ and the reset jump, one driven by $\mu_1 = 1$, and one driven by $\sigma_1^2 = 1$. Requiring $J_1 \to 0$ at the lower boundary gives $R_\mu = -j_\mu/j_\nu$ and $R_{\sigma^2} = -j_{\sigma^2}/j_\nu$, where $j$ are the fluxes of the three solutions at the boundary. The density was propagated with an exponential integrator, the flux with the trapezoidal rule, on a grid of step $\sigma_0/300$ extending to $\min(V_r, \mu_0) - 7\sigma_0$. The three solutions and their common normalization were rescaled jointly whenever they exceeded $10^{100}$, which leaves the ratios unchanged and prevents overflow at high frequency. Threshold and reset carried the Fourcaud–Brunel shift of the stationary theory; because the shift is proportional to $\sigma$, the variance response included the term $-R_\mu\,(\alpha/4\sigma_0)\sqrt{\tau_s/\tau_m}$, so that the $\lambda \to 0$ limit of the loop equals the slope of the stationary closure.

**Counting unstable modes.** The loop $L(\lambda)$ of Eq. (10) of the main text is analytic for $\mathrm{Re}\,\lambda \geq 0$, because the single-neuron response has its poles in the left half-plane and $H(\lambda)$ has a double pole at $-1/\tau_s$. The number of eigenvalues with $\mathrm{Re}\,\lambda > 0$ is therefore $Z = -\Delta \arg F/\pi$, where $\Delta \arg F$ is the continuous change of the argument of $F(i\omega) = 1 - L(i\omega)$ for $\omega$ from 0 to $\infty$ (by conjugate symmetry this is half the winding over the whole imaginary axis). The delay-free part of the loop was computed on a frequency grid with spacing $\min(0.5\,\mathrm{Hz}, \nu_0/20)$ up to $\max(4\nu_0 + 20, 300)$ Hz, refined in windows around the harmonics $k\nu_0$ of the firing rate, where regularly firing neurons have resonances of width about $2\pi^2 k^2 \mathrm{CV}^2 \nu_0$, and extended logarithmically beyond the synaptic corner frequency. The delay factor was then applied on a sub-grid fine enough that $\omega\delta$ changed by less than 0.05 rad wherever $|L| > 0.8$. Beyond the end of the grid $|L| < 1$, so $F$ stays in the right half-plane and adds no winding. Real-axis crossings of $L(i\omega)$ to the right of 1 were refined by Newton iteration in the complex plane, and the fastest-growing root is reported.

**Validation.** At $\lambda \to 0$ the threshold-integration rate agreed with the Siegert formula to within 0.3%, and $R_\mu$ and $R_{\sigma^2}$ agreed with finite-difference derivatives of the Siegert formula to within 0.5%. Near the firing-rate resonance the response changed by about 1% when the voltage step was reduced from $\sigma_0/300$ to $\sigma_0/800$. Recomputing 300 global designs on a uniformly fine frequency grid left every stability label unchanged. For the delta-synapse network of Brunel (2000) ($C_E = 1000$, $C_I = 250$, $J = 0.1$ mV, $V_r = 10$ mV, $\delta = 1.5$ ms), the method reproduced the four example states of that study: $g = 5$, $\nu_{\mathrm{ext}}/\nu_{\mathrm{thr}} = 2$ was stable; $g = 6$, $\nu_{\mathrm{ext}}/\nu_{\mathrm{thr}} = 4$ was unstable to a fast oscillation at 190 Hz; $g = 4.5$, $\nu_{\mathrm{ext}}/\nu_{\mathrm{thr}} = 0.9$ was unstable to a slow oscillation at 29 Hz; and the excitation-dominated state $g = 3$, $\nu_{\mathrm{ext}}/\nu_{\mathrm{thr}} = 2$ was unstable. For the global design, the renewal-based operating point was obtained with a faster CV quadrature; at the 4,469 designs where both were available, it reproduced the stationary rate of the full calculation to within 0.07% and $c$ with a median relative difference of 0.1%.

#### S4 Graph analysis of the connectivity sweep

**Methods.** For every realization of the $\epsilon$ series, the recurrent connectivity used by NEST was exported and converted to a simple directed binary graph (multapses collapsed to one edge, autapses removed); the simulated multigraph itself was not altered. Unique in-, out-, and total degree, density $M/[N(N-1)]$, and the Shannon entropy of the total-degree distribution were computed exactly on all 10,000 vertices. Strongly connected components were identified exactly. Outward eccentricity, diameter, and radius were computed exactly within the largest strongly connected component by blockwise breadth-first search. When $\min_i g_i^- + \min_j g_j^+ > N_{\text{SCC}} - 2$, every non-adjacent pair has a common neighbor and the diameter is exactly 2, which was used as a shortcut for dense graphs. Outward Wasserman–Faust closeness was computed from full-graph distances for 256 source vertices per realization (stratified 80:20 E:I). Directed clustering and approximate betweenness (64 sources) were computed on 300-vertex induced subgraphs and were treated as descriptive only, because path structure is not preserved under induced sampling. Autapse and multapse statistics are given in Table S2.

For the structure–dynamics association, four graph measures (density, degree entropy, mean closeness, and mean eccentricity) were correlated with four dynamical measures (excitatory and inhibitory rate, median $\text{CV}_{\text{ISI}}$, and $S_{\text{cons}}$) across all 100 realizations (Spearman; 5,000 bootstrap resamples stratified by $\epsilon$). The analysis was repeated after mean-centering both variables within each $\epsilon$, with 9,999 within-$\epsilon$ permutations. Benjamini–Hochberg correction was applied separately to each family of 16 tests.

**Results.** The realized graph changed as expected for a random fixed-indegree graph of increasing mean degree (Fig. S1). Unique directed edges increased from $9.99 \times 10^4$ to $6.32 \times 10^7$ and density from 0.001 to 0.632. Degree entropy rose from 3.69 to 7.86 bits because more absolute degree values were occupied, while the relative dispersion of degree fell (coefficient of variation of out-degree from 31.5% to 0.77%). The exact directed diameter fell from 7 at $\epsilon = 0.001$ to 2 for every realization with $\epsilon \geq 0.05$, and the largest strongly connected component contained all vertices for $\epsilon \geq 0.002$. Mean outward closeness increased from 0.236 to 0.731. At $\epsilon = 1$, redundant multapses accounted for 36.8% of recurrent synapses, so this condition is not an all-to-all simple graph. The accompanying activity changes (Fig. S2) are those reproduced by the stationary theory through $\Gamma$ in the main text. Spectral concentration was non-monotonic, with a maximum at $\epsilon = 0.01$ ($0.175 \pm 0.006$). All 16 across-$\epsilon$ associations were strong ($|\rho| = 0.92$–$0.99$) and survived correction. None of the 16 within-$\epsilon$ associations did (largest $|\rho| = 0.13$, minimum $q = 0.89$).

#### S5 Validation of directed three-node motif sampling

Directed three-node motifs (the 13 connected triad classes; Milo et al., 2002; Wasserman & Faust, 1994) were estimated from $5 \times 10^5$ edge-conditioned samples per realization: a realized edge was drawn uniformly, a third vertex was drawn uniformly, and the induced triad was classified. Each sample was weighted by the inverse of the number of edges in its triad, $\hat{p}_m \propto n_m/k_m$, which removes the bias of entering a triad through one of its $k_m$ edges. The null was computed analytically from the fixed-indegree generator with replacement: for two candidate sources in a population of size $N_s$ sampled $C_s$ times, the joint absence probability is $(1 - 2/N_s)^{C_s}$ and the single absence probability $(1 - 1/N_s)^{C_s}$, and these were enumerated over all E/I compositions and edge configurations. Agreement was summarized as $\log_2(\hat{p}_m/p_m^{\text{null}})$ for motifs with an expected count of at least 25, and as the Jensen–Shannon divergence between the observed and null connected profiles

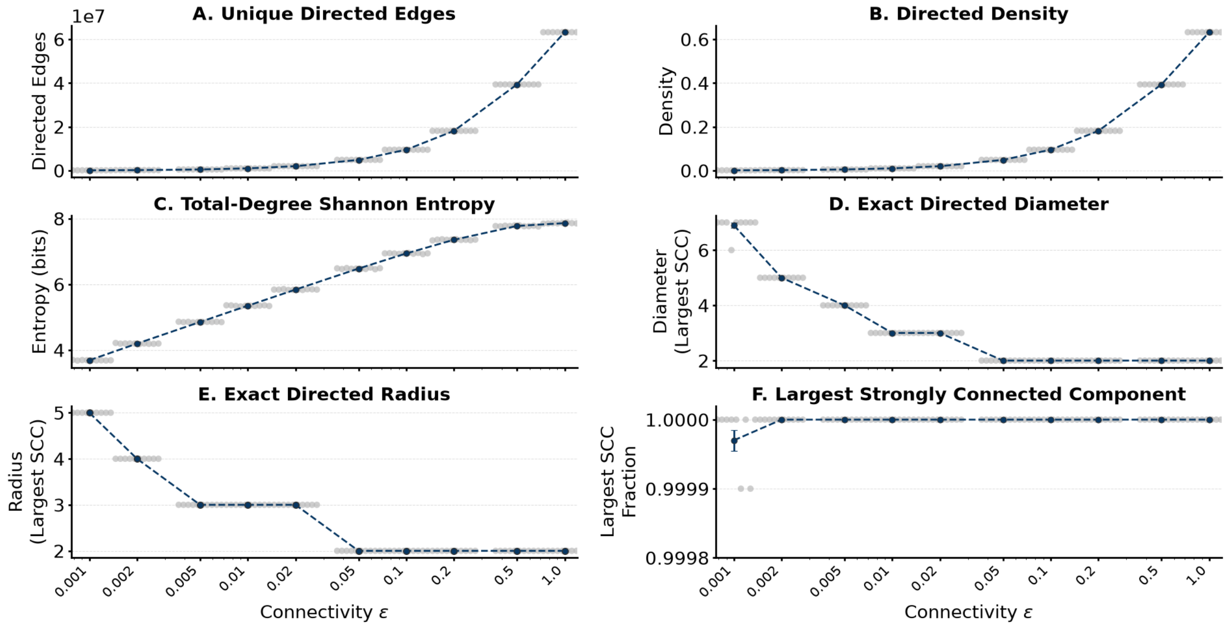


**Fig. S1 Realized directed graph across the connectivity sweep.** **(A)** Unique directed edges. **(B)** Density. **(C)** Shannon entropy of the total-degree distribution. **(D,E)** Exact directed diameter and radius of the largest strongly connected component (SCC). **(F)** Fraction of vertices in the largest SCC. Ten realizations per $\epsilon$; points are realizations, with mean ± SEM; logarithmic $\epsilon$ axis. Binary projection with multapses collapsed and autapses removed.

**Table S2** Autapse and multapse statistics across the $\epsilon$ connectivity series. Values are mean ± SEM across independent network realizations. Autapse fraction is the number of autaptic synapses divided by the total number of recurrent synapses. Multapse redundancy fraction is the number of redundant synapses beyond the first synapse for each repeated source–target pair divided by the total number of recurrent synapses. Multapses were retained in the NEST simulations but collapsed to single directed edges in the binary graph representation; autapses were retained in the simulations but excluded from the primary simple directed graph.

| $\epsilon$ | $n$ | Total recurrent synapses | Unique directed edges | Autaptic synapses | Autapses (%) | Multapse pairs | Redundant multapse synapses | Redundant fraction (%) |
|---|---|---|---|---|---|---|---|---|
| 0.001 | 10 | 100,000 ± 0 | 99,946 ± 2 | 11 ± 1 | 0.0107 ± 0.0009 | 43 ± 2 | 43 ± 2 | 0.0428 ± 0.0020 |
| 0.002 | 10 | 200,000 ± 0 | 199,801 ± 5 | 18 ± 2 | 0.0090 ± 0.0008 | 181 ± 5 | 181 ± 5 | 0.0907 ± 0.0024 |
| 0.005 | 10 | 500,000 ± 0 | 498,759 ± 11 | 48 ± 1 | 0.0097 ± 0.0002 | 1,191 ± 10 | 1,193 ± 10 | 0.238 ± 0.0021 |
| 0.01 | 10 | 1,000,000 ± 0 | 995,026 ± 18 | 105 ± 4 | 0.0105 ± 0.0004 | 4,856 ± 19 | 4,870 ± 19 | 0.487 ± 0.0019 |
| 0.02 | 10 | 2,000,000 ± 0 | 1,980,172 ± 45 | 198 ± 4 | 0.0099 ± 0.0002 | 19,509 ± 47 | 19,632 ± 46 | 0.982 ± 0.0023 |
| 0.05 | 10 | 5,000,000 ± 0 | 4,876,929 ± 91 | 500 ± 8 | 0.0100 ± 0.0002 | 120,583 ± 99 | 122,583 ± 89 | 2.45 ± 0.0018 |
| 0.1 | 10 | 10,000,000 ± 0 | 9,515,999 ± 207 | 996 ± 9 | 0.0100 ± 0.0001 | 467,368 ± 193 | 483,055 ± 205 | 4.83 ± 0.0020 |
| 0.2 | 10 | 20,000,000 ± 0 | 18,126,754 ± 192 | 2,019 ± 15 | 0.0101 ± 0.0001 | 1,751,117 ± 150 | 1,871,423 ± 198 | 9.36 ± 0.0010 |
| 0.5 | 10 | 50,000,000 ± 0 | 39,345,662 ± 994 | 4,979 ± 22 | 0.0100 ± 0.0000 | 9,018,803 ± 733 | 10,650,444 ± 980 | 21.3 ± 0.0020 |
| 1 | 10 | 100,000,000 ± 0 | 63,209,924 ± 821 | 9,995 ± 46 | 0.0100 ± 0.0000 | 26,422,865 ± 1,006 | 36,783,764 ± 821 | 36.8 ± 0.0008 |

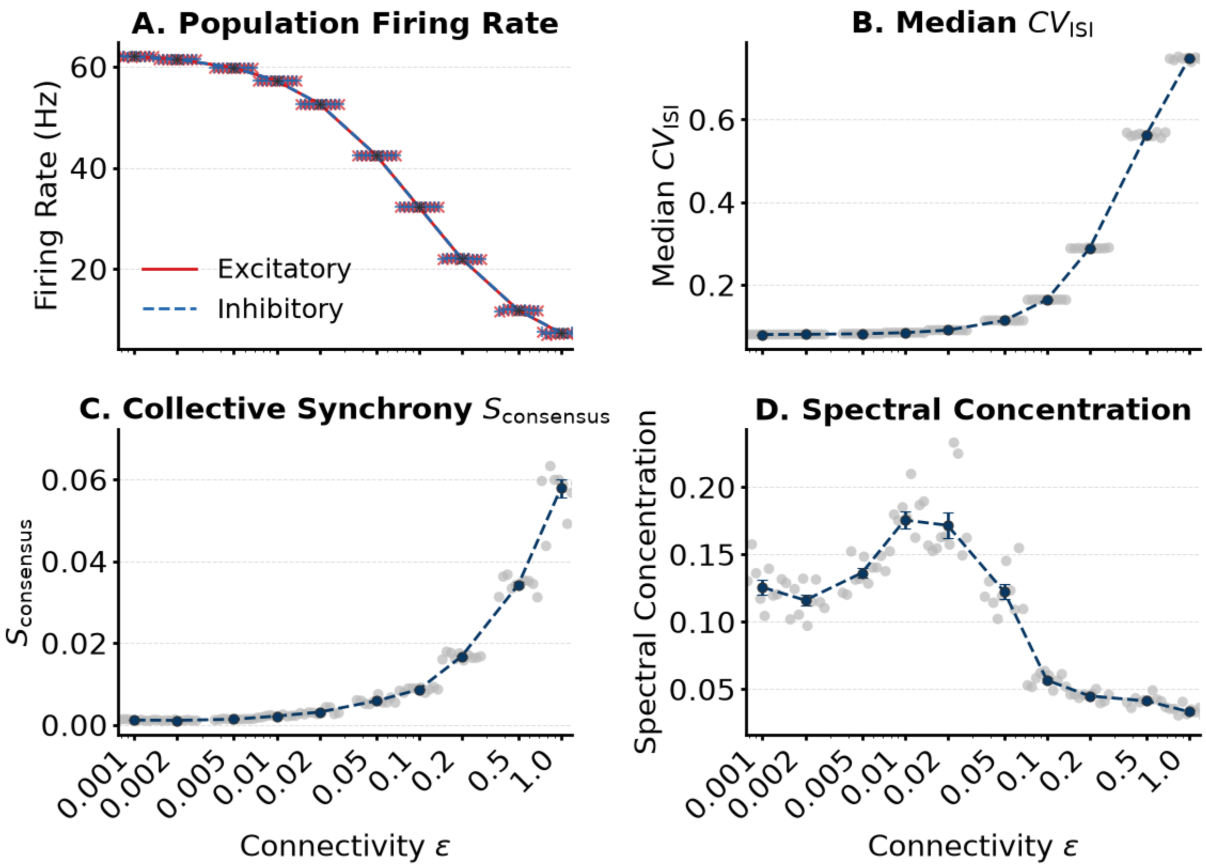


**Fig. S2 Activity across the connectivity sweep.** **(A)** Excitatory and inhibitory rates. **(B)** Median $\mathrm{CV_{ISI}}$. **(C)** $S_{\mathrm{cons}}$. **(D)** Spectral concentration. Ten realizations per $\epsilon$; mean $\pm$ SEM; logarithmic $\epsilon$ axis.

(realization as the unit; 10,000 bootstrap resamples).

Observed profiles matched the null at every $\epsilon$ (Fig. S3). Jensen–Shannon divergences ranged from $1.6 \times 10^{-6}$ to $6.8 \times 10^{-4}$, and the few individual enrichments whose intervals excluded zero were below 0.04 in magnitude (for example, 030T at $\epsilon = 0.02$, $-0.033$). Because the null reproduces the generator, this validates the sampler and the occupancy calculation; it is not a biological finding.

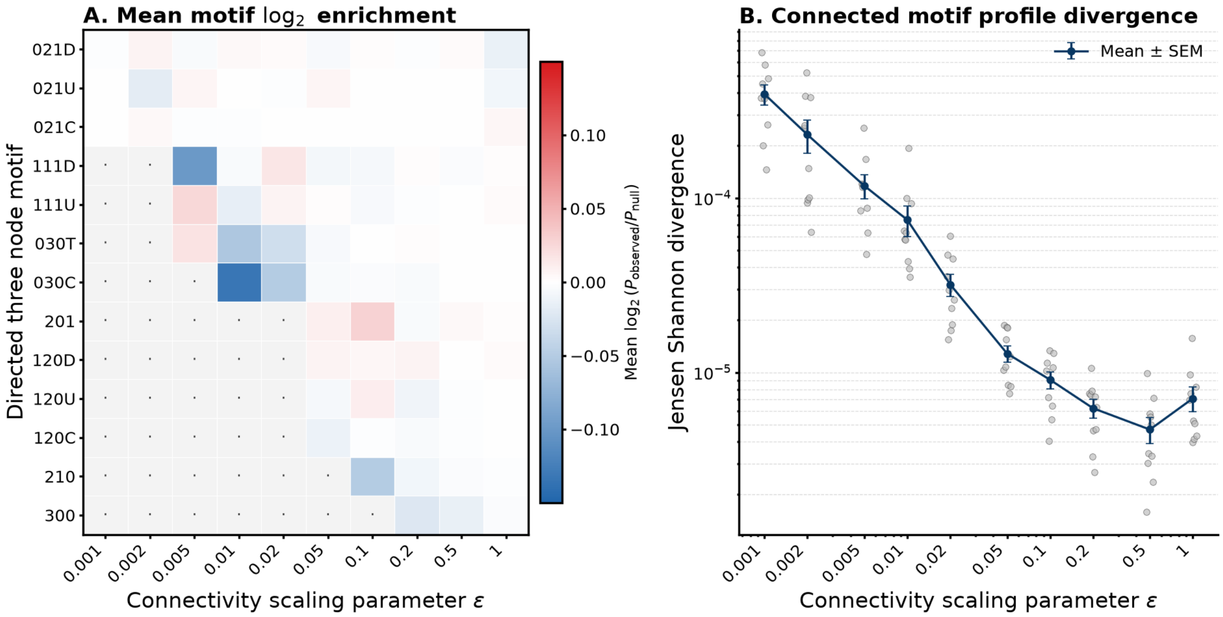


**Fig. S3 Validation of directed three-node motif sampling.** **(A)** Mean $\log_2$ enrichment of the 13 connected motifs relative to the analytical fixed-indegree null across $\epsilon$; grey cells were not estimable (expected count $< 25$). **(B)** Jensen–Shannon divergence between observed and null connected profiles for each realization (grey), with mean $\pm$ SEM (navy).

### S6 One-factor sweeps: condition-level results

Table S3 lists, for every condition of the eight sweeps, the simulated rate, regularity, synchrony, and dominant population frequency together with the stationary predictions and the linear stability labels. Figures S4 and S6 show the realization-level data for the $\gamma$, $\eta$, $\tau_m$, $C_m$, and $\theta$ sweeps, and Fig. S5 shows representative rasters for the delay and $\tau_s$ sweeps (the corresponding quantitative panels are in the main text, Fig. 6). Excitatory and inhibitory rates differed by less than 1% in every condition (at most 0.7 Hz, at $C_m = 10$ pF). Three features not discussed in the main text are worth noting. First, at $\gamma = 0$ the network fired at 390 Hz, close to the 500 Hz refractory ceiling, so its synchronous label carries little mechanistic information. Second, in the synchronous regular states produced by strong drive ($\eta \geq 6$) and short membrane time constants ($\tau_m \leq 4$ ms), the population frequency matched the single-neuron firing rate (within 1%, and within 6% at $\eta = 6$), as expected when regular neurons lock in phase. Third, several asynchronous conditions had weak spectral peaks well above the firing rate (for example 125–136 Hz for $\gamma = 6$–10 and 123–131 Hz for $C_m = 100$–125 pF); these are the weakly damped modes discussed in the main text (Section 3.8).

**Table S3 Condition-level results of the one-factor sweeps.** $\nu$: population rate (mean of 10 realizations); MF: classical stationary rate and renewal-based $CV_{ISI}$; $S_{cons}$: multiscale synchrony; $f_{pk}$: median dominant frequency of the population spectrum; states: modal labels with counts if not unanimous (Oth, Other active). Stability: linear stability of the stationary state at the classical (cl.) and renewal-based (ren.) operating points; S, stable; U, unstable, with the frequency (Hz) of the fastest-growing mode.

| Sweep | Value | $\nu$ (Hz) | MF | $CV_{ISI}$ | MF | $S_{cons}$ | $f_{pk}$ | States | cl. | ren. |
|---|---|---|---|---|---|---|---|---|---|---|
| $\epsilon$ | 0.001 | 62.0 | 61.9 | 0.0796 | 0.0828 | 0.0011 | – | AR | S | S |
| | 0.002 | 61.4 | 61.4 | 0.0800 | 0.0835 | 0.0011 | – | AR | S | S |
| | 0.005 | 59.8 | 59.7 | 0.0814 | 0.0855 | 0.0013 | – | AR | S | S |
| | 0.01 | 57.2 | 57.2 | 0.0840 | 0.0890 | 0.0021 | 57 | AR | S | S |
| | 0.02 | 52.6 | 52.8 | 0.0901 | 0.0962 | 0.0031 | 53 | AR | S | S |
| | 0.05 | 42.5 | 42.8 | 0.113 | 0.120 | 0.0058 | 43 | AR | S | S |
| | 0.1 | 32.3 | 32.6 | 0.164 | 0.172 | 0.0086 | 32 | AR | S | U (169) |
| | 0.2 | 22.0 | 22.2 | 0.288 | 0.337 | 0.017 | 94 | AR | S | U (146) |
| | 0.5 | 11.8 | 11.8 | 0.563 | 0.675 | 0.034 | 67 | Oth | S | U (105) |
| | 1 | 7.32 | 6.80 | 0.748 | 0.834 | 0.058 | 56 | Oth | S | U (92) |
| $\gamma$ | 0 | 390 | 393 | 0.0338 | 0.00373 | 0.072 | 88 | SR 8, AR 2 | U (422) | U (1574) |
| | 1 | 357 | 360 | 0.00527 | 0.00584 | 0.018 | 143 | AR | S | U (1077) |
| | 2 | 293 | 299 | 0.0116 | 0.0110 | 0.016 | 207 | AR | S | S |
| | 3 | 173 | 174 | 0.0320 | 0.0300 | 0.0076 | 173 | AR | S | S |
| | 4 | 61.8 | 62.8 | 0.0839 | 0.0863 | 0.046 | 62 | AR 7, SR 3 | S | S |
| | 5 | 32.3 | 32.6 | 0.164 | 0.172 | 0.0086 | 32 | AR | S | U (169) |
| | 6 | 21.5 | 21.7 | 0.262 | 0.304 | 0.0058 | 125 | AR | S | U (157) |
| | 7 | 16.1 | 16.3 | 0.362 | 0.439 | 0.0043 | 136 | AR | S | U (144) |
| | 8 | 12.9 | 13.1 | 0.448 | 0.542 | 0.0037 | 132 | AR | S | U (134) |
| | 9 | 10.8 | 11.0 | 0.517 | 0.616 | 0.0030 | 132 | Oth | S | U (126) |
| | 10 | 9.35 | 9.54 | 0.572 | 0.672 | 0.0027 | 130 | Oth | S | U (120) |
| $\eta$ | 1 | 6.63 | 6.99 | 0.566 | 0.679 | 0.014 | 32 | Oth | S | S |
| | 2 | 32.3 | 32.6 | 0.164 | 0.172 | 0.0086 | 32 | AR | S | U (169) |
| | 3 | 53.8 | 54.5 | 0.124 | 0.123 | 0.0097 | 54 | AR | S | U (185) |
| | 4 | 74.0 | 74.9 | 0.108 | 0.106 | 0.015 | 146 | AR | S | U (165) |
| | 5 | 93.3 | 94.3 | 0.0978 | 0.0961 | 0.027 | 185 | AR | S | U (194) |
| | 6 | 118 | 113 | 0.158 | 0.0884 | 0.36 | 125 | SR | S | U (221) |
| | 7 | 136 | 130 | 0.111 | 0.0820 | 0.65 | 137 | SR | S | U (144) |
| | 8 | 150 | 146 | 0.0893 | 0.0766 | 0.64 | 150 | SR | U (158) | U (158) |
| | 9 | 162 | 162 | 0.0783 | 0.0717 | 0.58 | 162 | SR | U (168) | U (171) |
| | 10 | 173 | 176 | 0.0692 | 0.0674 | 0.52 | 173 | SR | U (178) | U (183) |
| $\delta$ (ms) | 0.1 | 32.2 | 32.6 | 0.163 | 0.172 | 0.0064 | 32 | AR | S | S |
| | 0.15 | 32.2 | 32.6 | 0.164 | 0.172 | 0.0064 | 32 | AR | S | S |
| | 0.25 | 32.2 | 32.6 | 0.163 | 0.172 | 0.0065 | 33 | AR | S | S |
| | 0.45 | 32.2 | 32.6 | 0.164 | 0.172 | 0.0069 | 32 | AR | S | S |
| | 1.5 | 32.3 | 32.6 | 0.164 | 0.172 | 0.0086 | 32 | AR | S | U (169) |
| | 3 | 32.3 | 32.6 | 0.164 | 0.172 | 0.013 | 64 | AR | S | U (110) |
| | 5 | 32.4 | 32.6 | 0.165 | 0.172 | 0.018 | 64 | AR | S | U (75) |
| | 9 | 35.5 | 32.6 | 0.0594 | 0.172 | 0.66 | 35 | SR | U (36) | U (47) |
| | 15 | 32.6 | 32.6 | 0.184 | 0.172 | 0.42 | 30 | SR | U (26) | U (31) |
| | 18 | 32.4 | 32.6 | 0.292 | 0.172 | 0.25 | 25 | SR | U (23) | U (28) |
| | 21 | 32.5 | 32.6 | 0.337 | 0.172 | 0.18 | 21 | SR | U (20) | U (25) |
| | 27 | 33.7 | 32.6 | 0.463 | 0.172 | 0.58 | 50 | SR | S | U (53) |
| | 30 | 32.8 | 32.6 | 0.474 | 0.172 | 0.45 | 16 | SR | U (43) | U (47) |
| | 50 | 33.3 | 32.6 | 0.376 | 0.172 | 0.37 | 28 | SR 8, B 1, Oth 1 | U (28) | U (30) |
| | 100 | 32.9 | 32.6 | 0.610 | 0.172 | 0.44 | 29 | Oth 6, B 3, SR 1 | U (33) | U (34) |

| Sweep | Value | $\nu$ (Hz) | MF | $CV_{ISI}$ | MF | $S_{cons}$ | $f_{pk}$ | States | cl. | ren. |
|---|---|---|---|---|---|---|---|---|---|---|
| | 300 | 32.7 | 32.6 | 1.56 | 0.172 | – | 2 | B | U (31) | U (32) |
| | 500 | 31.3 | 32.6 | 2.29 | 0.172 | – | 1 | B | U (33) | U (31) |
| $\tau_s$ (ms) | 0.05 | 35.0 | 34.9 | 0.155 | 0.151 | 0.0092 | 36 | AR | S | U (255) |
| | 0.1 | 34.5 | 34.5 | 0.156 | 0.154 | 0.0092 | 35 | AR | S | U (241) |
| | 0.25 | 33.5 | 33.7 | 0.159 | 0.161 | 0.0086 | 34 | AR | S | U (207) |
| | 0.5 | 32.3 | 32.6 | 0.164 | 0.172 | 0.0086 | 32 | AR | S | U (169) |
| | 1 | 30.6 | 30.9 | 0.176 | 0.193 | 0.0086 | 31 | AR | S | S |
| | 2.5 | 27.2 | 27.3 | 0.228 | 0.257 | 0.0092 | 28 | AR | S | S |
| | 5 | 23.4 | 23.3 | 0.357 | 0.374 | 0.011 | 36 | AR | S | S |
| $\tau_m$ (ms) | 2 | 208 | 191 | 0.0565 | 0.144 | 0.48 | 208 | SR | U (192) | U (213) |
| | 4 | 147 | 123 | 0.0977 | 0.161 | 0.64 | 147 | SR | U (150) | U (208) |
| | 10 | 59.4 | 59.9 | 0.159 | 0.171 | 0.011 | 118 | AR | S | U (183) |
| | 20 | 32.3 | 32.6 | 0.164 | 0.172 | 0.0086 | 32 | AR | S | U (169) |
| | 40 | 17.0 | 17.1 | 0.164 | 0.172 | 0.0070 | 17 | AR | S | S |
| | 100 | 7.07 | 7.11 | 0.161 | 0.170 | 0.0058 | 7 | AR | S | S |
| | 200 | 3.59 | 3.61 | 0.158 | 0.169 | 0.0044 | 4 | AR | S | S |
| | 400 | 1.82 | 1.82 | 0.165 | 0.168 | 0.0037 | 18 | AR | S | S |
| $C_m$ (pF) | 10 | 99.5 | 106 | 3.19 | 2.96 | 0.0077 | 2 | AI | S | S |
| | 25 | 84.5 | 90.7 | 2.19 | 1.83 | 0.0085 | 180 | AI | S | S |
| | 50 | 71.1 | 76.5 | 1.06 | 1.21 | 0.013 | 150 | AI | S | S |
| | 100 | 57.0 | 59.5 | 0.390 | 0.420 | 0.016 | 123 | AR | S | U (149) |
| | 125 | 51.6 | 53.3 | 0.300 | 0.285 | 0.013 | 131 | AR | S | U (164) |
| | 250 | 32.3 | 32.6 | 0.164 | 0.172 | 0.0086 | 32 | AR | S | U (169) |
| | 500 | 6.76 | 6.96 | 0.500 | 0.601 | 0.0088 | 32 | AR 5, Oth 5 | S | S |
| | 1000 | 0 | 0 | – | – | – | – | Q | S | S |
| | 1250 | 0 | 0 | – | – | – | – | Q | S | S |
| | 2500 | 0 | 0 | – | – | – | – | Q | S | S |
| $\theta$ (mV) | 2 | 14.5 | 14.5 | 1.48 | 1.43 | 0.017 | 4 | AI | S | S |
| | 4 | 15.5 | 16.8 | 0.959 | 1.08 | 0.014 | 60 | AI | S | S |
| | 10 | 23.0 | 23.9 | 0.401 | 0.489 | 0.012 | 68 | AR | S | S |
| | 20 | 32.3 | 32.6 | 0.164 | 0.172 | 0.0086 | 32 | AR | S | U (169) |
| | 40 | 42.4 | 42.4 | 0.0764 | 0.0835 | 0.0072 | 43 | AR | S | S |
| | 100 | 53.0 | 52.8 | 0.0385 | 0.0426 | 0.0098 | 53 | AR | S | S |
| | 200 | 57.7 | 57.5 | 0.0265 | 0.0280 | 0.028 | 116 | AR 9, SR 1 | S | S |
| | 400 | 61.2 | 60.1 | 0.0190 | 0.0191 | 0.92 | 61 | SR | S | U (121) |

### S7 Generality test: condition-level results

Table S4 lists the 19 conditions of the pre-specified generality test (main text, Section 2.9) with their simulated and predicted rates, regularity, synchrony, and states. The predictions were written to `generality_predictions_preregistered.csv` on 3 October 2026, the day before the simulations, and the values recomputed afterwards from the calibration stored with each simulation were identical. The calibration of each condition is recorded in its `run_metadata.json`.

### S8 Recurrence quantification and permutation entropy

**Methods.** Recurrence quantification (Marwan et al., 2007) used the multineuron population state (per-neuron spike counts filtered with a causal 10 ms exponential kernel on a 1 ms grid, sampled

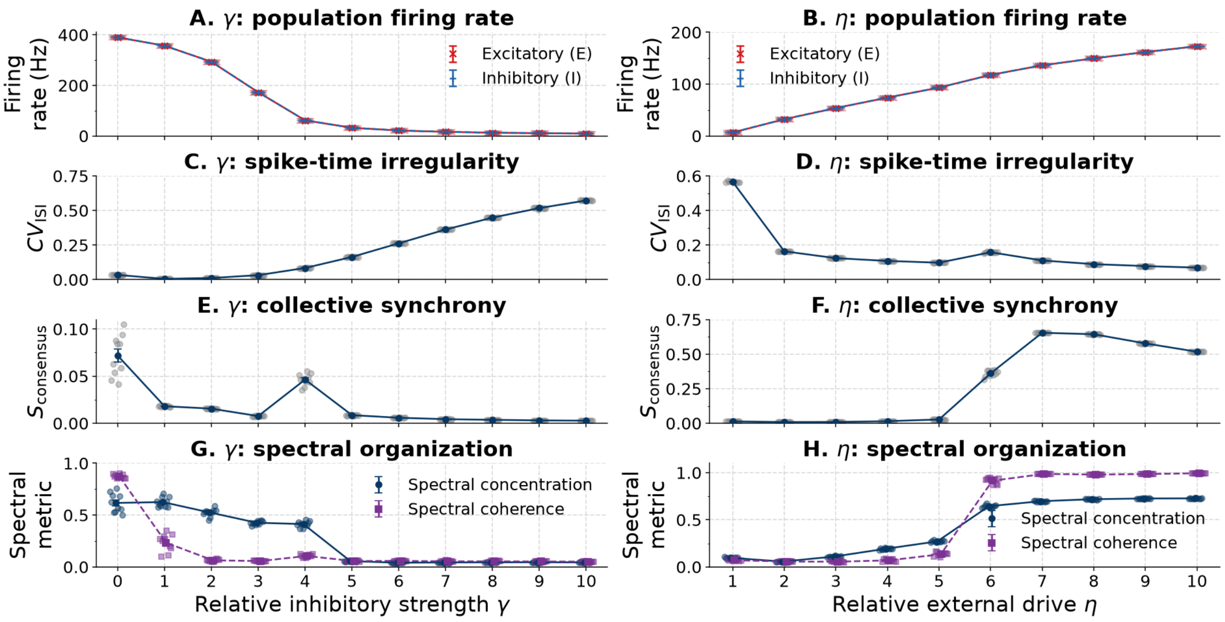


**Fig. S4 Inhibitory strength and external drive sweeps.** **(A,B)** Excitatory and inhibitory rates versus $\gamma$ and $\eta$. **(C,D)** Median $\mathrm{CV_{ISI}}$. **(E,F)** $S_{\mathrm{cons}}$. **(G,H)** Spectral concentration and coherence. Points are realizations ($n = 10$ per value), with mean $\pm$ SEM. Synchrony and spectral organization rise abruptly between $\eta = 5$ and 6.

**Table S4 Generality test: condition-level results.** $\nu$: population rate, mean $\pm$ SD over three seeds; pred.: classical prediction (rate) and renewal-based prediction ($\mathrm{CV_{ISI}}$), recorded before the simulations; states: classifier labels of the three seeds with counts if not unanimous (Oth, Other active); predicted state from the advance predictions (NS, non-stationary); stability: classical/renewal-based linear stability (S, stable; U, unstable).

| Condition | $\nu$ (Hz) | pred. | $\mathrm{CV_{ISI}}$ | pred. | $S_{\mathrm{cons}}$ | States | Predicted | Stability |
|---|---|---|---|---|---|---|---|---|
| A: base | $7.73 \pm 0.10$ | 8.22 | 0.757 | 0.863 | 0.007 | Oth | AI | S/S |
| A: $\epsilon = 0.05$ | $12.07 \pm 0.02$ | 12.85 | 0.611 | 0.721 | 0.005 | Oth | Oth | S/S |
| A: $\epsilon = 0.2$ | $4.84 \pm 0.07$ | 5.00 | 0.839 | 0.956 | 0.011 | AI | AI | S/S |
| A: $\gamma = 5$ | $13.12 \pm 0.01$ | 14.08 | 0.658 | 0.780 | 0.014 | Oth | Oth | S/S |
| A: $\gamma = 8$ | $4.40 \pm 0.01$ | 4.73 | 0.833 | 0.934 | 0.004 | AI | AI | S/S |
| A: $\theta = 10$ mV | $6.09 \pm 0.08$ | 6.70 | 0.980 | 1.10 | 0.008 | AI | AI | S/S |
| A: $\theta = 40$ mV | $10.69 \pm 0.08$ | 11.07 | 0.533 | 0.636 | 0.006 | Oth | Oth | S/S |
| A: $\tau_m = 10$ ms | $14.20 \pm 0.11$ | 15.28 | 0.780 | 0.871 | 0.011 | Oth | AI | S/S |
| A: $\tau_m = 40$ ms | $4.07 \pm 0.07$ | 4.31 | 0.738 | 0.857 | 0.006 | Oth | AI | S/S |
| A: $C_m = 125$ pF | $12.15 \pm 0.19$ | 12.73 | 1.15 | 1.21 | 0.008 | AI | AI | S/S |
| A: $C_m = 500$ pF | 0.003 | 0.002 | – | 1.00 | 0.001 | Q | Q | S/S |
| B1: $\theta = 10$ mV | $51.52 \pm 0.08$ | 53.32 | 0.300 | 0.285 | 0.013 | AR | AR | S/U |
| B1: $\theta = 15$ mV | $40.70 \pm 0.05$ | 41.48 | 0.195 | 0.192 | 0.010 | AR | AR | S/U |
| B1: $\theta = 30$ mV | $19.13 \pm 0.02$ | 19.07 | 0.192 | 0.229 | 0.009 | AR | AR | S/S |
| B1: $\theta = 40$ mV | $6.76 \pm 0.04$ | 6.96 | 0.500 | 0.601 | 0.009 | Oth 2, AR 1 | Oth | S/S |
| B2: $\tau_s = 0.25$ ms | $7.11 \pm 0.03$ | 7.20 | 0.478 | 0.593 | 0.011 | AR | Oth | S/S |
| B2: $\tau_s = 0.35$ ms | $21.03 \pm 0.03$ | 20.97 | 0.180 | 0.209 | 0.008 | AR | AR | S/S |
| B2: $\tau_s = 1$ ms | $50.59 \pm 0.07$ | 52.46 | 0.276 | 0.287 | 0.015 | AR | AR | S/U |
| B2: $\tau_s = 1.5$ ms | $59.43 \pm 0.09$ | 62.39 | 0.465 | 0.641 | 0.025 | AR | Oth | S/S |

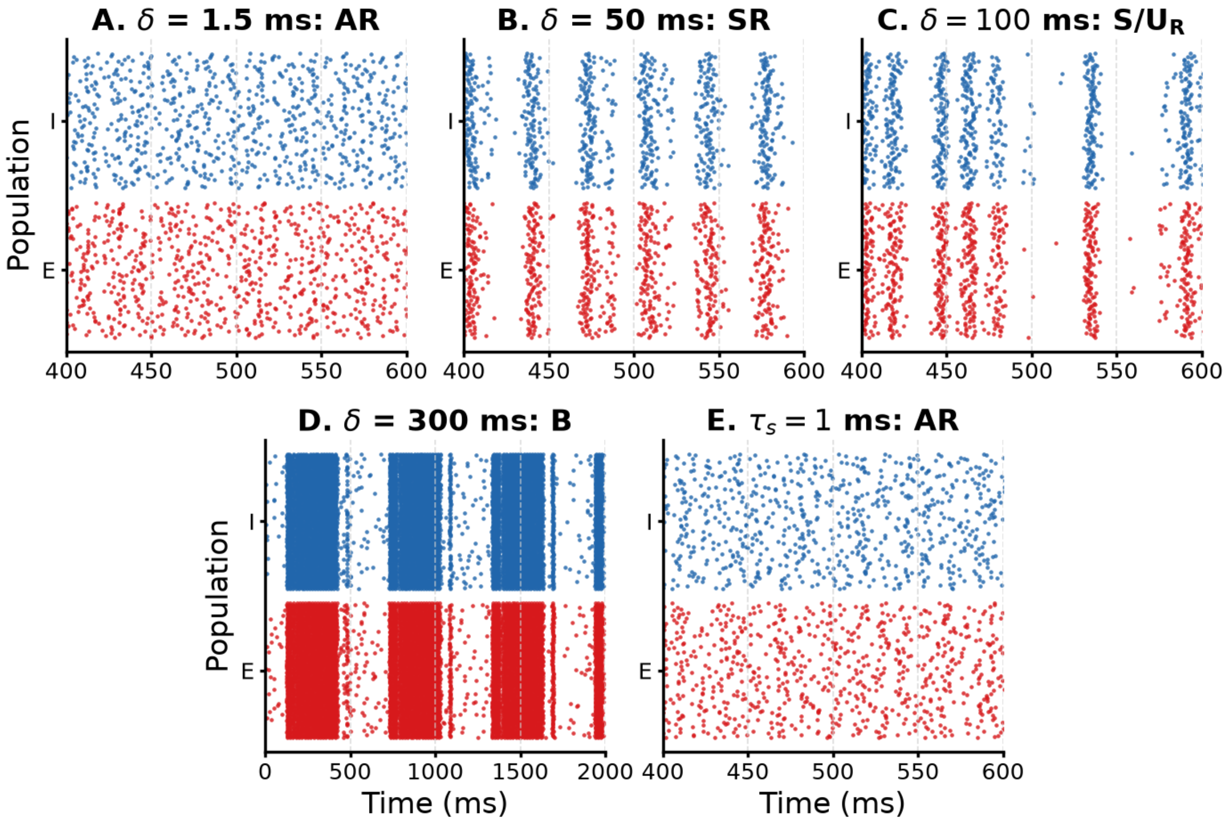


**Fig. S5 Representative rasters for the delay and synaptic time-constant sweeps.** **(A)** $\delta = 1.5$ ms, asynchronous regular. **(B)** $\delta = 50$ ms, synchronous regular. **(C)** $\delta = 100$ ms, synchronous with intermediate regularity, with episodic population activity that did not meet the bursting criteria. **(D)** $\delta = 300$ ms, bursting (longer time window). **(E)** $\tau_s = 1$ ms, asynchronous regular, representative of the whole $\tau_s$ sweep. Representative realizations were those closest to the multivariate centroid of their condition; 120 excitatory and 120 inhibitory neurons are shown. Rasters were not used for classification.

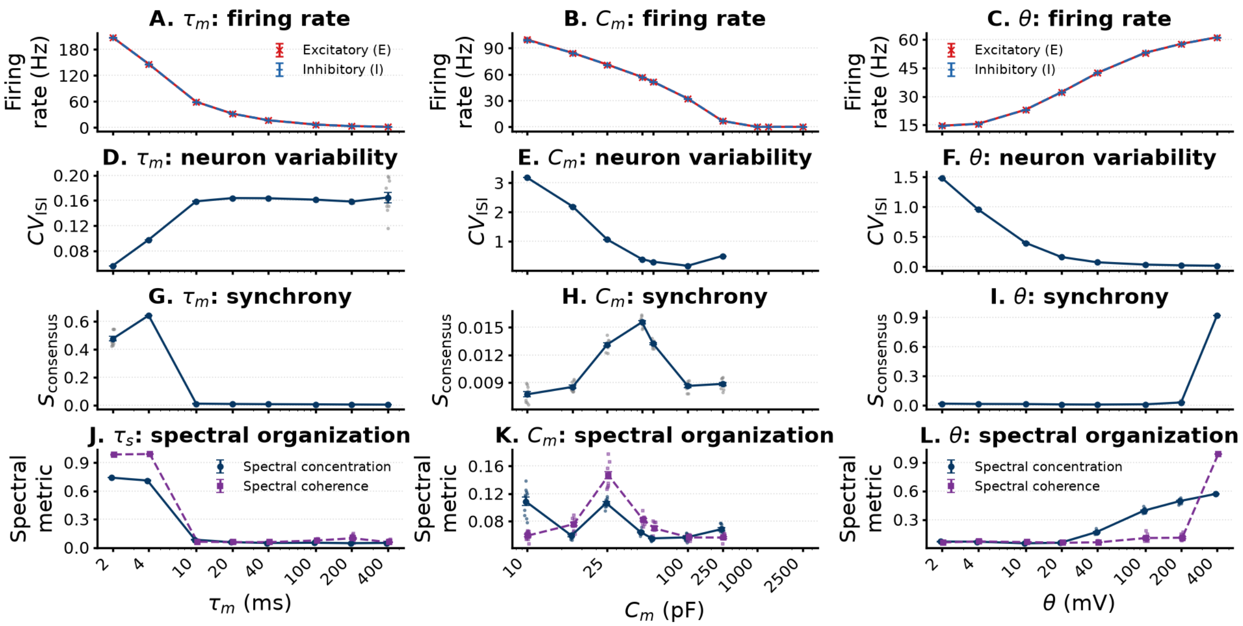


**Fig. S6 Membrane time constant, capacitance, and threshold sweeps.** Columns: $\tau_m$, $C_m$, $\theta$. **(A–C)** Excitatory (red) and inhibitory (blue) rates. **(D–F)** Median $\mathrm{CV}_{\mathrm{ISI}}$. **(G–I)** $S_{\mathrm{cons}}$. **(J–L)** Spectral concentration and coherence. Points are realizations ($n = 10$), with mean ± SEM; logarithmic axes. Measures other than rate are undefined, and not plotted, for the quiescent $C_m \geq 1000$ pF conditions.

every 10 ms after discarding five filter time constants). Euclidean recurrence thresholds were set per realization to a recurrence rate of 0.05 with a 10 ms Theiler window; determinism (DET) and laminarity (LAM) used minimum line lengths of two points. Modified permutation entropy (Bandt & Pompe, 2002; Bian et al., 2012; Riedl et al., 2013) was computed from the population spike count at 10 ms resolution (500 samples) with order $m = 4$, delay 1, and ties retained. Line measures are undefined for the constant trajectories of quiescent runs, whose mPE is zero.

**Results.** The achieved recurrence rate was 0.0500 in every active run, so DET and LAM differences reflect recurrence geometry rather than density. DET fell from about 0.6 to 0.03 across the $\epsilon$ sweep while LAM increased, and both were non-monotonic in $\gamma$, $\eta$, $\tau_m$, $C_m$, and $\theta$ (Fig. S7). Delay produced the clearest reorganization: DET dropped between 9 and 18 ms and recovered to about 0.5 at 300–500 ms, where LAM exceeded 0.7. All $\tau_s$ conditions were AR, yet DET fell from about 0.2 to 0.015, showing that these measures register temporal structure that the categorical labels do not. Because rate- and regularity-preserving surrogates could not be computed from the retained outputs and each estimate rests on 500 samples, these measures are reported descriptively only and support no conclusion in the main text.

### S9 Original local sensitivity summary

### S10 Seed reproducibility of the global state labels

All five stochastic realizations of each of the 10,240 global designs (51,200 simulations) were classified with the classifier of Section S1. The labels were highly reproducible. All five seeds agreed on the seven-state label in 95.5% of designs (Fleiss' $\kappa = 0.969$) and on the non-stationary indicator (SR, SI,

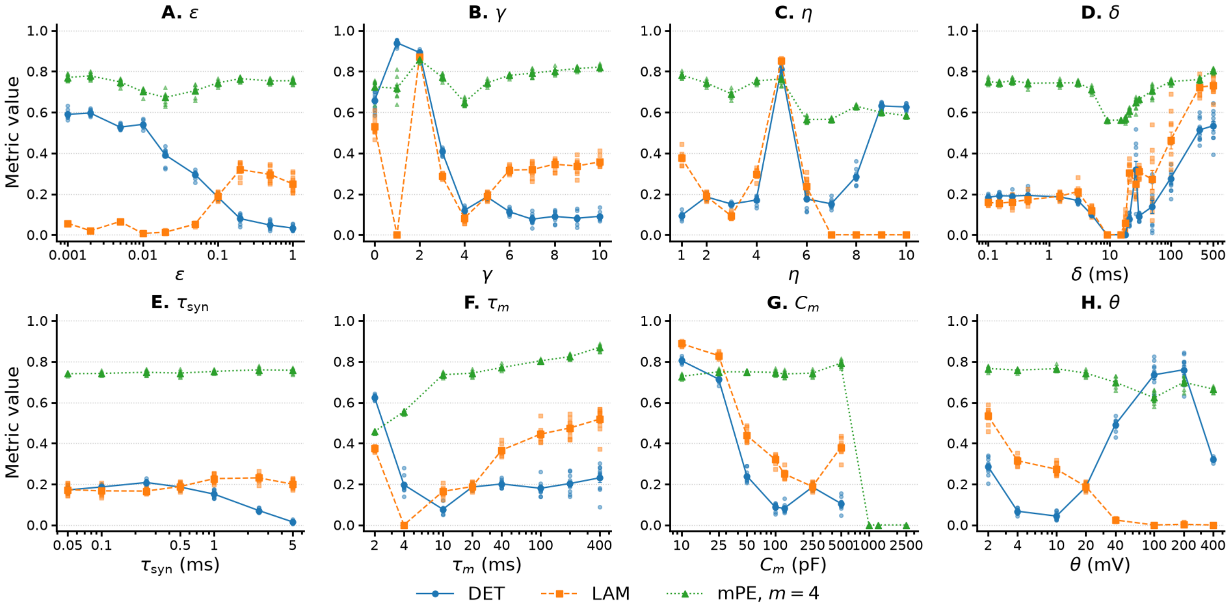


**Fig. S7 Recurrence structure and permutation entropy across the eight sweeps. (A)** $\epsilon$, **(B)** $\gamma$, **(C)** $\eta$, **(D)** $\delta$, **(E)** $\tau_s$, **(F)** $\tau_m$, **(G)** $C_m$, **(H)** $\theta$. Determinism (DET), laminarity (LAM), and normalized modified permutation entropy (mPE, $m = 4$); points are realizations, with mean ± SEM ($n = 10$). Line measures are undefined for quiescent $C_m$ conditions. These are descriptive measures and are not interpreted as evidence of deterministic chaos.

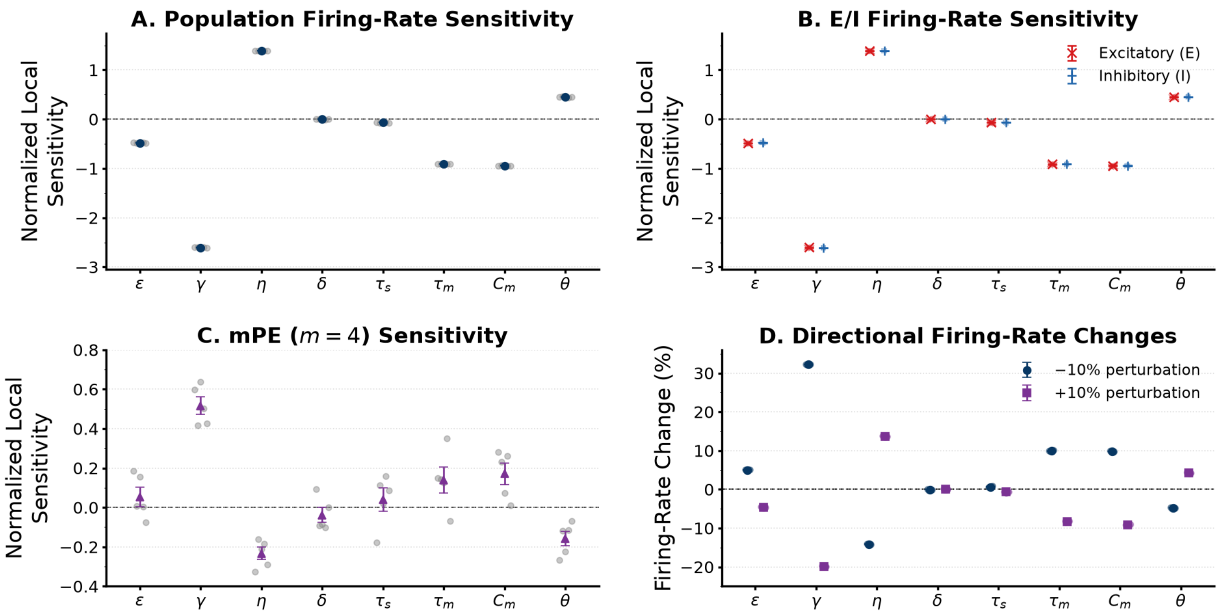


**Fig. S8 Seed-level local sensitivity of rate and mPE. (A)** Normalized centered firing-rate elasticities for each matched realization (grey) and their mean ± SEM ($n = 5$). **(B)** Excitatory and inhibitory elasticities, which differ by less than 0.005 for every parameter. **(C)** Elasticities of normalized modified permutation entropy ($m = 4$). **(D)** Directional percentage change of the population rate after $-10\%$ and $+10\%$ perturbations, showing response asymmetry (for example, $\gamma$: $+32.2\%$ and $-19.9\%$). All 85 realizations were classified AR.

or B) in 96.2% ($\kappa = 0.963$). Replicate 1 matched the modal label of its design in 98.8% of designs, and the state fractions over all simulations (AR 33.4%, AI 5.5%, SR 38.9%, SI 2.7%, B 10.1%, Q 4.6%, Other active 4.7%) differed from those of replicate 1 by at most 0.07 percentage points. Designs whose seeds disagreed on the non-stationary indicator were rare in every stability category but most frequent among designs predicted unstable only at the renewal-based operating point (7.2%), against 4.2% among designs stable at both operating points (a category that here also includes the quiescent designs) and 2.2% among designs unstable at both; the mean five-seed probability of a non-stationary state in these three categories was 0.59, 0.17, and 0.86. The total-order Sobol indices of the five-seed state probabilities reproduced those of replicate 1 to within 0.03 for every parameter and output (Table S5).

**Table S5 Total-order Sobol indices of the global dynamical state: replicate 1 versus five seeds.** Replicate 1: indicator of the state in the first realization of each design. Five seeds: probability of the state over the five realizations, with 95% whole-block bootstrap intervals. Non-stationary: SR, SI, or B; synchronous: SR or SI; bursting: B.

| | Non-stationary | | Synchronous | | Bursting | |
|---|---|---|---|---|---|---|
| Parameter | Rep. 1 | Five seeds | Rep. 1 | Five seeds | Rep. 1 | Five seeds |
| $\epsilon$ | 0.45 | 0.44 (0.39–0.49) | 0.47 | 0.47 (0.42–0.52) | 0.33 | 0.33 (0.26–0.41) |
| $\gamma$ | 0.58 | 0.57 (0.52–0.63) | 0.65 | 0.66 (0.60–0.72) | 0.58 | 0.58 (0.49–0.69) |
| $\eta$ | 0.39 | 0.38 (0.33–0.43) | 0.40 | 0.38 (0.34–0.43) | 0.21 | 0.21 (0.15–0.28) |
| $\delta$ | 0.64 | 0.63 (0.57–0.69) | 0.52 | 0.52 (0.46–0.57) | 0.69 | 0.69 (0.59–0.80) |
| $\tau_s$ | 0.30 | 0.28 (0.23–0.32) | 0.32 | 0.29 (0.25–0.34) | 0.06 | 0.05 (0.02–0.08) |
| $\tau_m$ | 0.46 | 0.44 (0.39–0.49) | 0.51 | 0.49 (0.44–0.54) | 0.27 | 0.24 (0.18–0.31) |
| $C_m$ | 0.46 | 0.45 (0.40–0.50) | 0.45 | 0.44 (0.39–0.49) | 0.29 | 0.29 (0.21–0.36) |
| $\theta$ | 0.38 | 0.37 (0.32–0.42) | 0.46 | 0.46 (0.41–0.51) | 0.40 | 0.39 (0.31–0.48) |

### S11 Global design: response regions and between-seed variability

Designs were labelled silent if both populations had zero mean rate, and high-rate, high-variability, or E/I-discrepant if, respectively, the larger population rate, the larger across-seed standard deviation, or the absolute E–I rate difference was at or above its empirical 99th percentile. Seed sensitivity was summarized by the within-design span $\Delta M = \max_r M_r - \min_r M_r$ of the five replicate mean rates $M_r = (E_r + I_r)/2$, with the top 1% defined as large. These are descriptive categories of the sampled response, not dynamical regimes.

Of the 10,240 designs, 435 (4.25%) were silent, and each upper-tail category contained 103 designs (Fig. S9C,D). Silent and high-rate designs were reproducible across seeds (median span of the high-rate designs 0.0018 Hz). High-variability designs were not: 96 of 103 were also in the top 1% of span (median span 32.5 Hz, maximum 187 Hz), which reflects between-realization heterogeneity rather than, by itself, switching or multistability. Only 12 of the 103 E/I-discrepant designs had a large span. Twenty designs mixed silent and active replicates, but every active replicate fired below 0.0007 Hz, a finite-count effect at the edge of silence rather than switching between activity states.

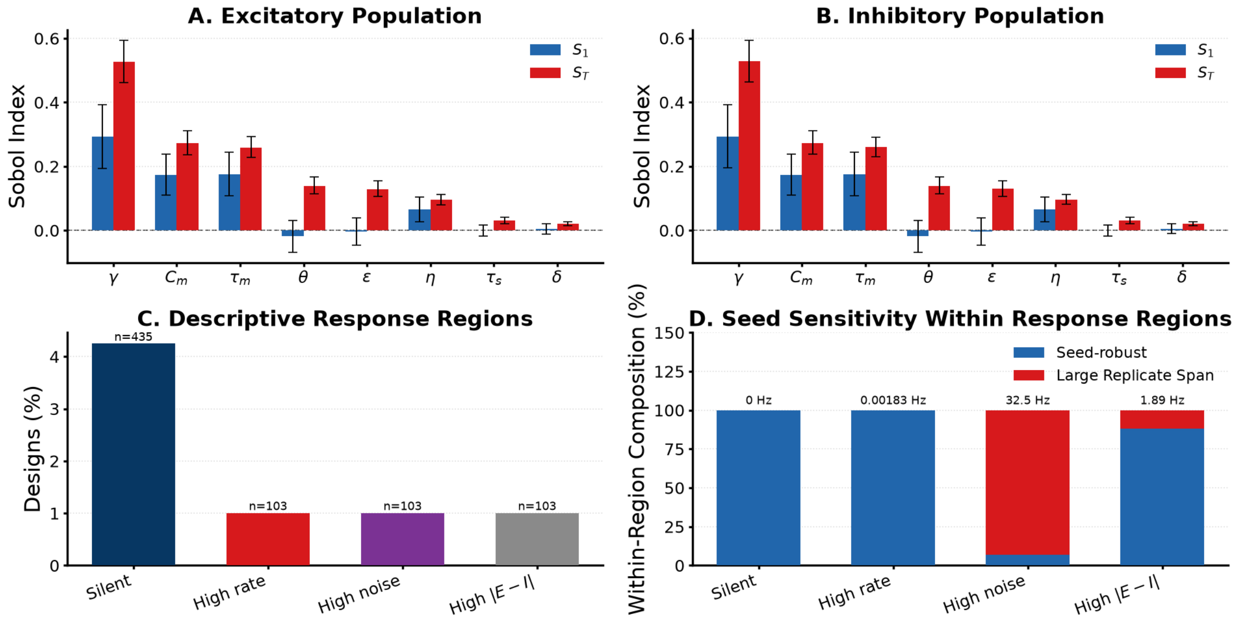


**Fig. S9 Original raw-rate Sobol analysis and response regions.** **(A,B)** First-order ($S_1$) and total-order ($S_T$) indices of excitatory and inhibitory raw firing rate, with percentile 95% intervals from 10,000 whole-block bootstrap resamples; negative $S_1$ near zero reflects estimator variability. **(C)** Fraction of the 10,240 designs in four descriptive response regions (silent; high rate, high replicate variability, and high E/I discrepancy, each defined by the empirical 99th percentile). **(D)** Fraction of each region also in the top 1% of within-design replicate span; values above bars are median spans. These regions are descriptive and are not dynamical regimes.

.88.174102

Bian, C., Qin, C., Ma, Q.D.Y., Shen, Q. (2012). Modified permutation-entropy analysis of heartbeat dynamics. *Physical Review E*, *85*(2), 021906, https://doi.org/10.1103/PhysRevE.85.021906

Brunel, N. (2000). Dynamics of sparsely connected networks of excitatory and inhibitory spiking neurons. *Journal of Computational Neuroscience*, *8*(3), 183–208, https://doi.org/10.1023/A:1008925309027

Marwan, N., Romano, M.C., Thiel, M., Kurths, J. (2007). Recurrence plots for the analysis of complex systems. *Physics Reports*, *438*(5–6), 237–329, https://doi.org/10.1016/j.physrep.2006.11.001

Milo, R., Shen-Orr, S., Itzkovitz, S., Kashtan, N., Chklovskii, D., Alon, U. (2002). Network motifs: Simple building blocks of complex networks. *Science*, *298*(5594), 824–827, https://doi.org/10.1126/science.298.5594.824

Morrison, A., Straube, S., Plesser, H.E., Diesmann, M. (2007). Exact subthreshold integration with continuous spike times in discrete-time neural network simulations. *Neural Computation*, *19*(1), 47–79, https://doi.org/10.1162/neco.2007.19.1.47

Richardson, M.J.E. (2007). Firing-rate response of linear and nonlinear integrate-and-fire neurons to modulated current-based and conductance-based synaptic drive. *Physical Review E*, *76*(2), 021919, https://doi.org/10.1103/PhysRevE.76.021919

Riedl, M., Müller, A., Wessel, N. (2013). Practical considerations of permutation entropy: A tutorial review. *The European Physical Journal Special Topics*, *222*(2), 249–262, https://doi.org/10.1140/epjst/e2013-01862-7

Rotter, S., & Diesmann, M. (1999). Exact digital simulation of time-invariant linear systems with applications to neuronal modeling. *Biological Cybernetics*, *81*(5–6), 381–402, https://doi.org/10.1007/s004220050570

Wasserman, S., & Faust, K. (1994). *Social network analysis: Methods and applications* (Vol. 8). Cambridge: Cambridge University Press.
